# R-Spondin1 regulates fate of enteric neural progenitors via differential LGR4/5/6-expression in mice and humans

**DOI:** 10.1101/2025.01.31.635899

**Authors:** Melanie Scharr, Simon Scherer, Jörg Fuchs, Bernhard Hirt, Peter H. Neckel

## Abstract

Regeneration and cytodifferentiation of various adult epithelial stem cell compartments are controlled by the Wnt-agonist R-Spondin1 (RSPO1) and the Leucin-rich-repeat-containing G-protein-coupled-receptors (LGR4/5/6). We hypothesized that RSPO1-LGR-signaling also is involved in regulating neuro-regeneration and homeostasis of the postnatal enteric nervous system (ENS). If applied to murine and human ENS-progenitors, RSPO1 led to an increased proliferation, followed by enhanced enteric neurogenesis. This coincided with an upregulation of LGR4 expression during ENS-progenitor proliferation *in vitro*. In contrast, we observed a reduced proliferation in ENS-progenitors expressing LGR5, while LGR6 was not expressed by proliferative ENS-progenitors. Instead, LGR5 and LGR6 expression increased over the course of induced neuronal differentiation, consistent with the *in vivo* expression. LGR-receptor expression therefore might represent a previously unknown mechanism influencing the fate decision of ENS-progenitor cells between proliferation and neuronal differentiation. Thus, our study is essential for our understanding of regenerative aspects of the postnatal ENS in health and disease.

## Introduction

Neurons and glial cells of the enteric nervous system (ENS) are constantly exposed to an ever-changing environment, including motility patterns^1^, microbiome^2^ or immunological interactions^3^. The ENS needs to adapt to these fluctuations to maintain its cellular homeostasis. Generally, tissue homeostasis involves self-renewal, characterized by cell proliferation, cytodifferentiation of stem and progenitors, and functional integration (reviewed in detail by Seifert et al.^4^).

In the murine ENS, neurogenesis is detectable within the first postnatal days^5,6^. Additionally, refinements of the neurochemical coding and synaptic wiring continues until adolescence^7,8^. If, however, a proliferation is preserved throughout postnatal life is controversially discussed^9^. So far, neurogenesis in the adult ENS has been convincingly described only under pathological/experimental settings like chemical denervation^10^ and colitis^11^. Nevertheless, many studies demonstrated the isolation, expansion, and differentiation of neurogenic cells from the postnatal ENS of rodents and human patients^12-14^. This suggests that under physiological conditions ENS-progenitors remain quiescent *in vivo*, arguably due to the local microenvironment. This ENS-progenitor niche remains poorly characterized^15^ and requires explorative *in vitro* studies.

In the intestinal epithelium, the interplay between cycling intestinal stem cells (ISCs)^16^, quiescent stem cells^17^, and dedifferentiating cell types^18^ enabling self-renewal and regeneration, is well-described. The hierarchical differentiation processes from ISCs to mature functional cells are tightly controlled by cell-cell-communication systems and secreted factors from the epithelium and the surrounding mesenchyme^19^. Among these factors, the Wnt-agonists R-Spondin1-4 (RSPO1-4) and Leucin-rich-repeat-containing G-protein-coupled-receptors (LGR4/5/6) are crucial to maintain the proliferative capacity of ISCs^20^. LGRs are related to glycoprotein hormone receptors, whereby LGR4/5/6 function as receptors for RSPOs^21^. RSPOs and LGR4/5/6-receptors underly dynamic expression patterns that overlap with Wnt-ligands in proliferative compartments in diverse embryonic tissues^22^. Moreover, LGR4/5 proved as useful markers to generate intestinal organoids from the mouse fetal^23^ or adult intestinal epithelium^24^. Depending on the maturation stage of the intestinal epithelium, LGR4/5-receptors serve different functions maintaining active and quiescent stem cells^25^. Whether in analogy, LGR4/5/6 play comparable roles in ENS-homeostasis is not known.

Previous works have highlighted the role of Wnt-signaling^26,27^ in postnatal ENS-progenitors from rodent models and patients *in vitro*. Moreover, we have recently shown that RSPO1-4 and LGR4/5/6-receptors are expressed on mRNA level in ENS-cells in adult mice *in vivo^28^*. Interestingly, Stavely and colleagues demonstrated that enteric mesenchymal cells (EMCs) shape the fate of postnatal ENS-progenitors by secreting morphogens including Wnt-ligands, as well as RSPO1 and RSPO3 with corresponding receptors expressed on ENS-progenitors like LGR4^29^.

The present study evaluates the cell biological function of RSPO1-LGR4/5/6 signaling in the postnatal ENS. We found that RPSO1 influences fate decision between proliferation and neuronal differentiation in murine and human ENS-progenitors, possibly due to differential expression of LGR4/5/6-receptors. Our data sets previous *in-vivo*-studies on ENS regeneration into a new context and paves the way for a mechanistic understanding of activating quiescent ENS-progenitors in the postnatal ENS.

## Results

### RSPO1-stimulation has a pro-proliferative effect on postnatal ENS-progenitors

RSPO-LGR interaction enhances active Wnt-signaling by inhibition of the Wnt-receptor-degrading ligases Rnf43 and Znrf3^30^. Hence, increasing canonical^31^ and non-canonical Wnt-signaling activity^32^ by elevating the concentration of Frizzled (FZD)-family Wnt-receptors and the co-receptors LRP5/6 on the cell-surface. We evaluated postnatal ENS-progenitors for RSPO-ligand and LGR-receptor expression using enterospheres of newborn (P0) C57BL6/J mice. After culturing for 5 days *in vitro* (5 div), we detected a robust mRNA expression of the ligands *Rspo1-4*, *Lgr4-6,* as well as *Znfr3* using RT-PCR (Supplementary Figure 1). Together with previous studies on *Fzd-* and *Lrp5/6*-receptors expression in murine enterospheres^26,28,33^, this indicates that our *in vitro* model is suitable for pharmacological probing with RSPO1.

To evaluate the cell biological function of RSPO1, ENS-progenitors from P0 C57BL6/J mice were expanded 5 days and treated with RSPO1, WNT3A alone or in combination after 1 div. Untreated cultures served as controls. After 5 div, the number (ANOVA, Fisher LSD post-hoc test, mean fold change ±SD: RSPO1: 1.53±0.25 (P=0.002); WNT3A: 1.70±0.08 (P≤0.001); Combination: 1.53±0.14 (P=0.002); n=3) and cumulative volume (ANOVA, Fisher LSD post-hoc test, mean fold change ±SD: RSPO1: 2.29±0.42 (P=0.005); WNT3A: 2.30±0.64 (P=0.005); Combination: 2.46±0.35 (P=0.003); n=3) of enterospheres in all treated groups increased compared to control (Figure 1A-1C).

**Figure 1:**
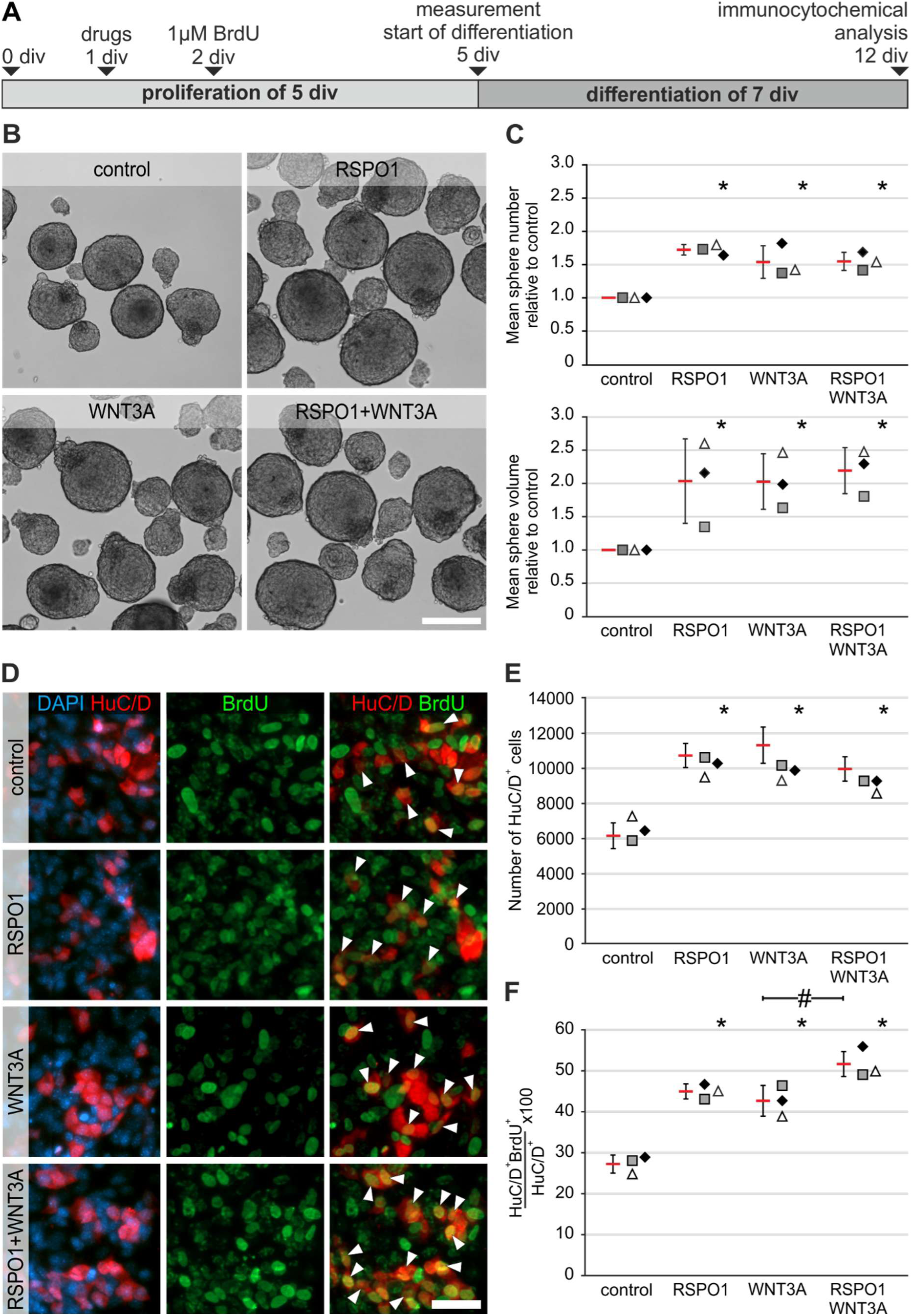
Pro-proliferative effect of RSPO1 increases enteric neurogenesis in mice. **A:** Enterospheres were cultured for 12 div. RSPO1 and/or WNT3A was applied after 1 div, BrdU after 2 div. **B:** representative enterospheres-cultures after 5 div, **Scale: 100 µm. C:** RSPO1, Wnt3A, and RSPO1+Wnt3A increased the number and cumulative volume of enterospheres compared to control. **D:** Immunostainings for BrdU and HuC/D after 12 div. BrdU^+^HuC/D^+^ neurons were detected in all groups (arrowheads). **Scale: 40 µm. E:** Number of HuC/D^+^ neurons increased after RSPO1-stimulation compared to control. **F:** RSPO1-stimulation increased the number of BrdU^+^HuC/D^+^ cells compared to control. RSPO1+WNT3A stimulation had an additional significant effect to WNT3A stimulation. Asterisks indicate significant differences compared to controls in C, E, and F (ANOVA, Fisher LSD, bars and error bars are mean±SD; n=3). See also Supplementary Figure 1-2. Source data are provided as a Source data file.

This suggests a stimulation of proliferation in ENS-progenitors by RSPO1. Therefore, we evaluated the proliferative rate of P75^+^ neural cells after RSPO1-stimulation using Ki67-stainings. The percentage of P75^+^ cells barely changed after RSPO1-stimulation compared to controls (ANOVA, Fisher LSD post-hoc test, mean±SD: Control: 33.0%±5.06%; RSPO1: 39.9%±2.31%; n=3; P=0.098; Supplementary Figure 2A-2B). However, the percentage of actively cycling Ki67^+^P75^+^ cells significantly increased after RSPO1-stimulation by 1.68-fold compared to control, suggesting that among all neural cells a subpopulation proliferates in response to RSPO1-stimulation (ANOVA, Fisher LSD post-hoc test, mean±SD: Control: 28.7 %±5.58%; RSPO1: 48.1%±7.55%; n=3; P=0.023; Supplementary Figure 2C). To analyse whether RSPO1-stimulation induces neurogenesis, stimulated enterospheres were differentiated for 7 div and stained for the pan-neuronal marker HuC/D (Figure 1D). The number of HuC/D^+^ neurons increased by 1.59-fold after RSPO1-stimulation compared to control. This effect was comparable to a WNT3A-stimulus (ANOVA, Fisher LSD post-hoc test, mean±SD: Control: 6382±733; RSPO1: 10117±1035 (P<0.001); WNT3A: 10229±688 (P<0.001), Combination: 9220±694 (P<0.002); n=3; Figure 1E).

Additionally, BrdU-incorporation revealed new-born BrdU^+^HuC/D^+^ neurons in all experimental groups suggesting that these cells derived from proliferating ENS-progenitors after isolation (Figure 1D). RSPO1-stimulation increased the percentage of newborn BrdU^+^HuC/D^+^ neurons by 1.65-fold compared to control. Further, we observed an additional significant effect of RSPO1+WNT3A (1.99-fold compared to control), (ANOVA, Fisher LSD post-hoc test, mean±SD: Control: 27.2%±2.20%; RSPO1: 45.0%±1.83% (P<0.001); WNT3A: 42.7%±3.75% (P<0.001), Combination: 51.6%±3.73% (P<0.001); n=3; Figure 1F). Therefore, RSPO1 has a pro-proliferative effect on neural cells, which is further reflected in a higher yield of newborn BrdU^+^HuC/D^+^ neurons *in vitro*.

### RSPO1-mediated effect acts directly on ENS-progenitors

To exclude additive effects by WNT-ligand secreting mesenchymal cells^29^, and to assess if RSPO1 acts directly on neural cells, we cultured FACS-purified ENS-cells using 2-month-old wnt1-tomato mice (Figure 2A). In brief, FACS-purified ENS-cells (Supplementary Figure 3), both derived from the small and large intestine, were cultured 7 div under proliferative conditions. Then, similar-sized neurospheres were transferred to a 96-well-plate containing proliferation medium, and brightfield images were taken. Afterwards, neurospheres were stimulated with RSPO1, WNT3A, or RSPO1+WNT3A, and cultured for another 7 div. Brightfield images were taken after 14 div. Untreated spheres served as control. Additionally, the single-sphere cultures exclude fusion artefacts during proliferation.

**Figure 2:**
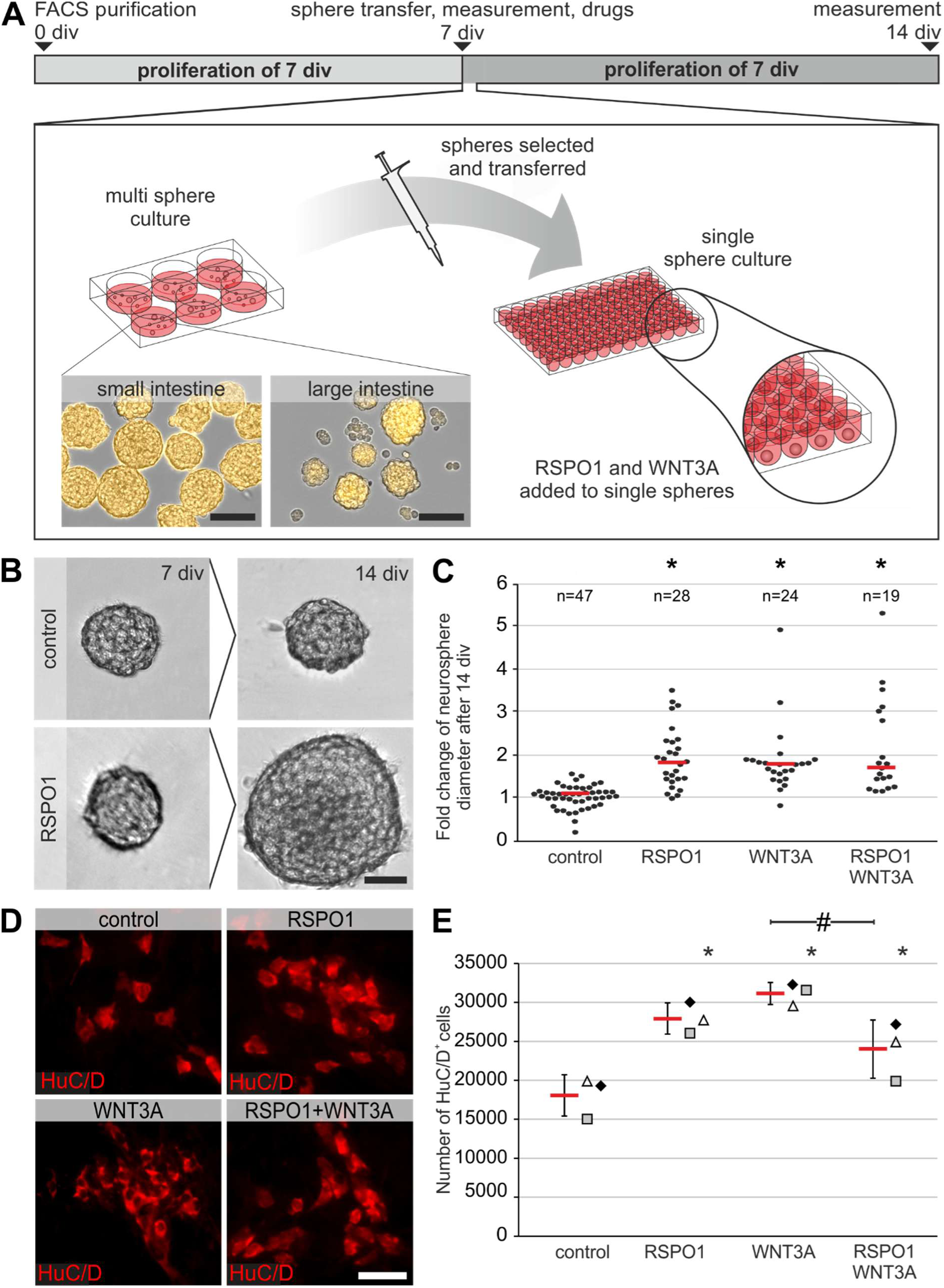
RSPO1-effect acts directly on murine ENS-cells. **A**: FACS-purified ENS-progenitors of small and large intestine were cultured for 7 div as a multisphere culture. Afterwards, equally sized neurospheres were transferred into single sphere culture until 14 div. Brightfield images were taken before RSPO1 and/or WNT3A were added and after 14 div. **Scale: 100 µm. B:** Representative images of the same neurosphere after 7 div and 14 div under control and RSPO1-treated condition. **Scale: 40 µm. C:** Dot plot depicts the fold change of neurosphere diameter after 14 div. Each dot represents one neurosphere, red bars represent the median. RSPO1- and WNT3A-stimulation significantly increased neurosphere diameter compared to control (ANOVA on Ranks, Dunn’s method, bars indicate the median fold change; four biological replicates). **D:** FACS-purified ENS-cells were proliferated as multisphere cultures until 7 div; RSPO1 and/or WNT3A was added directly after seeding. Afterwards, cells were differentiated for 7 div. Micrographs show representative images of HuC/D^+^ neurons in all experimental groups. **Scale: 50 µm. E:** Number of HuC/D^+^ neurons increased after RSPO1-stimulation compared to control (ANOVA, Fisher LSD, bars and error bars are mean±SD; n=3). Asterisks indicate significant differences compared to controls (C and E); the hash indicates significant difference between groups. See also Supplementary Figure 3-4. Source data are provided as a Source data file.

Again, we found an increase in neurosphere size in all treated groups, whereas control spheroids largely stagnated (Figure 2B-2C). Similar to WNT3A, RSPO1-stimulation significantly increased neurosphere-size (ANOVA on Ranks, post-hoc: Dunn’s method, median fold change: control: 1.12, n=47; RSPO1: 1.84, n=28, (P≤0.001); WNT3A: 1.81, n=24, (P≤0.001), RSPO1+WNT3A: 1.74, n=19, (P≤0.001), four biological replicates; Figure 2C). Comparable results were obtained from large intestinal neurospheres (ANOVA on Ranks, post-hoc: Dunn’s method, median fold change: control: 1.02, n=32; RSPO1: 1.95, n=19, (P≤0.001); WNT3A: 2.08, n=15, (P≤0.001), RSPO1+WNT3A: 1.56, n=15, (P≤0.001); four biological replicates; Supplementary Figure 4A-4B).

Further, we evaluated neurogenesis by quantifying the total amount of HuC/D^+^ neurons (Figure 2D-2E). In cultures, both derived from small and large intestine, RSPO1-stimulation significantly increased the number of HuC/D^+^ cells compared to control control (small intestine: ANOVA, Fisher LSD post-hoc test, mean±SD: control: 18072±2649; RSPO1: 27933±1995 (P=0.002); WNT3A: 31131±1412 (P<0.001), RSPO1+WNT3A: 23996±3731 (P=0.023); large intestine: ANOVA, Fisher LSD post-hoc test, mean±SD: control: 7246±1564; RSPO1: 10844±1375 (P=0.007); WNT3A: 10543±1126 (P<0.012), RSPO1+WNT3A: 10319±735 (P=0.016); n=3). Interestingly, the combinatory treatment clearly showed less effect on FACS-purified ENS-cells from small intestine compared to large intestine (Supplementary Figure 4C-4D). Overall, the described RSPO1 effect (1) does not depend on WNT-secreting mesenchymal cells but could be further augmented. (2) RSPO1 acts directly on postnatal ENS-progenitors, and (3) this effect is reproduceable on ENS-progenitors isolated from later postnatal stages.

### RSPO1 stimulates proliferation and neurogenesis in human ENS-progenitors

To assess the clinical relevance of our findings, we evaluated the RSPO1 effect on ENS-progenitors from four individual patients. Human enterospheres proliferated for 14 div; drugs were added directly after seeding. BrdU was added after 5 div and 10 div. After cell expansion, cells were differentiated for 1 week before HuC/D- and BrdU-immunolabeling (Figure 3A and 3F). After 14 div, the number (ANOVA on Ranks, post-hoc: Student-Newman-Keuls Method, median fold change (red bar): RSPO1: 2.09 (P=0.028); WNT3A: 2.27 (P=0.028); RSPO1+WNT3A: 2.16 (P=0.028); n=4; Figure 3B) and volume (ANOVA on Ranks, post-hoc: Student-Newman-Keuls Method, median fold change (red bar): RSPO1: 1.64 (P=0.023); WNT3A: 1.71 (P=0.023); RSPO1+WNT3A: 1.81 (P=0.023); n=4; Figure 3C) of human enterospheres significantly increased in all treated groups compared to control. Also, we found BrdU^+^HuC/D^+^ neurons in all groups, indicating proliferation *in vitro* (Figure 3D).

**Figure 3:**
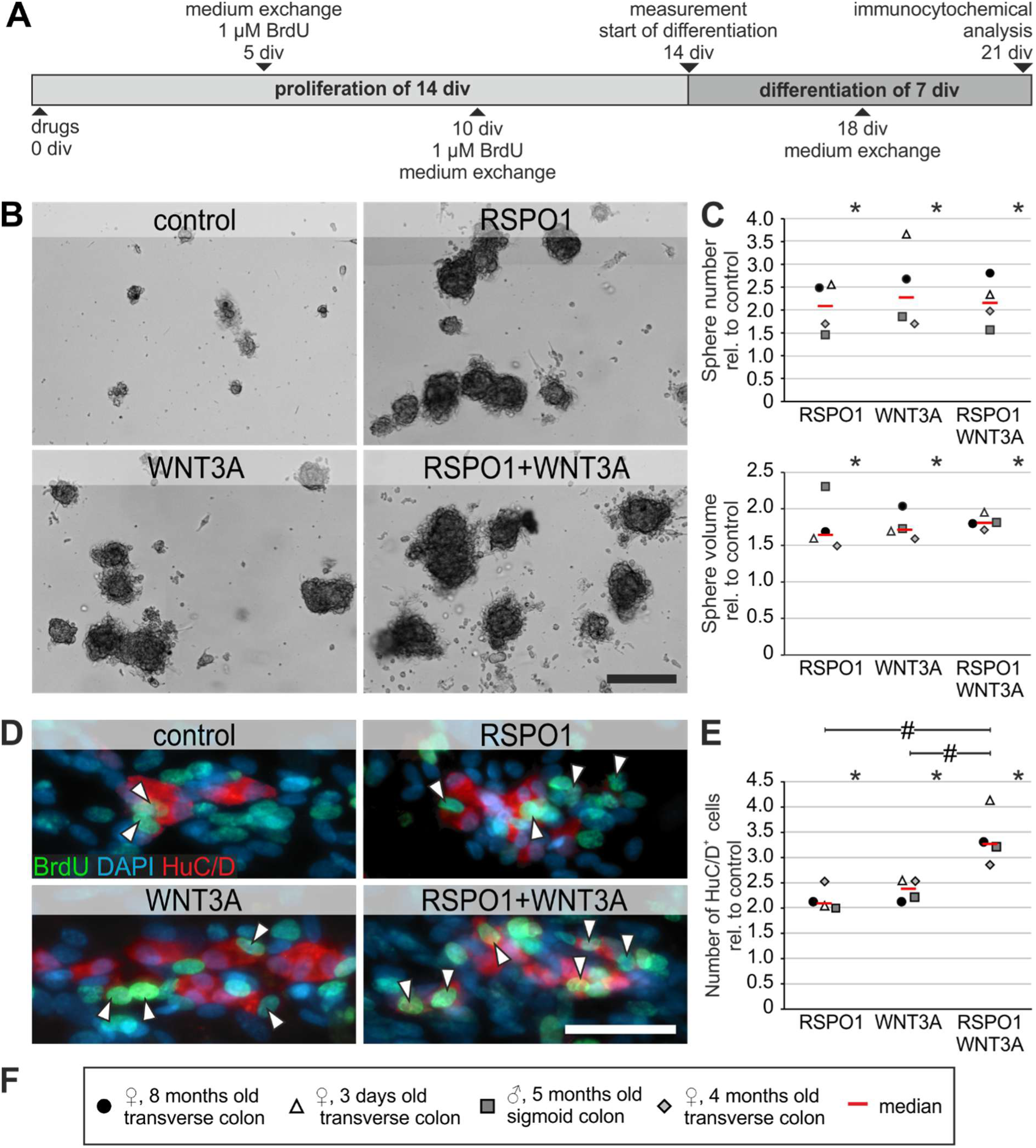
Neurogenesis is increased in human ENS-progenitors by RSPO1. **A:** human enterospheres were cultured for 21 div. RSPO1 and/or WNT3A were added directly after seeding. BrdU was added after 5 and 10 div. **B:** Representative human enteropsheres after 14 div. **Scale: 200 µm**. **C:** After 14 div, RSPO1 and/or WNT3A led to an increase in the number and cumulative volume of enterospheres relative to control. Data points for different patients are represented by different symbols. **D:** BrdU-incorporation showed BrdU^+^HuC/D^+^ neurons in all groups (arrowheads). **Scale: 50 µm. E:** The number of HuC/D^+^ neurons significantly increased after RSPO1-stimulation compared to control. Notable is the significant additional effect of RSPO1 with WNT3A. Asterisks indicate significant differences compared to controls (C and E); the hash indicates significant difference between groups (ANOVA on Ranks, Student-Newman-Keuls, median fold change (red bar); n=4). **F:** Patient data of gut specimens used. Source data are provided as a Source data file.

Moreover, RSPO1-stimulation increased the median number of HuC/D^+^ cells compared to control stimulationIn contrast to murine ENS-progenitors, the combinatory treatment significantly increased the amount of HuC/D^+^ cells by 3.27-fold (ANOVA on Ranks, post-hoc: Student-Newman-Keuls Method, median fold change: RSPO1: 2.09 (P=0.004); WNT3A: 2.38 (P=0.004); RSPO1+WNT3A: 3.27 (P=0.004); n=4; Figure 3E). Thus, the RSPO1 effect was reproducible on human ENS-progenitors suggesting a conserved signaling mechanisms among species.

### LGR5- and LGR6-receptors are expressed mainly on differentiated enteric neurons and less in glial cells in the human gut

We then analysed the expression of LGR4/5/6-receptors in the human small and large intestine with immunhistochemistry. While we detected LGR4/5/6-immunoreactivity in the ENS and other tissues of the gut wall (Figure 4 and Supplementary Figure 5-8, Supplementary Table 2), we did not observe region-specific differences between small and large intestine. Our stainings confirmed LGR4/5/6-expression in smooth muscle cells in the *Tunica muscularis* and arterioles within the intestinal wall (Figure 4, Supplementary Figure 5A for negative controls and 5B1-B3). Moreover, LGR4 was expressed throughout the crypt region up to the villus border and slightly in the *Lamina muscularis mucosae* (Supplementary Figure 5C). Further, we found a slight LGR4-immunoreactivity in PGP9.5-labeled neurites surrounding the mucosal crypts (Supplementary Figure 5C). LGR5-expression was restricted to epithelial cells located at the crypt bottom (Supplementary Figure 5D). LGR6 was expressed throughout the *Tunica mucosa* predominantly in the *Lamina muscularis mucosae* (Supplementary Figure 5E). Additionally, we detected LGR6 in PGP9.5-expressing neurites outside the ganglia within the *Tunica muscularis* and in the neurites surrounding mucosal crypts (Supplementary Figure 5E).

**Figure 4:**
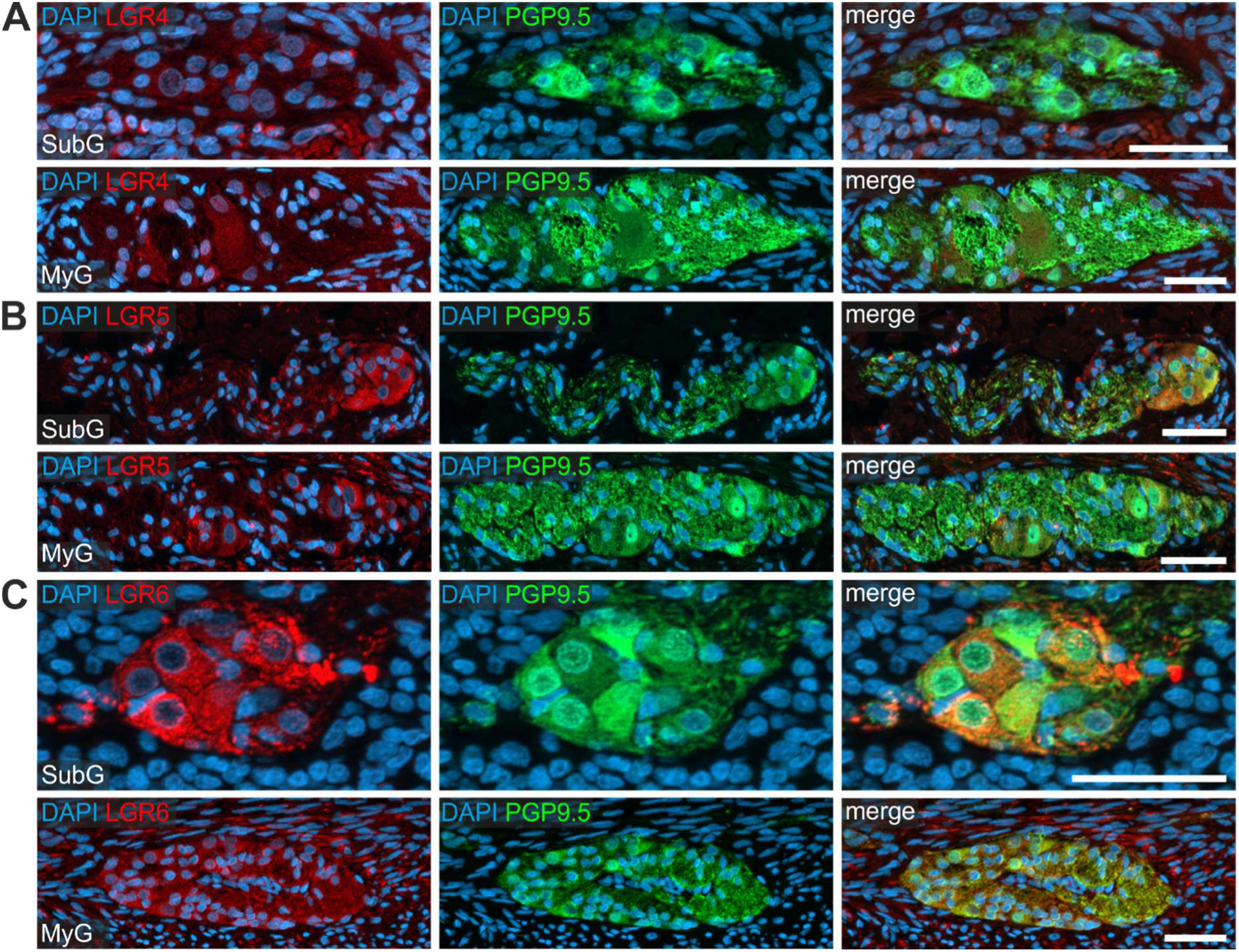
Human enteric neurons express the RSPO1-receptors LGR5 and LGR6. Immunostainings of small intestinal resectates of pediatric patients for PGP9.5 and **A:** LGR4, **B:** LGR5, **C:** LGR6. LGR5- and LGR6-expression was predominantly found in enteric neurons of submucosal (SubG) and myenteric (MyG) ganglia, whereas little to no immunoreactivity was observed for LGR4. **Scale: 50 µm.** See also Supplementary Figure 5-8.

In the ENS we found a weak homogeneous cytoplasmic LGR4-immunoreactivity in PGP9.5-labeled enteric neurons (Figure 4A) and in S100beta-labeled glial cells (Supplementary Figure 6A) of submucosal and myenteric ganglia (Figure 4A) of the small and large intestine (Supplementary Figure 7A and 8A). LGR5 and LGR6 showed a bright cytoplasmic immunoreactivity predominantly in PGP9.5-labeled enteric neurons (Figure 4B-4C, Supplementary Figure 7B-7C), yet less in S100beta-expressing glial cells (Supplementary Figure 6B and 6C). Furthermore, we detected LGR5- and LGR6-expression as fluorescent punctae in the neuropil (Figure 4, Supplementary Figure 6-8, Supplementary Figure 5 for negative controls). In summary, LGR5/6 is particularly expressed in human enteric neurons, whereas LGR4-immunoreactivity was weak, but evenly distributed throughout enteric ganglia.

### FACS-purified LGR5 negative ENS-cells gave rise to newborn neurons and glial cells

Considering the pro-proliferative RSPO1-effect on ENS-progenitors, LGR-expressing cells might display their neuro/gliogenic potential only under a highly enriched *in vitro* microenvironment. Therefore, we established a FACS-protocol for isolating LGR5-expressing cells from the human *Tunica muscularis*. We co-stained with the Wnt-receptor FRIZZLED4 (FZD4) as a useful marker for enteric neural cells from pediatric gut samples^34^ (Figure 5A-B, Supplementary Figure 9A-9D).

**Figure 5:**
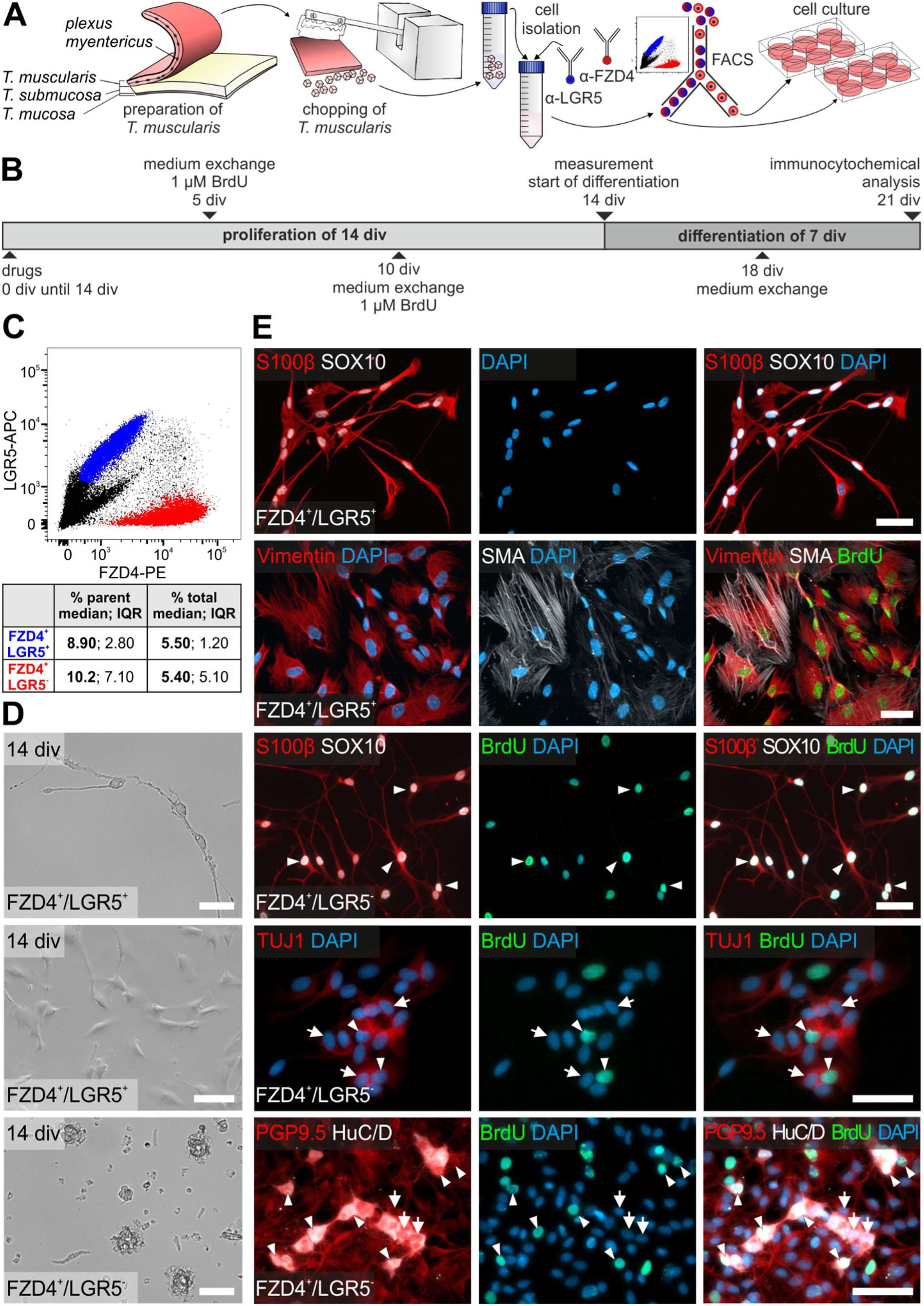
FACS-purified LGR5^-^ human ENS-cells give rise to newborn neurons and glial cells. **A, B:** Cells from human *Tunica muscularis* were sorted using anti-LGR5 and anti-FZD4. After FACS, purified cells were proliferated until 14 div and differentiated for another 7 div. BrdU was added after 5 and 10 div. **C:** Representative FACS-Blot of FZD4^+^LGR5^+^ (blue) vs. FZD4^+^LGR5^-^ (red) cells. The cell populations were readily distinguishable and numerically stable across patients (table in C, n=5). **D:** Representative brightfield images of FZD4^+^LGR5^+^ and FZD4^+^LGR5^-^ cell population after 14 div. FZD4^+^LGR5^-^ cultures gave rise to spheroids under proliferative conditions, whereas in FZD4^+^LGR5^+^ cultures we found neural cells, arguably mature at isolation, and cells with smooth muscle/fibroblast-like morphology. **E:** Representative immunostainings for neuronal markers (PGP9.5, HuC/D, ß-III-tubulin), glial markers (S100beta, SOX10), fibroblasts and smooth muscle cells (vimentin, SMA), as well as BrdU and DAPI in FZD4^+^LGR5^+^ and FZD4^+^LGR5^-^ sorts after 21 div. We found BrdU-co-labeled Vimentin^+^ and SMA^+^ cells in the FZD4^+^LGR5^+^cell population, but not BrdU-incorporating neurons or glial cells, indicating that neurogenic and gliogenic cells were only found in the FZD4^+^LGR5^-^ population (arrows: BrdU^-^ neural cells; arrowheads: BrdU^+^ neurons). **Scale: 50 µm.** See also Supplementary Figure 9. Source data are provided as a Source data file.

After FACS, purified cells proliferated for 14 div and differentiated for another 7 div. After 21 div, cell cultures were fixed and analysed. We found that FZD4^+^LGR5^+^ cells (median: 8.90%; IQR: 2.80% of counts in parent population; n=5) are a numerically stable cell pool clearly distinguishable from the FZD4^+^LGR5^-^ population (median: 10.2%; IQR 7.10% of counts in parent population; n=5; Figure 5C). Under proliferative conditions, only the FZD4^+^LGR5^-^ cell population gave rise to free-floating spheroids (Figure 5D). After differentiation, BrdU-co-labeling demonstrated that several cells were also TUJ1^+^, PGP9.5^+^ and/or HuC/D^+^, indicating that these cells divided and successively differentiated into neurons *in vitro*. However, most neurons were BrdU-negative. Moreover, we detected BrdU^+^SOX10^+^S100beta^+^ glial cells in FZD4^+^LGR5^-^ sorted-cells (Figure 5E).

In contrast, FZD4^+^LGR5^+^ cells were largely adherent under proliferative conditions, with a recognizable differentiated neuronal/glial morphology (Figure 5D). We were not able to detect any neurons in the FZD4^+^LGR5^+^ cell pool after differentiation, arguably due to stressful FACS-purification process. Yet, we found differentiated BrdU^-^ glial cells either SOX10^+^S100beta^+^, or only S100beta^+^. Moreover, these cultures were dominated by BrdU^+^SMA^+^Vimentin^+^ myofibroblast-like cells (Figure 5E).

Thus, unlike the restricted LGR5-expression to proliferative epithelial ICS, LGR5-expression in the ENS was restricted to postmitotic cells. This raises the question if the observed RSPO1-effect is mediated by LGR4 or LGR6.

### LGR4 and LGR6 negative ENS-cells were neurogenic and gliogenic in culture

To validate LGR4 or LGR6 for their neuro/gliogenic potential, we FACS-purified FZD4^+^LGR4^+^ and FZD4^+^LGR6^+^ cells and cultured them as outlined above (Figure 6, Supplementary Figure 9, Supplementary Figure 10). FZD4^+^LGR4^+^ cells and FZD4^+^LGR6^+^ cells were clearly distinguishable from the FZD4^+^LGR4^-^ cells (FZD4^+^LGR4^+^: median: 7.80%; IQR: 3.30% of counts in parent population, and FZD4^+^LGR4^-^: median: 8.25%; IQR: 7.63%; of counts in parent population n=4; Figure 6A) and the FZD4^+^LGR6^-^ cells (FZD4^+^LGR6^+^: median: 7.90%; IQR: 12.5% of counts in parent population, and FZD4^+^LGR6^-^ median: median: 5.40%; IQR: 4.35% of counts in parent population; n=4; Figure 6D). Again, only FZD4^+^LGR4^-^ and FZD4^+^LGR6^-^ cell population gave rise to spheroids during proliferation (Figure 6B and 6E). After differentiation, we detected BrdU^+^SOX10^+^S100beta^+^ (Figure 6C and 6F) and BrdU^-^ S100beta^+^ glial cells in both LGR-negative populations. Additionally, we found BrdU^+^PGP9.5^+^HuC/D^+^ and BrdU^-^PGP9.5^+^HuC/D^+^ enteric neurons in the LGR-negative cultures (Figure 6F).

**Figure 6:**
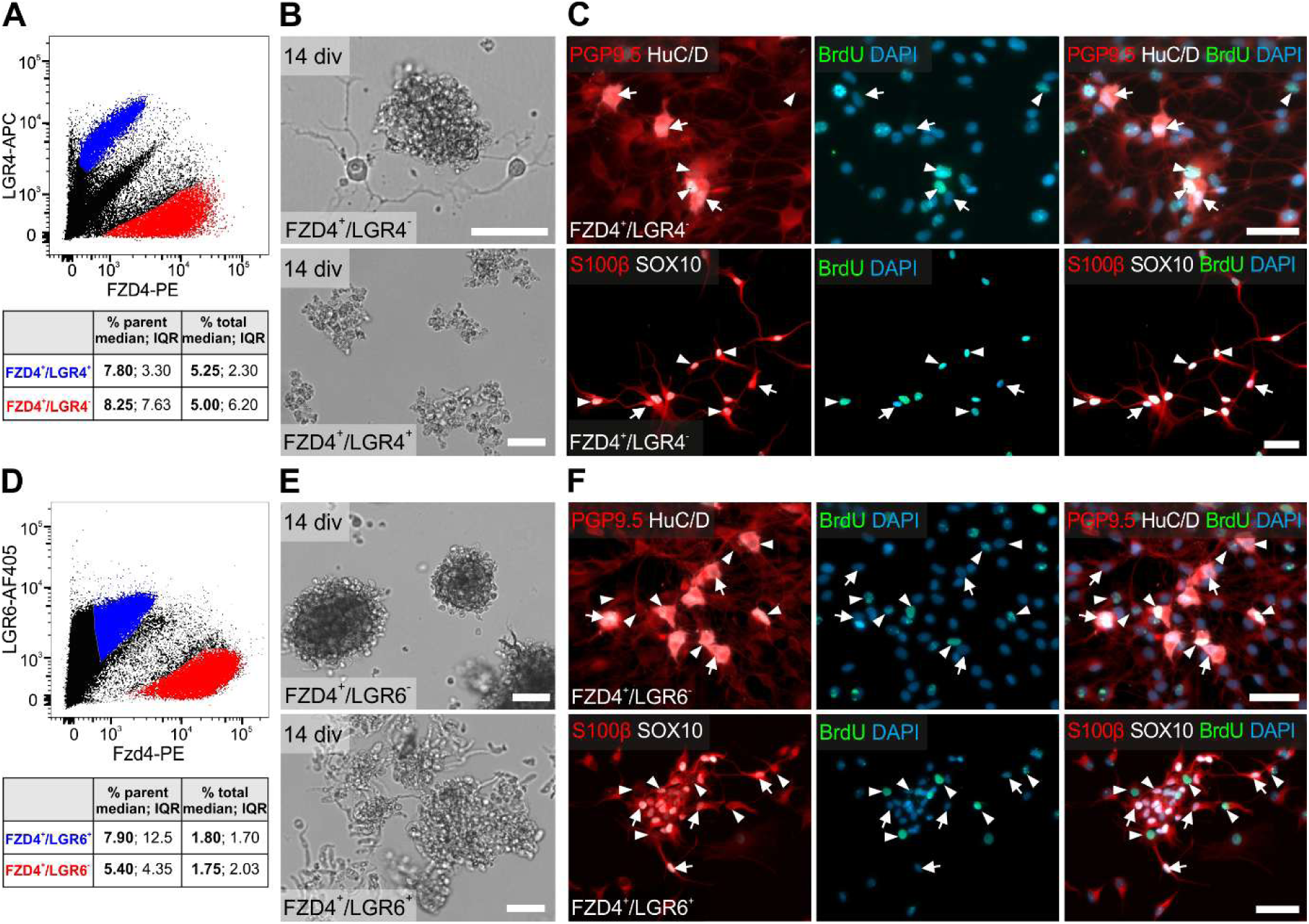
Human LGR4^-^ and LGR6^-^ ENS-cells are neurogenic and gliogenic. **A**: Representative FACS-Blot of FZD4^+^LGR4^+^ cells (blue), numerically stable across patients and readily distinguishable from FZD4^+^LGR4^-^ cells (red; table in A, n=4). **B:** Only FZD4^+^LGR4^-^ cells formed spheroids cells after 14 div; FZD4^+^LGR4^+^ cultures largely exhibited cell debris. **C:** Micrographs depict representative immunostainings for neuronal (PGP9.5, HuC/D) and glial markers (S100beta, SOX10), as well as BrdU and DAPI in FZD4/LGR4-sorted populations. Only FZD4^+^LGR4^-^ cells showed neurogenic and gliogenic potential (BrdU^+^, arrowheads); arrows indicate BrdU^-^ cells. **D:** Representative FACS-Blot of FZD4^+^LGR6^+^ cells (blue), numerically stable across patients and clearly distinct from the FZD4^+^LGR6^-^ cells (red; table in D, n=4). **E:** Only FZD4^+^LGR6^-^ cells formed spheroids *in vitro*; FZD4^+^LGR4^+^ cultures were mostly composed of cell debris. **F:** Representative immunostainings for neuronal (PGP9.5, HuC/D) and glial markers (S100beta, SOX10), as well as BrdU and DAPI in FZD4/LGR6-sorted cell populations. New-born neurons and glial cells were detectable in FZD4^+^LGR6^-^ cultures (BrdU^+^, arrowhead; arrows indicate BrdU^-^ cells). **Scale: 50 µm.** See also Supplementary Figure 9-10. Source data are provided as a Source data file.

In contrast, cell sorting of FZD4^+^LGR4^+^ and FZD4^+^LGR6^+^ populations resulted mostly in non-viable cells during proliferation (Figure 6B and 6E). We were not able to detect neurons or glial cells in FZD4^+^LGR4^+^ and FZD4^+^LGR6^+^ sorted cell pool after differentiation. Again, these cultures were dominated by BrdU^+^Vimentin^+^ and BrdU^+^SMA^+^Vimentin^+^ myofibroblast-like cells (Supplementary Figure 10). Thus, our FACS-data combined with our immunohistochemical analysis showed that especially LGR5- and LGR6-receptors are present on differentiated neurons *in vivo*. Moreover, LGR4 is weakly expressed in the postnatal ENS *in vivo*. The apparent contradiction between our *in vivo* findings versus the RSPO1-mediated pro-proliferative effect *in vitro* might be due to a culture-dependent LGR-receptor expression on ENS-cells during proliferation and differentiation.

### LGR4 and LGR5, but not LGR6, are expressed in proliferative ENS-progenitors

To assess the regulation of LGR-receptors, we used FACS-purified tdTomato^+^ ENS-cells from wnt1-tomato reporter mice (P0, Supplementary Figure 11), seeded them on collagen-coated cover slips, and proliferated them for 5 div. Afterwards, one group was kept under proliferative conditions, whereas the other group entered differentiation. After 6 div, 9 div, and 12 div cultures were fixed and LGR4/5/6-receptor expression was evaluated by immunocytochemistry (Figure 7A).

**Figure 7:**
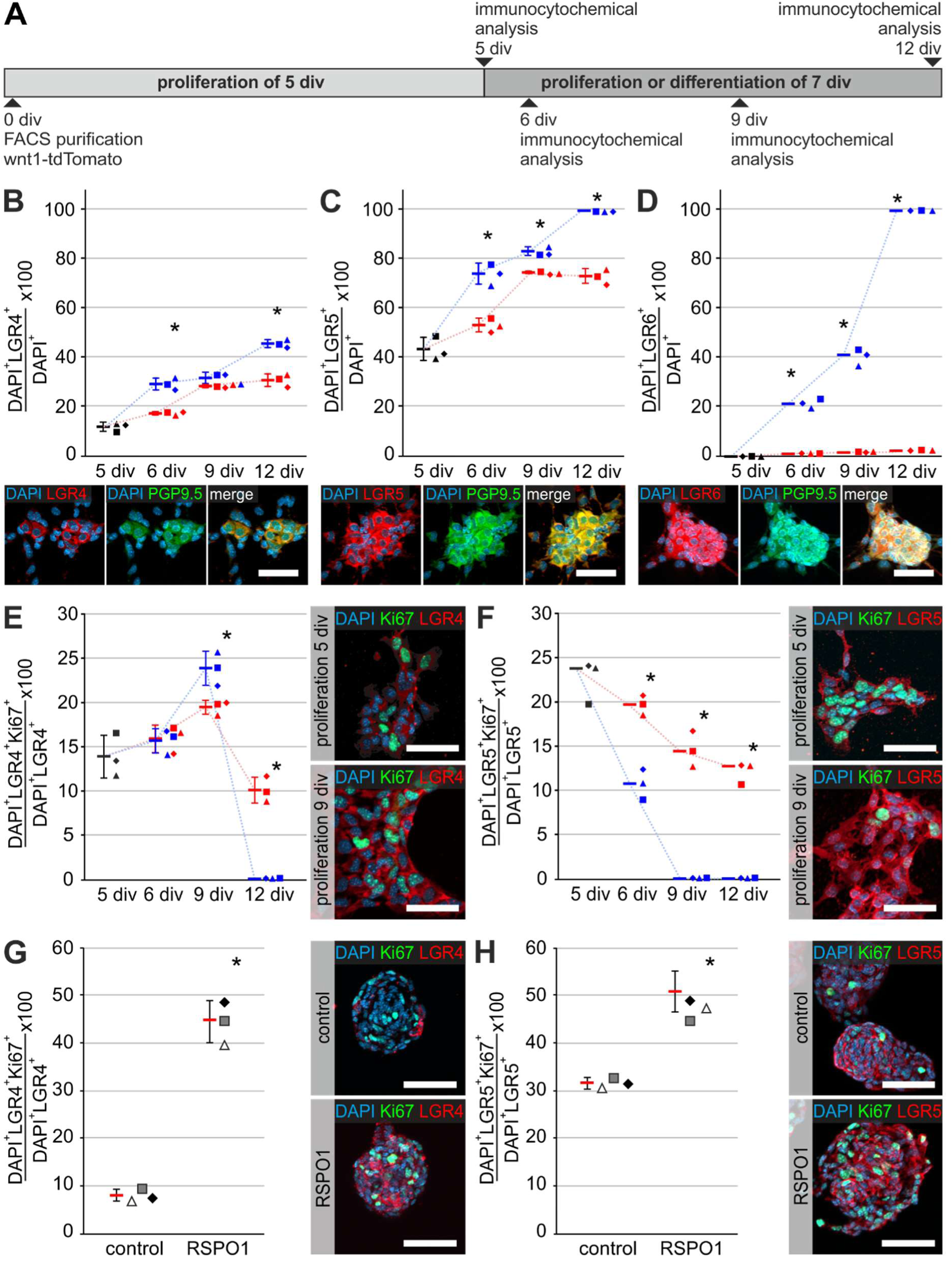
Different roles of LGR4, -5, and -6 in ENS-progenitor proliferation and differentiation. **A:** murine ENS-neurospheres (P0, wnt1-tomato) were adherently proliferated for 5 div. Afterwards, one group was kept under proliferation conditions until 6 div, 9 div, and 12 div, whereas the second group was differentiated. **B-D:** Percentages of LGR4^+^ **(B)**, LGR5^+^ **(C)** and LGR6^+^ **(D)** cells of all DAPI^+^ cells throughout proliferation (red) and differentiation (blue). Asterisk indicates significance (proliferation vs. differentiation); bars and error bars are mean±SD. Independent replicates are represented by different symbols. Micrographs depict representative LGR-stainings, PGP9.5, and DAPI after 9 div during differentiation. The percentages of LGR4^+^/DAPI^+^ and LGR5^+^/DAPI^+^ cells increased in proliferation and differentiation (ANOVA, Fisher LSD; bars and error bars are mean±SD; n=3). LGR6-expression was absent during proliferation, but increased during differentiation (ANOVA on Ranks, Student-Newman-Keuls; bars are medians; n=3). **E-F:** Dotplots illustrate the percentages of proliferative, Ki67^+^/LGR4^+^ (E) or Ki67^+^/LGR5^+^ (F) cells among LGR4^+^ or LGR5^+^ cells, throughout proliferation (red) and differentiation (blue). Asterisk indicates significance (proliferation vs. differentiation); bars and error bars are mean±SD. Independent biological replicates are represented by different symbols. Micrographs depict representative LGR-stainings, Ki67, and DAPI on 5 div and 9 div during proliferation. The percentage of Ki67^+^/LGR4^+^ cells increased during proliferation and differentiation until 9 div (ANOVA, Fisher LSD, bars and error bars are mean±SD; n=3). In contrast, the percentage of Ki67^+^/LGR5^+^ cells decreased constantly *in vitro* (ANOVA on Ranks, Student-Newman-Keuls, bars indicate medians; n=3). **G-H:** RSPO1-stimulated and unstimulated enterospheres were cultured for 5 div. Fixed-frozen cryosections were stained for LGR4 or LGR5, Ki67, and DAPI. Asterisk indicates significant difference to control. Data points for independent replicates are represented by different symbols. RSPO1 stimulation enhanced the percentage Ki67^+^LGR4^+^ by 5.40-fold and Ki67^+^LGR5^+^ cells by 1.62-fold (ANOVA, Fisher LSD, bars and error bars are mean±SD; n=3). **Scale: 50 µm.** See also Supplementary Figure 11-12. Source data are provided as a Source data file.

LGR4-receptor was expressed only on very few cells after 5 div, nevertheless its expression increased throughout the culture, irrespective of the culture-setup. At the end of differentiation, we detected LGR4-expression in differentiated PGP9.5 positive neurons (ANOVA, Fisher LSD post-hoc test, mean±SD proliferation vs. differentiation: 5 div: 11.5%±1.85%; 6 div: 17.0%±0.75% vs. 28.8%±2.40% (P≤0.001); 9 div: 28.0%±0.68% vs. 31.3%±2.25% (P=0.047); 12 div: 30.4%±2.56% vs. 45.2%±1.60% (P≤0.001); n=3; Figure 7B). Interestingly, LGR5-receptor-expression showed a similar trajectory as LGR4. However, LGR5 was more prominent after 5 div. Further, at the end of differentiation (12 div), nearly every PGP9.5^+^ neuron expressed LGR5 (ANOVA, Fisher LSD post-hoc test, mean±SD proliferation vs. differentiation: 5 div: 44.1%±4.67%; 6 div: 53.8%±2.75% vs. 74.5%±4.25% (P≤0.001); 9 div: 75.1%±0.49% vs. 83.6%±1.78% (P=0.003); 12 div: 73.6%±3.01% vs. 100%±0.00% (P≤0.001); n=3; Figure 7C).

In contrast to LGR4 and LGR5, LGR6-expression was hardly present during proliferation, however its expression increased during differentiation until almost every cell expressed LGR6 after 12 div (ANOVA on Ranks, post-hoc: Student-Newman-Keuls Method, median proliferation vs. differentiation: 5 div: 0.01%; 6 div: 1.32% vs. 21.7% (P≤0.05); 9 div: 19.1% vs. 41.4% (P≤0.05); 12 div: 25.6% vs. 100% (P≤0.05); n=3; Figure 7D).

To evaluate proliferation in these cultures, we analysed the expression of Ki67 in LGR4/5/6-expressing cells (Figure 7A). The percentage of Ki67^+^LGR4^+^ cells increased steadily until 9 div under proliferative and differentiation conditions. By 12 div, proliferation rapidly decreased, possibly due to contact inhibition (ANOVA, Fisher LSD post-hoc test, mean±SD proliferation vs. differentiation: 5 div: 13.9%±2.41%; 6 div: 15.9%±1.54% vs. 15.6%±1.37% (P=0.853); 9 div: 19.4%±0.78% vs. 23.8%±1.91% (P=0.003); 12 div: 10.1%±1.44% vs. 0.12%±0.01% (P≤0.001); n=3; Figure 7E). In contrast, the percentage of Ki67^+^LGR5^+^ cells constantly decreased in culture, and we detected hardly any Ki67^+^LGR5^+^ cell after 9 div in differentiation conditions (ANOVA on Ranks, post-hoc: Student-Newman-Keuls Method, median proliferation vs. differentiation: 5 div: 23.8%; 6 div: 19.7% vs. 10.8% (P≤0.05); 9 div: 14.4% vs. 0.02% (P≤0.05); 12 div: 12.7% vs. 0.05% (P≤0.05); n=3; Figure 7F). We did not detect proliferative LGR6 positive cells.

### LGR4^+^ and less so LGR5^+^ cells are drivers of RSPO1-induced neural proliferation

Finally, to assess whether proliferative LGR4- or LGR5-expressing cells are responsive to RSPO1-stimulation, we used fixed-frozen cryosections of enterospheres proliferated for 5 div and stimulated with RSPO1. Untreated enterospheres served as controls. Immunocytochemical analysis for LGR4- and LGR5-receptor (Figure 7G-7H) showed, that the percentage of LGR4^+^DAPI^+^ and LGR5^+^DAPI^+^ cells increased in RSPO1-stimulated spheres compared to controls (LGR4^+^DAPI^+^: ANOVA, Fisher LSD post-hoc test, mean±SD: Control: 8.19%±3.72%; RSPO1: 30.3%±7.53%; n=3; P=0.010; LGR5^+^DAPI^+^: ANOVA, Fisher LSD post-hoc test, mean±SD: Control: 42.6%±7.02%; RSPO1: 52.8%±8.24%; n=3; P=0.179; Supplementary Figure 12). The percentage of rarely found LGR6^+^DAPI^+^ cells did not change (LGR6: ANOVA, Fisher LSD post-hoc test, mean±SD: Control: 4.64%±0.32%; RSPO1: 4.53%±1.74%; n=3; P=0.918, Supplementary Figure 12). Surprisingly, RSPO1-stimulation impressively increased the percentage of Ki67^+^LGR4^+^ by 5.40-fold (ANOVA, Fisher LSD post-hoc test, mean±SD: Control: 8.26%±1.31%; RSPO1: 44.6%±4.37%; n=3; P≤0.001; Figure 7G), whereas it was far less drastic in Ki67^+^LGR5^+^ cells (1.62-fold; ANOVA, Fisher LSD post-hoc test, mean±SD: Control: 31.3%±1.20%; RSPO1: 50.6%±4.80%; n=3; P=0.002; Figure 7H), indicating that the pro-proliferative effect of RSPO1 is primarily mediated by LGR4-expressing cells.

Therefore, our results indicate that RPSO1 acts on ENS-homeostasis depending on the LGR-receptor expressed: while LGR5 and LGR6 might play a role in neuronal differentiation and maintenance, LGR4 arguably mediates pro-proliferative signals and is itself upregulated in proliferative/regenerative microenvironments.

## Discussion

Recent spatial mRNA profiling revealed that Wnt-signaling components, including RSPOs and LGRs, are expressed in the postnatal ENS^28^ suggesting a role of RSPO-LGR signaling in ENS homeostasis. However, LGR-receptor knockouts are associated with severe developmental deficiencies also in the gastrointestinal tract^35,36^ and are not viable until the desired postnatal age. Hence, isolated postnatal murine and patient-derived ENS-progenitors are highly valuable, to evaluate cellular and molecular characteristics of RSPO1-LGR-signaling. Previously we described the expression of Wnt- and RPSO-signaling components in postnatal ENS-progenitors *in vitro*^26,33^, which complements our current results on the presence of *Rspo*-ligands, *Lgr4/5/6*-receptors, and the *Znrf3*-ligase in enterospheres underlining the translational value of this culture-model.

### Cellular effect of RSPO1-stimulation on murine and human ENS-cells

Here, we showed that RSPO1 has a pro-proliferative effect on postnatal murine and human ENS-progenitors leading to a higher yield of newborn BrdU^+^HuC/D^+^ neurons *in vitro*. Similar RPSO1-effects have been reported in neural stem cells (NSCs) from the adult murine hippocampus, where proliferative NSCs expressed LGR4/5, while neighbouring astrocytes shaped a pro-proliferative microenvironment by secreting RSPO1^37^. Thus, an ENS-progenitor niche with secreted Wnt- and RSPO-ligands as well as respective receptors on neural progenitors is conceivable. Indeed, Stavely et al. reported that EMCs secrete Wnt-ligands, *Rspo1,* and *Rspo3*, whereas *Frizzled*- and *Lgr*-receptors are expressed on ENS-progenitors^29^. Additionally, our results on FACS-purified murine ENS-neurospheres suggest that ENS-cells themselves secret Wnt-ligands. This is supported by our recently published mRNA-expression atlas showing that Wntless *(Wls),* which is indispensable for transportation and secretion of Wnt-ligands, is expressed in murine enteric ganglia^28^. Interestingly, this mRNA-expression atlas showed that *Lgr5* and *Lgr6*-transcripts displayed a distinct localization within enteric neuronal perikarya^28^, which is in line with the immune-stainings on human gut sections in the current study. Moreover, single-cell RNA-sequencing data demonstrated the expression of *Rspos* and *Lgr*-receptors in enteric neuronal and glial cell clusters^38^. Therefore, enteric neurons and glial cells might be source and receiver of RSPO-signaling in the ENS.

### Role of LGR4/5/6-receptor expression during postnatal ENS maturation

Differentiation of NSCs in the murine hippocampus is associated with downregulation of LGR4/5-receptor expression, contributing to the attenuation of the canonical Wnt/β- catenin signaling pathway observed in late neural progenitors^39,40^. Similar results have been described for the LGR5-dependent maturation of adult epithelial ISCs^41^, whereby RSPO- and WNT-ligands have qualitatively distinct, non-interchangeable roles. While WNT proteins maintain LGR5-receptor expression on ISCs, RSPO ligands augment ongoing Wnt-signaling, thereby actively controlling the extent of stem-cell expansion^42^. Moreover, RSPO inhibition induces lineage commitment and differentiation by reduced Wnt-signaling activity^43^.

Previously, we showed that canonical Wnt-signaling is inactivated in enterospheres after the induction of cellular differentiation^44^. Strikingly, in the current study the observed pro-proliferative effect of RSPO1 was paralleled by a reduced rate of actively cycling ENS-cells expressing LGR5 *in vitro*. However, we observed that LGR5 was increasingly expressed over the course of enteric neuronal differentiation, which correlates with the observed *in vivo* expression in human gut specimens. Thus, LGR5-expressing ENS-cells might be either less proliferative, or are in a late proliferation-phase, ready to leave the cell-cycle. Still, these comprise a small proportion of cells recruitable to a proliferative state by RSPO1-stimulation. Interestingly, non-cycling ISCs can also express LGR5. These cells are quiescent precursors that committed to the Paneth cell and enteroendocrine lineages. Notably they can be recalled to the stem-cell state after injury^41^.

Intriguingly, in the central nervous system LGR5-expression is predominantly found in postmitotic neurons, partly supporting our results. Miller and colleagues reported that LGR5 and RSPO1 are expressed in murine cerebellar granule neurons. Of note, this expression was restricted to the Wnt-dependent maturation phase and was not observed in adults^45^. Furthermore, predominantly LGR5-expression has been reported in the neurogenic area of the olfactory bulb. However, LGR5-labeling was not detected in stem cells but in fully differentiated neurons of various subtypes and maturation stages^46^.

In contrast to several studies suggesting LGR6-receptor as a stem cell marker and tumour suppressor^47,48^, we barely found LGR6-expression in ENS-cells during proliferation *in vitro*. Similar results have been described for murine adult hippocampal NSCs, where LGR6 exhibit the lowest expression levels of LGR4/5/6 during proliferation *in vitro*^37^. Moreover, we detected that LGR6-expression increased during differentiation suggesting a yet to be elucidated function in neuronal maturation.

Surprisingly, we detected LGR4-receptor expression in actively cycling ENS-cells, which increased during proliferation *in vitro*. Further, although its expression also enhanced towards differentiation of ENS-cells, it was hardly detectable in the mature ENS *in vivo*. Interestingly, murine knockout studies for LGR4- or LGR5-receptor have shown, that LGR4- and LGR5-receptors fulfil comparable functions at least during fetal and postnatal development of the ISCs niche and could compensate each other. However, LGR4 deficiency could not be rescued by LGR5 function in the adult mouse^23,49^, hence highlighting different LGR4/5-receptor functions during proliferation, cell-cycle exit and fate commitment.

LGR4-mediated Wnt-signaling has been reported to function as a protective factor in tissue homeostasis of the gastrointestinal tract *in vivo*. LGR4-deficient mouse models were prone to dextran sodium sulfate (DSS)-induced inflammatory bowel disease (IBD)^50^, related to impaired proliferation and differentiation of intestinal crypts and Paneth cells during tissue regeneration. RSPO1 administration in these mouse models accelerated mucosal regeneration and restoring intestinal architecture, arguably by restored Wnt-signaling activity and stem cell function^51^. Thus, the exerted LGR-function in our study might depend on the developmental and/or regenerative potential of the underlying cellular context, which is arguably more supportive *in vitro* than *in vivo*. To what extent LGR-receptors alone or cooperatively execute proliferation-dependent functions in ENS-progenitors is not assessable with the presented study. Altogether, our results indicate that RSPO1-mediated signaling might have different functional outcomes during ENS-progenitor proliferation and fate commitment via LGR4/5, as well as in maintenance of differentiated ENS-cells arguably via LGR6-expression.

### Clinical implications

Although we did not focus on the role of RSPO-LGR signaling on enteric glial cells, the *in vivo* expression data in this study, as well as recently published data^19,28^ highly suggest a crosstalk with enteric glial cells. Baghdadi et al. highlighted the expression of several Wnt-ligands by enteric glial cells in the *Lamina propria mucosae*. This in turn maintains the self-renewal capacity of epithelial ISCs and influences epithelial regeneration and barrier function in mice^19^. Whether enteric glial cells also communicate bidirectionally with enteric neurons via the Wnt-signaling pathway to influence neuronal or glial homeostasis during postnatal maturation remains elusive. Yet, enteric glial communication with long-lived anti-inflammatory M2 macrophages in the *Tunica muscularis* has been recently outlined to control maintenance of ENS integrity, structure and function^3^. Of note, RSPO1-LGR4 receptor signaling has been reported to initiate a phenotype switch of anti-inflammatory M2 macrophages due to tissue inflammation^52^. Furthermore, alterations in this phenotype have been implicated in gastrointestinal motility disorders like post-operative ileus (POI)^53^. Interestingly, Leven and Schneider et al. demonstrated that enteric glial cells upregulate *LGR4-* and Ki67-expression during the inflammatory manifestation of enteric gliosis in a model of POI. Thereby, the amount of proliferative Ki67^+^/SOX10^+^ cells increased 24h after POI-induction, which was further reflected in an increase in glial cell number during the recovery timepoint 72h later compared to 24h^54^. Thus, a link between RPO-LGR signalling to intestinal inflammation and neural proliferation is conceivable, although the underlying mechanism is yet to be unravelled.

Nevertheless, our data suggests that LGR4 expression could be an indicator for an imminent cell division, arguably triggered by highly proliferative/regenerative conditions (e.g., inflammatory microenvironments or cell cultures). Therefore, conditional knockouts of single LGRs and LGR-combinations are needed in future studies to clarify the *in vivo* role of LGR4/5/6 in ENS homeostasis. Taken together, our results highlight that LGR-receptors serve a diverse function in cellular maturation of postnatal ENS-progenitors *in vitro*. Thereby differential LGR-receptor expression could function as indicators for a turning point in ENS-progenitor fate: a fork stuck in the road either to proliferation or neuronal differentiation.

## Resource availability

### lead contact

Requests for further information and resources should be directed to and will be fulfilled by the lead contact Peter Neckel.

### data and code availability

We provided a Supplementary data file with all numerical source data making up all graphs and charts (*Supplementary Data*).

### material availability

This study did not generate new unique reagents.

## Authors’ contributions

MS: acquisition of data; analysis and interpretation of data; drafting of the manuscript; critical revision of the manuscript for important intellectual content.

SS: acquisition of data; critical revision of the manuscript for important intellectual content.

JF: analysis and interpretation of data; critical revision of the manuscript for important intellectual content.

BH: analysis and interpretation of data; critical revision of the manuscript for important intellectual content.

PHN: study concept and design; acquisition of data; analysis and interpretation of data; drafting of the manuscript; critical revision of the manuscript for important intellectual content; study supervision.

## Acknowledgments

The project was supported by a grant from the German Research Foundation (DFG, Grant number: 438504601). We would like to thank Karin Seid for their technical assistance, Adrian Krestel for his help in the animal facility as well as Andreas Mack for their helpful comments on the manuscript.

## Declaration of interest

The authors declare no competing interests.

## Methods

### Animals

Animals were handled in accordance with national and international guidelines (Notification number AT 01/19 M). Mice were housed in standard cages with standard pathogen-free breeding and a standard 12-hour light/dark cycle at 22 ± 2 °C and 60 ± 5 % humidity. Germ-free food and water were available *ad libitum*. For isolation of unpurified ENS-cultures neonatal (P0-P5, postnatal day 0-5) C57BL/6J mice were used without regard to sex. For FACS-based isolation of ENS cells, we crossbred B6;129S6-Gt(ROSA)26Sortm9(CAG-tdTomato)Hze/J (Jackson Laboratory, Bar Harbor, ME, USA; stock no. 007914) mice with B6.Cg-Tg(Wnt1-cre)2Sor/J (Jackson Laboratory, Bar Harbor, ME, USA; stock no. 022501) mice. The F1 offsprings expressed the red-fluorescent protein tdTomato within all ENS cells and are termed *wnt1-tomato* in this work. For isolation of purified ENS-cultures, neonatal (P0-P5, postnatal day 0-5) and two month (P60, postnatal day 60) old wnt1-tomato mice were used without regard to sex.

### Human Specimens

Human gut samples were obtained from 16 male and 14 female patients aged between 3 days and 12 years, who were operated due to imperforate anus, intestinal obstruction syndrome, or short-gut syndrome (Supplementary Table 1). All samples were collected after approval by the local ethical committee (Project Nr. 652/2019BO2 and 066/2023BO2), and according to the declaration of Helsinki.

### Cell culture of enteric neural progenitors

ENS-progenitors from mice and human patients were isolated from the *Tunica muscularis* and cultured as previously described^33,34^ (see detailed description in the following paragraphs). The culture of unpurified murine and human Tunica muscularis cells (including ENS-progenitors) under proliferation conditions resulted in 3-dimensional spheroids, termed *enterospheres*. In contrast, 3-dimensional spheroids, derived from FACS-purified neural crest−derived cells were termed *neurospheres*. Enterosphere-cultures also comprised non-neuronal cells of the *Tunica muscularis* such as smooth muscle cells or fibroblasts (but not submucosal and mucosal cells, such as epithelial cells). This is an advantage, as it resembles the environment of the *in vivo* situation more accurately.

### Cell isolation of murine enteric neural progenitors

Postnatal day 0-5 old mice were killed by decapitation. P60 old mice were anesthetized with isoflurane, followed by cervical-dislocation. The whole intestine convolute was removed and transferred in *murine preparation buffer* (Hanks’ balanced salt solution (HBSS) without Ca^2+^ / Mg^2+^ (Sigma-Aldrich, Taufkirchen, Germany), penicillin (100 U/mL; PAA, Cambridge,UK), streptomycin (100mg/mL; Sigma-Aldrich, Taufkirchen, Germany)). Adherent mesenteria were dissected, and the longitudinal and circular muscle layers containing myenteric plexus were stripped off the small intestine and collected in murine preperation buffer. After chopping, tissue was enzymatically digested in collagenase type XI (750 U/mL; Sigma-Aldrich, Taufkirchen, Germany)/dispase type II (250 mg/mL; Roche Diagnostics, Mannheim, Germany) solved in HBSS with Ca^2+^ / Mg^2+^ (Sigma-Aldrich, Taufkirchen, Germany) for 20 minutes at 37 °C. Tissue was carefully triturated every 10 minutes with a fire-polished 1-mL pipette tip. Before the first trituration step, cell suspension was treated with 0.05 % (w/v) DNase I (Sigma-Aldrich, Taufkirchen, Germany). After tissue dissociation, 10 % (v/v) fetal calf serum ((FCS), Biochrom, Berlin, Germany) was added. To remove any residual enzymes, two washing steps in HBSS without Ca^2+^ / Mg^2+^, were performed at 200 g. Finally, undigested larger tissue pieces were removed with a 30-µm cell strainer (Miltenyi Biotec GmbH, Bergisch Gladbach, Germany). The pellet was resuspended in *murine proliferation media* (Dulbecco’s modified Eagle’s medium with Ham’s F12 medium (DMEM) 1:1; Life technologies, Darmstadt, Germany) containing N2 supplement (1:100; Life Technologies, Darmstadt, Germany), penicillin (100 U/mL), streptomycin (100 mg/mL), L-glutamine (2 mM; Sigma-Aldrich, Taufkirchen, Germany), EGF (20 ng/mL; Sigma-Aldrich, Taufkirchen, Germany), and hbFGF (20 ng/mL; Sigma-Aldrich, Taufkirchen, Germany). Cells were seeded at a concentration of 2.0×10^4^ cells/cm^2^ and the media was supplemented with B27 (1:50; gibco® Thermo Fisher Scientific, MA, USA) once before seeding. Cells were cultured up to 5 days *in vitro* (div) under proliferation conditions, whereby growth factors (20 ng/ml hEGF/20 ng/ml hbFGF) were added daily. 1 µM BrdU was added after 2 div during the proliferation phase.

### Cell isolation of human enteric neural progenitor cells

For human specimen, the resectates were cut open along the longitudinal axis and rinsed twice with *human preparation buffer* (HBSS without Ca^2+^ / Mg^2+^, penicillin (100 U/mL), streptomycin (100mg/mL), Cibrofloxacin Kabi (5mm/mL; Fresenius Kabi, Bad Homburg, Germany), and metronidazole (50 mg/mL; B. Braun, Melsungen, Germany). The *Tunica adventitia* and scar tissue were removed and the *Tunica muscularis* was peeled off the *Tela submucosa*. *Tunica muscularis* preparations were then stored in *human preparation medium* at 4 °C overnight. On the next day, the *Tunica muscularis* was chopped multiple times (800 mm each) using a Mcllwain tissue chopper (Mickle Laboratory Engineering Co, Guildford, UK). The pieces were enzymatically digested in collagenase type XI (750 U/mL)/dispase type II (250 mg/mL) dissolved in HBSS with Ca^2+^ / Mg^2+^ containing 0.05 % (w/v) DNase I and incubated for up to 60 minutes at 37°C.

Tissue was triturated every 20 minutes with a fire-polished 25-mL serologic pipette. After tissue dissociation, fetal calf serum was added to a concentration of 10 % (v/v). The cells were pelleted at 200 g and erythrocyte lysis was performed using RBC Lysis buffer (eBioscience, Frankfurt a.M., Germany). After a second centrifugation at 200 g, the pellet was resuspended in HBSS without Ca^2+^ / Mg^2+^ and filtered using 100-µm, and 70-µm cell strainers. Cells were pelleted again at 200 g and resuspended in *human proliferation medium* (DMEM containing N2 supplement (1:100), penicillin (100 U/mL), streptomycin (100 mg/mL), L-glutamine (2 mM), EGF (20 ng/mL) and hbFGF (20 ng/mL). Cell suspension was finally filtered using a 30-µm cell strainer and seeded in a concentration of 2.0×10^4^ cells/cm^2^. The medium was supplemented with B27 (1:50) once before seeding. Cells were cultured up to 14 days *in vitro* (14 div) under proliferation conditions, whereby growth factors (20 ng/ml hEGF/20 ng/ml hbFGF) were added daily, and culture medium was exchanged every 5 days. 1 µM BrdU was added after 5 div and 10 div during the proliferation phase.

### Fluorescence-Activated Cell Sorting of purified murine cultures

For FACS analysis, cells were isolated from the *Tunica muscularis* of P0-P5, and P60 wnt1-tomato mice using the same procedure described above. After the 30-μm straining step, cells were collected in Hibernate A (gibco® Thermo Fisher Scientific, MA, USA) medium supplemented with N2 supplement (1:100), penicillin (100 U/mL), streptomycin (100 mg/mL), L-glutamine (2 mM), EGF (20 ng/mL), and hbFGF (20 ng/mL) and B27 (1:50). FACS was then performed with a BD FACS Aria flow cytometer (BD Biosciences) using a 100-μm nozzle. Forward-sideward scatter dot plots were used to exclude debris and cell aggregates. Endogenous tdTomato was excited by a 488-nm laser. Emission filter was 576/26 nm. For all experiments, purified cells were seeded as outlined above at a concentration of 1.0×10^5^ counts/cm^2^.

### Fluorescence-Activated Cell Sorting of human LGR-FZD4 cultures

For FACS analysis, cells were isolated from the *Tunica muscularis* of human gut specimens as outlined above. After filtration through a 30-µm cell strainer, cells were pelleted at 200 g, resuspended in 20 µl Gamunex (10%-solution, Talecris Biotherapeutics, NY, USA) per 10^6^ cells isolated and incubated for 20 minutes on ice. Then, Fzd4-antibody combined with LGR4- or LGR5- or LGR6-antibody were added in the corresponding concentrations as outlined in the Key Resource Table and incubated for 15 minutes on ice. After antibody incubation, cells were washed two times in Hibernate A supplemented with N2 supplement (1:100), penicillin (100 U/mL), streptomycin (100 mg/mL), L-glutamine (2 mM), EGF (20 ng/mL), and hbFGF (20 ng/mL) and B27 (1:50). FACS-analysis was then performed with a BD FACS Aria flow cytometer using a 100- μm nozzle. Forward-sideward scatter dot plots were used to exclude debris and cell aggregates. Excitation and emission filter settings are described in the Key Resource Table. Purified cells were seeded as outlined above at a concentration of 2.0×10^5^ counts/cm^2^. The medium was supplemented with B27 (1:50) once before seeding. Cells were cultured up to 14 days *in vitro* (14 div) under proliferation conditions, whereby growth factors (20 ng/ml hEGF/20 ng/ml hbFGF), WNT3A (20 ng/ml, R&D Systems, Inc., MN, USA), R-Spondin1 (100 ng/ml, R&D Systems, Inc., MN, USA), as well as Rock Inhibitor (1µM; Selleck Chemicals GmbH, Köln, Germany) were added daily and culture medium was exchanged every 5 days. 1 µM BrdU was added after 5 div and 10 div during the proliferation phase.

### Stimulation of murine and human enteric progenitor cells

For cell culture experiments, the medium was supplemented either with WNT3A (20 ng/ml), R-Spondin1 (100 ng/ml) or the combination of both after 1 day *in vitro* (1 div) for murine cultures or on isolation day (0 div) for human cultures. Untreated cells served as control group. To quantify the absolute number of proliferating enteropsheres derived from neonate mice and human specimens brightfield images were taken of stimulated and unstimulated enteropsheres at 5 div respectively for human cultures at 14 div.

### Differentiation of murine and human enteric progenitor cells

After proliferation phase (5 div for murine and 14 div for human cultures), cell cultures were treated with 10 % (v/v) FCS for 2 hours to facilitate attachment of entero- and neurospheres to the culture dishes. Afterwards, *differentiation medium* (DMEM containing N2 supplement (1:100), penicillin (100 U/mL), streptomycin (100 mg/mL), L-glutamine (2 mM), 2 % (v/v) FCS and ascorbic acid-2-phosphate (200 mM; Sigma-Aldrich, Taufkirchen, Germany) was added and exchanged after 9 div for murine cultures and after 18 div for human cultures.

After the differentiation phase (12 div for murine, and 21 div for human cultures) cell cultures were fixed for immunocytochemical analysis.

### Single sphere assay

For single sphere assay, cells were isolated from the *Tunica muscularis* of wnt1-tomato mice aged two months (P60), using the same procedure described above for wild-type mice. After FACS-sorting, purified cells were seeded as outlined above at a concentration of 2.0×10^5^ cells/cm^2^. Cells were cultured up to 7 days *in vitro* (7 div) under proliferation conditions, whereby growth factors (20 ng/ml hEGF/20 ng/ml hbFGF) were added daily. After 7 div, single neurospheres (40-60µm diameter) were picked and transferred into a 96-well plate with *proliferation medium* (DMEM containing N2 supplement (1:100), penicillin (100 U/mL), streptomycin (100 mg/mL), L-glutamine (2 mM), EGF (20 ng/mL) and hbFGF (20 ng/mL). Before pharmacological stimulation, brightfield images were taken from each single sphere. Afterwards, the medium was supplemented either with WNT3A (20 ng/ml), R-Spondin1 (100 ng/ml) or the combination of both. Single spheres were cultivated under proliferative conditions, whereby growth factors (20 ng/ml hEGF/20 ng/ml hbFGF) were added daily. To measure diameter changes of stimulated neurospheres, brightfield images were taken after 14 days *in vitro*.

### Time-dependent LGR expression

For time series experiments, cells were isolated from the *Tunica muscularis* of wnt1-tomato mice using the same procedure described here for wild-type mice. After FACS-sorting, purified cells were seeded as outlined above at a concentration of 1.0×10^5^ counts/cm^2^ on collagen-type-I (Merck KGaA, Darmstadt, Germany) coated coverslips and cultivated under proliferative conditions, whereby growth factors (20 ng/ml hEGF/20 ng/ml hbFGF) were added daily. On 5 div, cells were cultivated either under proliferative or differentiation conditions, as already outlined above. On 6 div, 9 div and 12 div cells were fixed for immunocytochemical analysis.

### Immunostainings for differentiation markers and BrdU proliferation assay

After differentiation phase, murine neonate wildtype and human cell cultures were fixed with 4 % (w/v) phosphate buffered paraformaldehyde ((PFA), Merck KGaA, Darmstadt, Germany) for 20 minutes at room temperature and subsequently rinsed three times for 5 minutes in phosphate-buffered saline (PBS). To avoid unspecific binding of antibodies and to permeabilize cells for the detection of intracellular proteins, cell cultures were pre-treated with blocking solution containing 4 % (v/v) goat serum (Biochrom, Berlin, Germany), 0.1 % (w/v) bovine serum albumin ((BSA), Roth, Karlsruhe, Germany), and 0.1 % (v/v) Triton^®^ X-100 (Roth, Karlsruhe, Germany), for 30 minutes at room temperature. Afterwards, primary antibodies (Key Resource Table) diluted in PBS, 0.1 % (w/v) BSA, and 0.1 % (v/v) Triton^®^ X-100, were added and incubated overnight at 4 °C. Afterwards, cell cultures were rinsed with PBS three times for 5 minutes. Fluorescent conjugated secondary antibodies (Key Resource Table), diluted in PBS containing 0.1 % (w/v) BSA and 0.1 % (v/v) Triton^®^ X-100, were used for detection of primary antibodies and were incubated for one hour, light protected, at room temperature. Nuclear staining was performed together with the secondary antibody, using 4’,6-diamidino-2-phenylindolestain (DAPI 200 ng/ml, Roth, Karlsruhe, Germany).

For BrdU proliferation assay, cell cultures were pre-treated with 2 N HCl (Roth, Karlsruhe, Germany) in a humidity chamber for 30 minutes at 37 °C after the incubation of fluorescent conjugated secondary antibodies. Next, three washing steps were carried out, twice with borax buffer (0.1 M (w/v) di-Natriumtetraborat-10-hydrat pH: 8.5, Merck, Darmstadt Germany) and once with PBS for 5 minutes followed by incubation of primary BrdU antibody (Key Resource Table) diluted in PBS containing 0.1 % BSA, and 0.1 % (v/v) Triton^®^ X-100, in a humidity-chamber for 2 hours at 37 °C. Upon three washing steps with PBS for 5 minutes, cell cultures were treated with diluted secondary antibody in PBS containing 0.1 % (w/v) BSA, and 0.1 % (v/v) Triton^®^ X-100, together with DAPI (200 ng/ml) for 1 hour at room temperature. Eventually, cell cultures were rinsed again three times with PBS for 5 minutes.

### Immunohistochemistry and Ki67 stainings on 5-day-old enteropsheres

To assess proliferation within murine enteropsheres during the proliferation phase, 5-day-old enteropsheres were picked from the medium and fixed with 4 % (w/v) PFA for 20 minutes and rinsed three times with PBS. For cryoconservation, fixed enteropsheres were stored overnight in 30 % (w/v) sucrose solution (Applichem, Darmstadt, Germany) at 4 °C. Afterwards, samples were frozen in isopentane-nitrogen cooled TissueTek^®^ (Sakura, Staufen, Germany) and stored at -80 °C until further processing. Before staining, cryosections (10 µm) were dried for one hour at room temperature, following rehydration with distilled water for 15 minutes.

For paraffin embedding, fixed tissue samples were dehydrated in an ascending alcohol series, followed by xylene and overnight infiltration of Paraffin at 60 °C. Before staining, paraffin sections (5 µm) were dewaxed by xylene (twice, 5 minutes) and a descending alcohol series (once 100 % ethanol, once 96 % (v/v) ethanol, and once 70 % (v/v) ethanol, 5 minutes each). Finally, slides were rinsed once with distilled water. Next, sections were pre-treated with boiled citric acid monohydrate buffer (10 mM, pH 6.0, Merck, Darmstadt, Germany) for three minutes and cooled down at room temperature.

To prevent unspecific binding of antibodies, fixed enteropsheres and tissue samples were blocked for 30 minutes with PBS containing 4 % (v/v) goat serum, 0.1 % (w/v) BSA and 0.1 % (v/v) Triton^®^ X-100, followed by incubation of primary antibodies (Key Resource Table) diluted in PBS with 0.1 % (w/v) BSA and 0.1 % (v/v) Triton^®^ X-100 overnight at 4 °C in a humidity chamber. Afterwards, samples were washed with PBS three times for 5 to 10 minutes. The secondary antibody (Key Resource Table) was diluted in PBS, 0.1 % (v/v) Triton X^®^-100, and 0.1 % (w/v) BSA, and incubated for 60 minutes at room temperature. Nuclear staining was carried out with DAPI (200 ng/ml). After two washing steps with PBS for 5 minutes, the samples were washed in distilled water for 5 minutes, followed by mounting with Kaiser’s glycerol gelatine (Merck, Darmstadt, Germany).

### RNA isolation and reverse transcriptase-PCR

Total RNA of enteropsheres was isolated using the RNeasy Plus Mini Kit (Qiagen, Hilden, Germany) according to manufacturer’s instructions. RNA concentration and integrity were analysed using QIAxcel Advanced (Qiagen, Hilden, Germany) according to manufacturer’s instructions. Reverse transcription was carried out with QuantiTect Reverse Transcription Kit (Qiagen, Hilden, Germany). Purified RNA treated without reverse transcriptase served as a negative control. PCR was performed using the StepOnePlus™ Real-Time PCR System (Applied Biosystems, Darmstadt, Germany) and the PerfeCTa qPCR ToughMix ROX (Quantabio, Berverly, MA, USA) according to the manual instructions. The PCR conditions were 50 °C for 120 s and 95 °C for 10 min followed by 40 cycles of 95 °C for 15 s and 60 °C for 60 s. The acquisition was performed after the 60 °C step of each cycle. Glyceraldehyde 3-phosphate dehydrogenase (GAPDH), hypoxanthine-guanine phosphoribosyltrans-ferase (HPRT), and the TATA-binding protein (TBP) were used as reference genes. qRT-PCR was carried out following MIQE quality guidelines. Primers for ligand and receptors are shown in Key Resource Table.

### Microscopy

Images were acquired using a Zeiss Axio Imager.Z1 fluorescence microscope with Apotome module with 358, 488, 543, 647 nm for excitation and appropriate filter sets. Images were acquired using ZEN software, as well as using a the 10-objective (EC Plan-Neofluar 10x/0.30 Ph1), whereby exposure times for DAPI was 500-1000 ms, and neuronal and glial markers 1500-2500 ms. Additionally, the 20-objective (Plan-Apochromat 20x/0.8 M27) with exposure times for DAPI <100 ms, Ki67 500 ms, LGR-receptors 500-1000 ms and for the neuronal and glial markers 500-1000 ms was used.

### Data Analysis

For analysis of sphere growth, bright-field images were taken after 5 div for murine and after 14 div for human cell cultivation. In total 7452 murine enteropsheres and 4176 human enteropsheres were analysed. The area of spheroids was measured on bright-field images using Axiovision software (Zeiss, Oberkochen, Germany). Assuming an ideal sphere shape of enteropsheres, the theoretical diameter (d) was calculated by equation [1]: 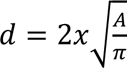 in µm whereby A is the measured area. Further, the total cell volume was calculated by equation [2]: 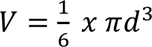 to evaluate a possible pharmacologically effect on the total cell volume of counted enteropsheres, whereby V is the obtained volume in µm³. It should be mentioned that human progenitors do not form compact spheroids with smooth edges as compared to mouse spheroids, for this reason we measured the dense compact core. For all, cumulative sphere number as well as volume was normalized to controls to calculate fold change. For each group two technical replicates were analysed. For the quantification of Ki67^+^/P75^+^/LGR4-6^+^ cells in 5-day-old enteropsheres, at least 500 DAPI positive nuclei with the respective co-labeling event was counted to calculate percentages. To quantify differentiated murine enteric neurons from unpurified cultures two technical replicates per experimental group were counted manually and mean values were calculated. To evaluate differentiated murine enteric neurons from purified cultures one technical replicate from small and large intestinal samples per experimental group were counted manually. For the analysis of human differentiation data four technical replicates per experimental group were manually counted and median value was calculated from four biological replicates. qRT-PCR experiments were carried out following MIQE quality guidelines with six independent experiments.

### Statistics and Reproducibility

For all experiments, at least three independent experiments (independent preparations on different days) were carried out. Statistical analysis was performed using SigmaStat 3.5 software. Results were considered significant at P≤0.05. All statistical tests and p-values are provided in the Source Data File.

## Supplementary Data

*Supplementary Figures 1-12*, Supplementary Table 1-2, Key Resource Table, Source data file

**Author names in bold designate shared co-first authorship**

## Notes

### Competing Interest Statement

The authors have declared no competing interest.

