## Supplementary Data for "R-Spondin1 regulates fate of enteric neural progenitors via differential LGR4/5/6-expression in mice and humans"

**Supplementary Figures**

| ligand receptor | mean CT value | ± SD CT value |
|---|---|---|
| *Rspo1* | 31.47 | 1.53 |
| *Rspo2* | 30.50 | 0.82 |
| *Rspo3* | 30.67 | 0.81 |
| *Rspo4* | 34.33 | 0.70 |
| *Lgr4* | 30.11 | 3.13 |
| *Lgr5* | 34.49 | 0.96 |
| *Lgr6* | 33.08 | 1.87 |
| *Znfr3* | 30.11 | 1.11 |
| *Rnf43* | not detectable | |
| *Gapdh* | 20.34 | 2.04 |
| *Hprt* | 26.97 | 2.15 |
| *Tbp* | 27.90 | 1.13 |

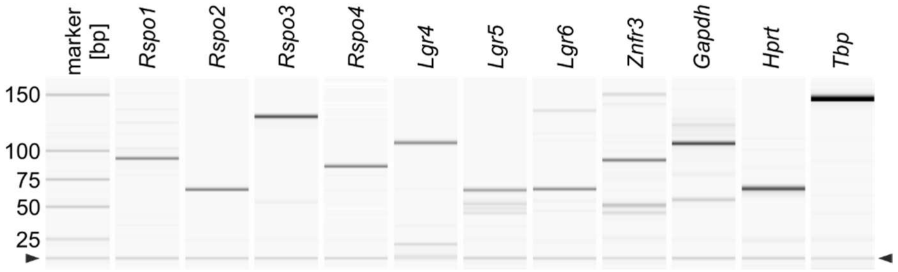


**Suppl. Fig. 1: RSPO-ligands and LGR-receptors are expressed in murine ENS-progenitors.** Identification of RSPO-ligands *Rspo1-Rspo4*, RSPO-receptor *Lgr4, Lgr5 and Lgr6,* as well as the transmembrane ligase *Znfr3* in 5-days-old enterospheres by qRT-PCR (CT-values expressed as mean±SD) and by capillary electrophoresis (arrowhead indicates position of alignment marker). Related to Figure 1.

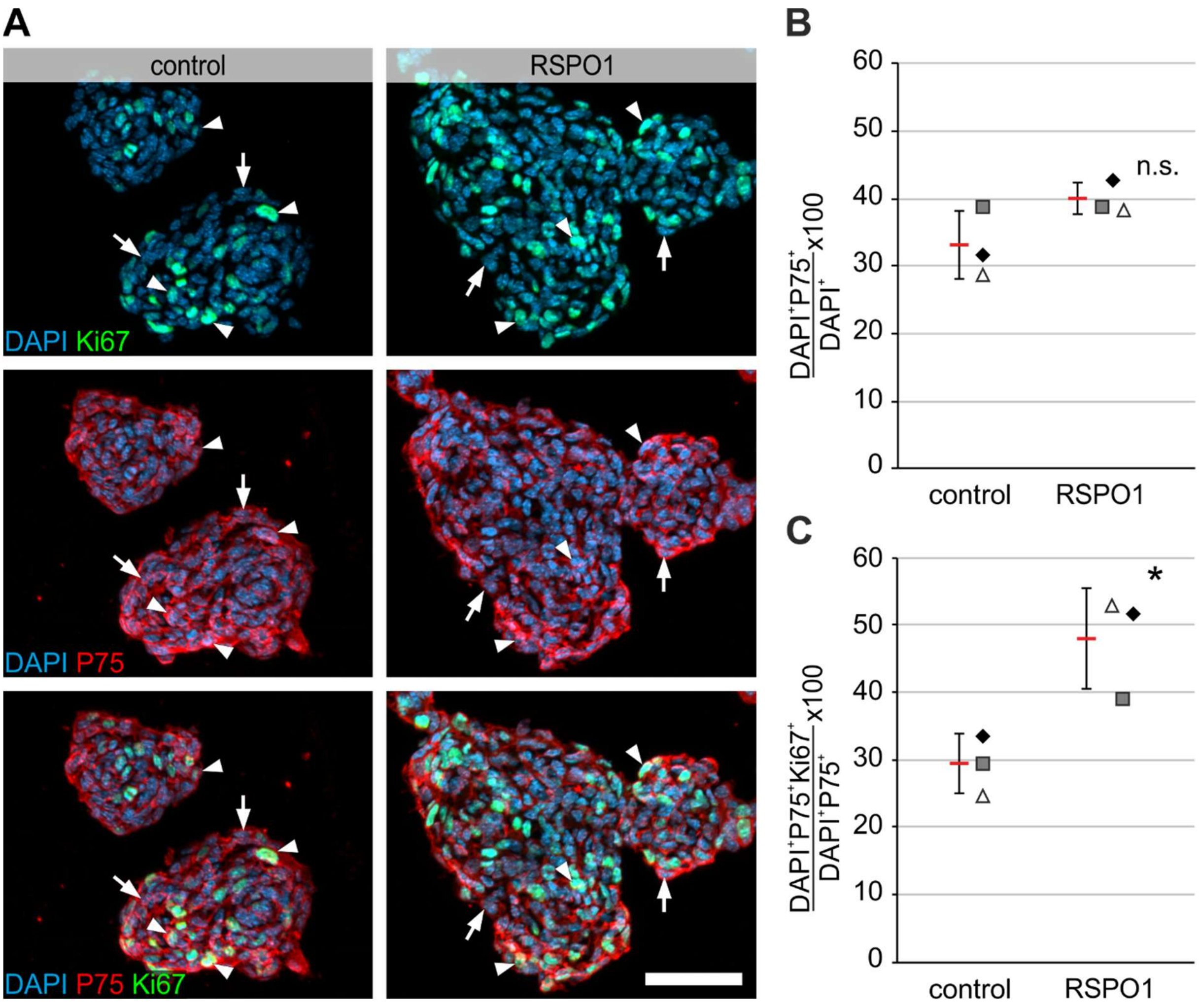


**Suppl. Fig. 2: RSPO1-stimulation increased the amount of proliferative Ki67$^+$P75$^+$ neural cells. A:** Micrographs display immunofluorescence co-labeling studies with Ki67 (green) and P75 (red) and the nuclear marker DAPI (blue), on paraffin-sections of enterospheres, that were cultured for 5 days *in-vitro* (5 div) under proliferative conditions. RSPO1 was applied after 1 div. Untreated enterospheres served as control, **scale bar: 50 µm. B:** The dotplots indicate the percentage of P75$^+$DAPI$^+$ and in **C:** proliferative P75$^+$Ki67$^+$DAPI$^+$ cells (mean±SD) for the control and RSPO1-stimulated group. Asterisk indicates significant differences in comparison to the control group. Data points for independent biological replicates are represented by different symbols. RSPO1-stimulation had no significant effect on P75$^+$DAPI$^+$ cells **(B),** (ANOVA, Fisher LSD post-hoc test, mean±SD: Control: 33.0%±5.06%; RSPO1: 39.9%±2.31%; n=3;

P=0.098), yet increased the number of proliferative Ki67$^{+}$P75$^{+}$DAPI$^{+}$ **(C),** (ANOVA, Fisher LSD, mean±SD: Control: 28.7%±5.58%; RSPO1: 48.1%±7.55%; n=3; P=0.023). Thus, suggesting, that within all P75$^{+}$ neural cells, a sup-population is capable to proliferate in response to RSPO1-stimulation. Related to Figure 1. Source data are provided as a Source data file.

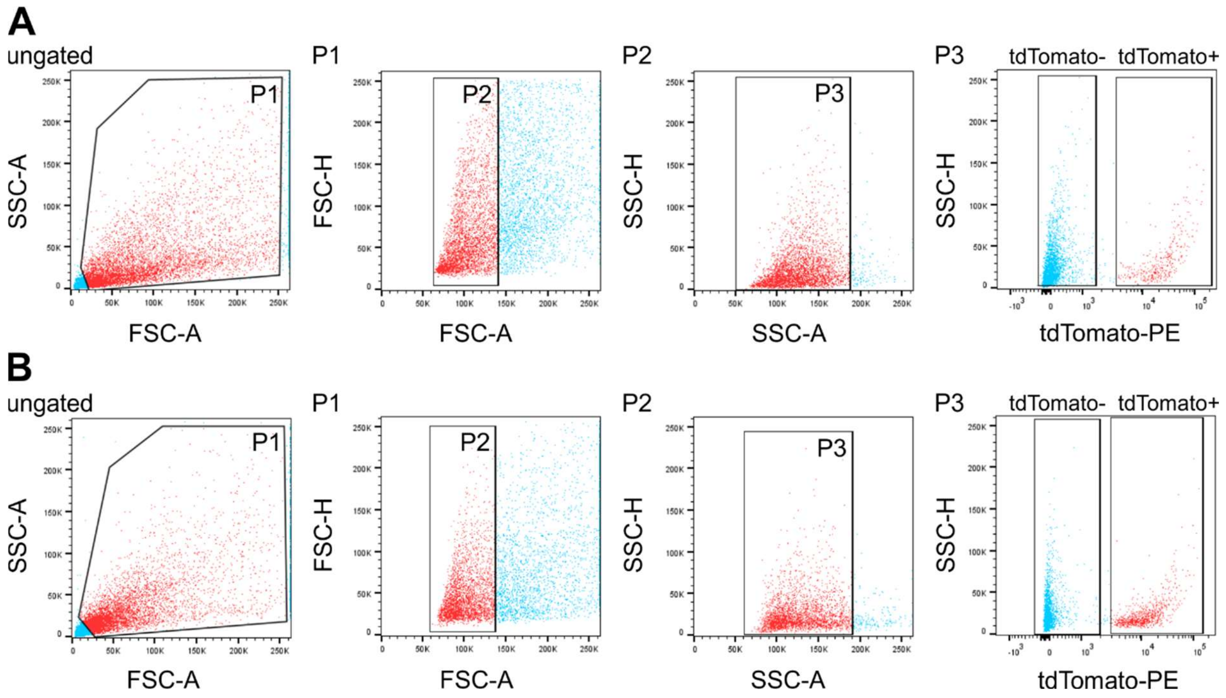


**Suppl. Fig. 3: Fluorescence-activated cell sorting (FACS) of tdTomato expressing ENS-cells derived from postnatal day 60 old Wnt1Cre2 mice.** The scatter blots represent the gating strategy (ungated, P1–P3) for tdTomato negative (P4) and tdTomato positive (P5) cell pools derived from small and large intestine. Each population, that underwent further gating are colored in red. All events counted were first gated as P1 to exclude dead cells and debris, as these could be found at the bottom left corner of the first dot blot of each sample. Next, P1 was gated regarding its properties in forward (P2) and sideward (P3) scatter mode to exclude doublets and aggregates. Finally, P3 was gated regarding its tdTomato-expressing property. See also the supplementary data file, that summarizes the gating data for all experiments used in this study. Related to Figure 2. Source data are provided as a Source data file.

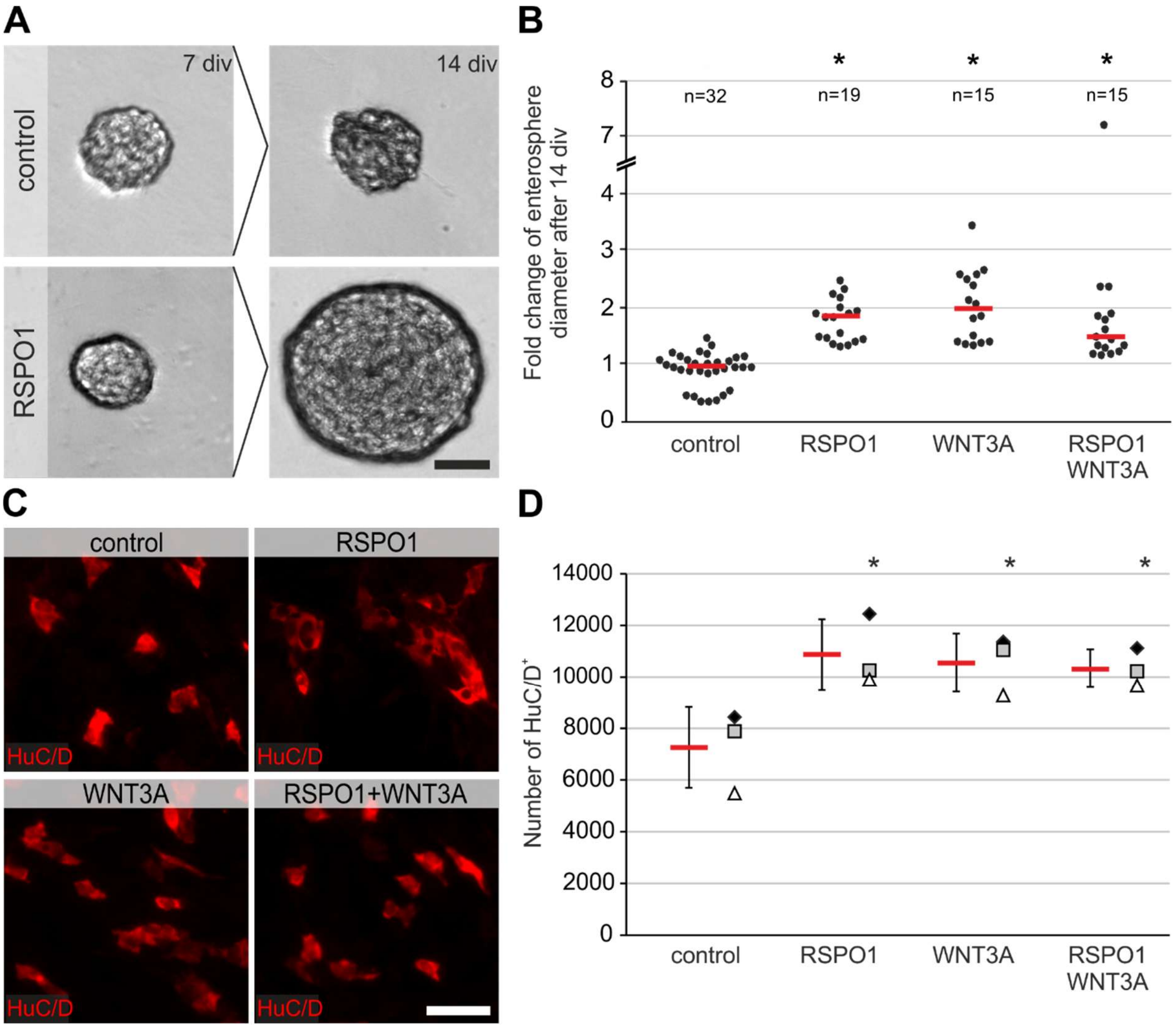


**Suppl. Fig. 4: RSPO1-mediated effect acts directly on large intestine derived ENS-progenitors.** FACS-purified ENS-progenitors of large intestinal samples from two-month-old (P60) mice were cultured for 7 days under proliferative conditions *in vitro* (7 div) as a multisphere culture. After 7 div, evenly sized neurospheres were selected, transferred into a 96 well plate, and cultivated under proliferative conditions as single sphere culture for further 7 div. Brightfield images were taken, before RSPO1 and/or WNT3A were added and again after 14 div. **A:** shows a representative image of the same single sphere derived from large intestine on 7 div and 14 div under control and RSPO1-treated condition. **Scale bar: 40 µm. B:** Dot plot depicts the fold change of neurosphere diameter after 14 div. Each dot represents one single neurosphere analyzed, red bar represents the median. RSPO1-stimulation had a comparable effect as WNT3A stimulation alone (ANOVA on Ranks, post-hoc: Dunn's method, median fold change: control: 1.02, n=32; RSPO1: 1.95, n=19, (P≤0.001); WNT3A: 2.08, n=15, (P≤0.001), RSPO1+WNT3A: 1.56, n=15, (P≤0.001); four biological replicates). Asterisk indicates significant differences compared to control. **C:** Beside single sphere assays, FACS-purified ENS-cells were cultivated as multisphere cultures until 7 div under proliferative conditions, stimulated with RSPO1 and/or WNT3A on the day of seeding (div 0). After 7 div, cells were cultured for 7 div under differentiation conditions, and underwent immunocytochemical analysis for HuC/D. Micrographs show representative images of HuC/D$^{+}$ neurons (red) in all experimental groups after differentiation. **Scale bar: 50 µm. D:** displays the quantification of the number of HuC/D$^{+}$ neurons. Data points for different experimental groups are represented by different symbols. Number of neurons increased after RSPO1-stimulation compared to untreated control (ANOVA, Fisher LSD post-hoc test, mean±SD: control: 7246±1564; RSPO1: 10844±1375 (P=0.007); WNT3A: 10543±1126 (P<0.012), RSPO1+WNT3A: 10319±735 (P=0.016); n=3). Asterisk indicates significant differences compared to control. Related to Figure 2. Source data are provided as a Source data file.

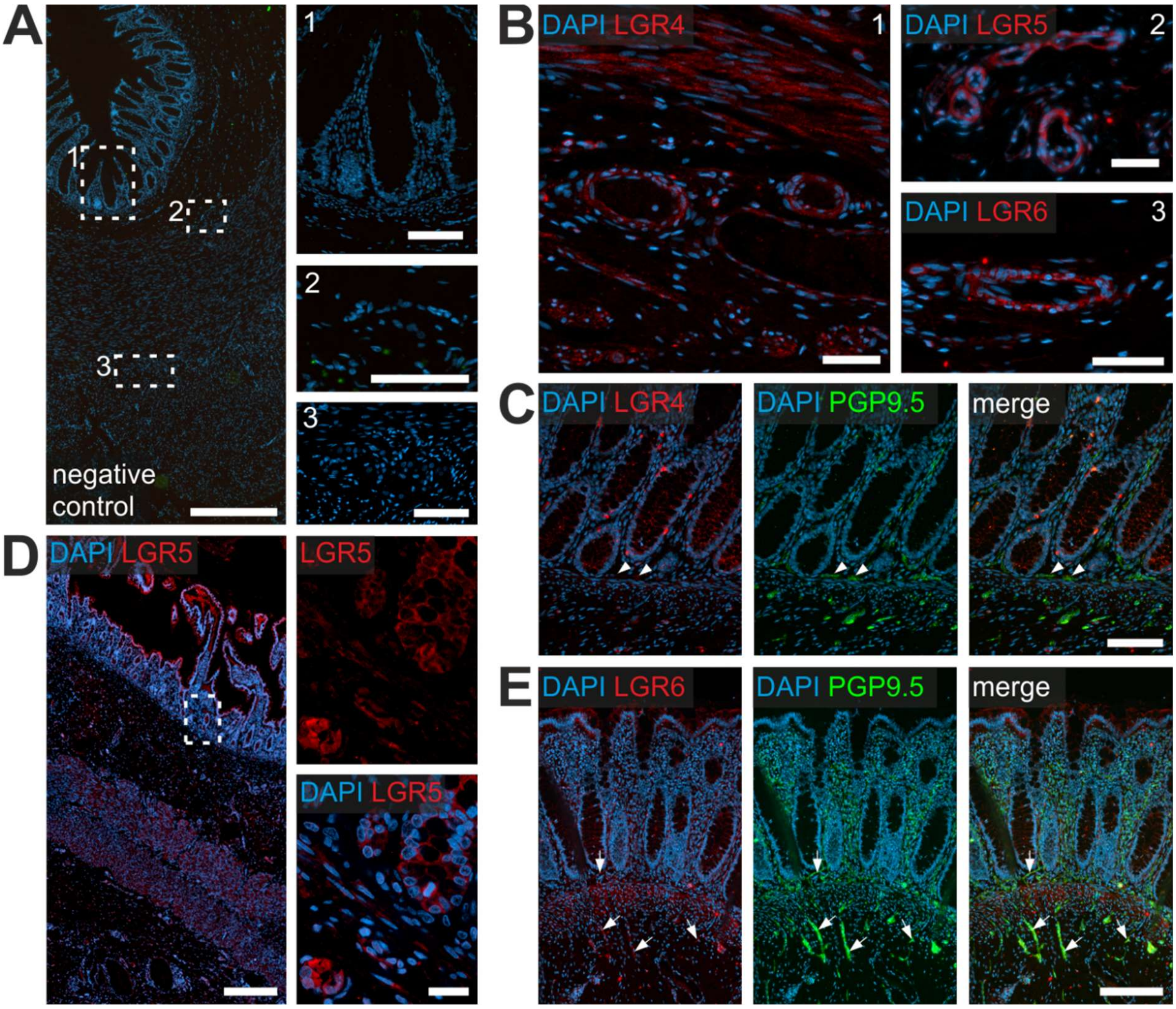


**Suppl. Fig. 5: LGR-receptor expression in other compartments of small intestine. A:** displays an overview of a transversal section of the human small intestine section stained with secondary antibodies only as a negative control and the nuclear stain DAPI (blue). White rectangles indicate the location of the high-power magnification micrographs 1-3 on the right. **Scale bars:** overview **500 µm**; details: **100 µm** (1), **50 µm** (2-3). **B:** show representative images of blood vessels located in the *Tela submucosa*. LGR4 (1), LGR5 (2) and LGR6 (3), (red) are expressed in the *Tunica media* of arterioles. Further we found a slight LGR4-expression in the *Lamina muscularis mucosae*. Nuclear marker DAPI (blue). **Scale bars: 50 µm (B1-B2), 25 µm (B3). C:** depicts a representative image of small intestine sample stained for LGR4 (red), the neuronal marker PGP9.5 (green), and the nuclear marker DAPI (blue). LGR4 was expressed throughout the crypt region till the villus border. Further we detected LGR4 co-localization with PGP9.5-co-labeled neurites surrounding the mucosal crypts (arrowheads), **Scale bar:100 µm**. **D:** depicts small intestinal sample stained for LGR5 and the nuclear marker DAPI (blue). LGR5-expression was restricted to epithelial cells located at the crypt-bottom (high power magnification), **scale bars:** overview **600 µm**; details: **20 µm**. **E:** shows representative image of human small intestine stained for stained for LGR6 (red), PGP9.5 (green) and the nuclear stain DAPI (blue). LGR6 was expressed throughout the *Tunica mucosa* predominantly in the *Lamina muscularis mucosa.* In addition, we detected some staining for LGR6 in PGP9.5-co-labeled neurites outside the ganglia within the *Tunica muscularis* and in the fine neurite network surrounding the mucosal crypts (arrow). Related to Figure 4. **Scale bar:100 µm**.

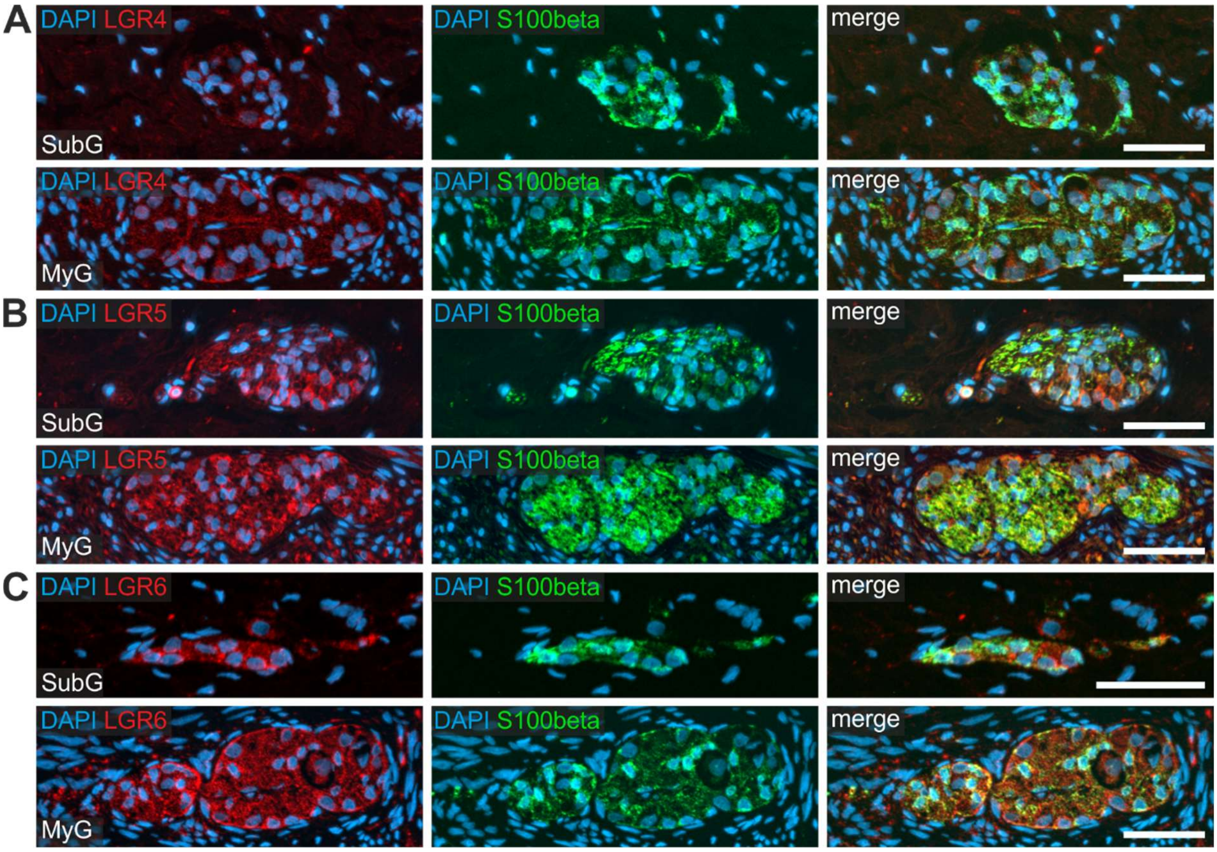


**Suppl. Fig 6: LGR5- and LGR6-receptor-expression in enteric glial cells of human small intestine.** Small intestinal resectates of pediatric patients underwent immunofluorescence co-labeling studies with the glia marker S100beta (green) and in **A:** LGR4 (red), **B:** LGR5 (red) and in **C:** LGR6 (red), and the nuclear marker DAPI (blue). In contrast to LGR4 staining intensity, LGR5- and LGR6-intensities were more intense in submucosal (SubG) and myenteric (MyG) ganglia. Thus, also glial cells express RSPO1-relevant LGR-receptors *in vivo*. Related to Figure 4. **Scale bars: 50 µm.**

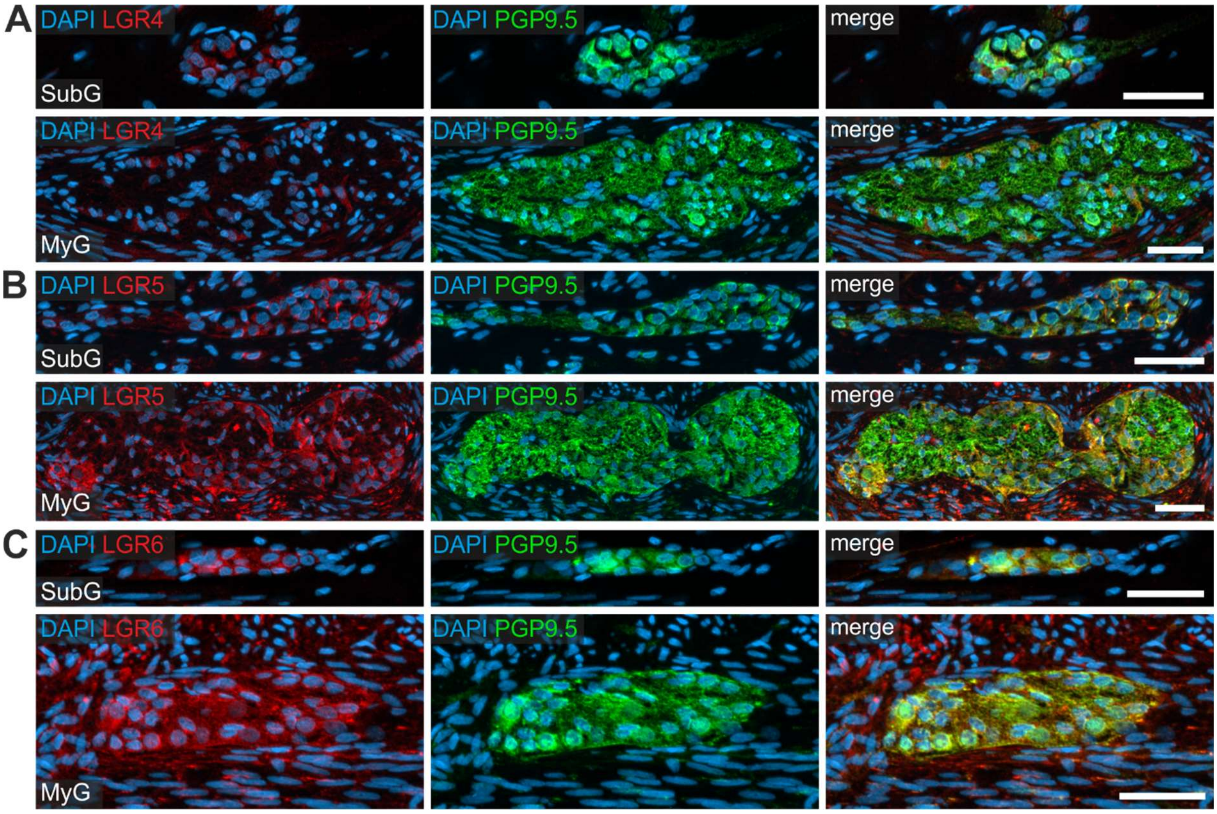


**Suppl. Fig 7: LGR5- and LGR6-receptors are expressed mainly on differentiated enteric neurons of the large intestine.** Large intestinal resectates of pediatric patients underwent immunofluorescence co-labeling studies with the neuronal marker PGP9.5 (green) and in **A:** LGR4 (red), **B:** LGR5 (red) and in **C:** LGR6 (red), and the nuclear marker DAPI (blue). Similar to the small intestine, LGR5- and LGR6-expression was predominantly found in enteric neurons of submucosal (SubG) and myenteric (MyG) ganglia, whereas little to no immunoreactivity was observed for LGR4-reeptor. Related to Figure 4. **Scale bars: 50 µm.**

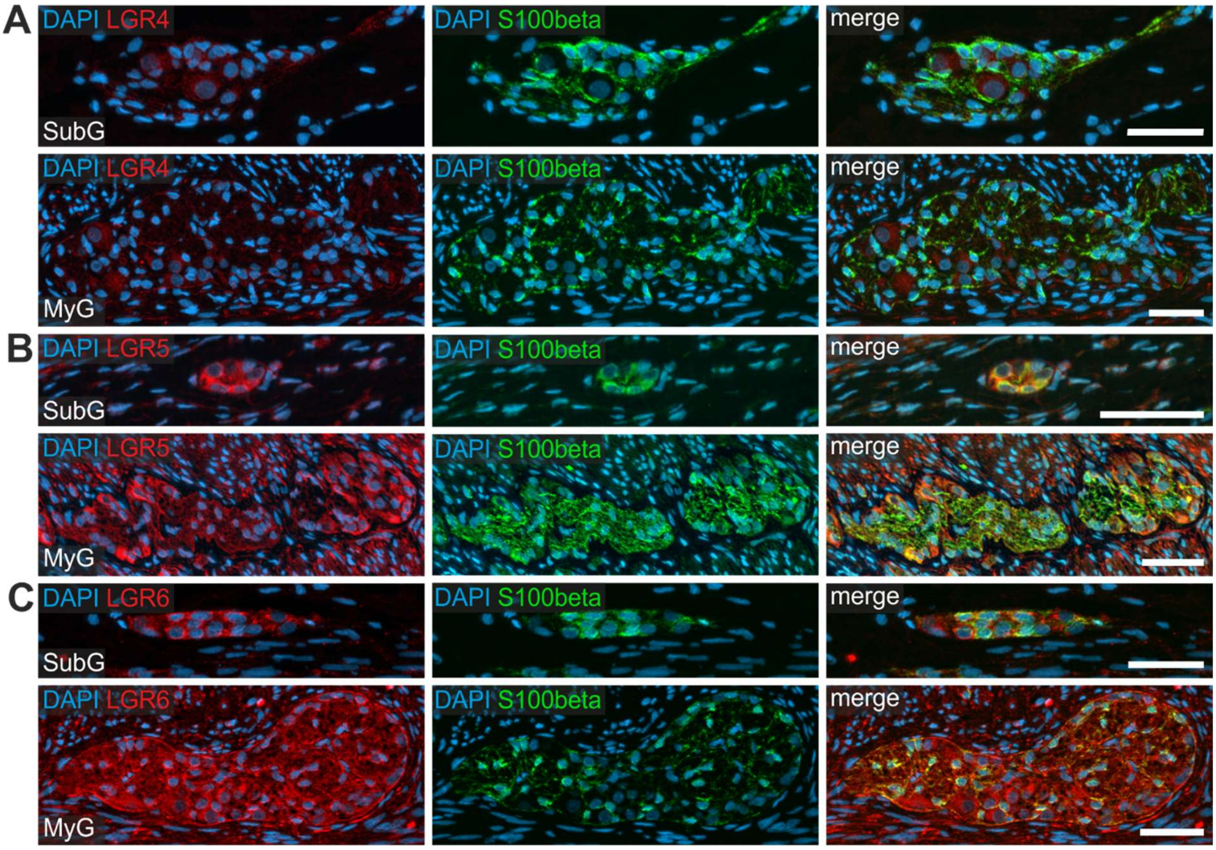


**Suppl. Fig. 8: Enteric glia cells in human large intestine express RSPO1-relevant LGR-receptors.** Small intestinal resectates of pediatric patients underwent immunofluorescence co-labeling studies with the neuronal marker S100beta (green) and in **A:** LGR4 (red), **B:** LGR5 (red) and in **C:** LGR6 (red), and the nuclear marker DAPI (blue). Similar to small intestinal resectates, LGR5- and LGR6-staining intensities were more intense in submucosal (SubG) and myenteric (MyG) ganglia, compared to the immunoreactivity for LGR4. Again, also enteric glia cells in large intestine express LGR-receptors. Related to Figure 4. **Scale bars: 50 µm.**

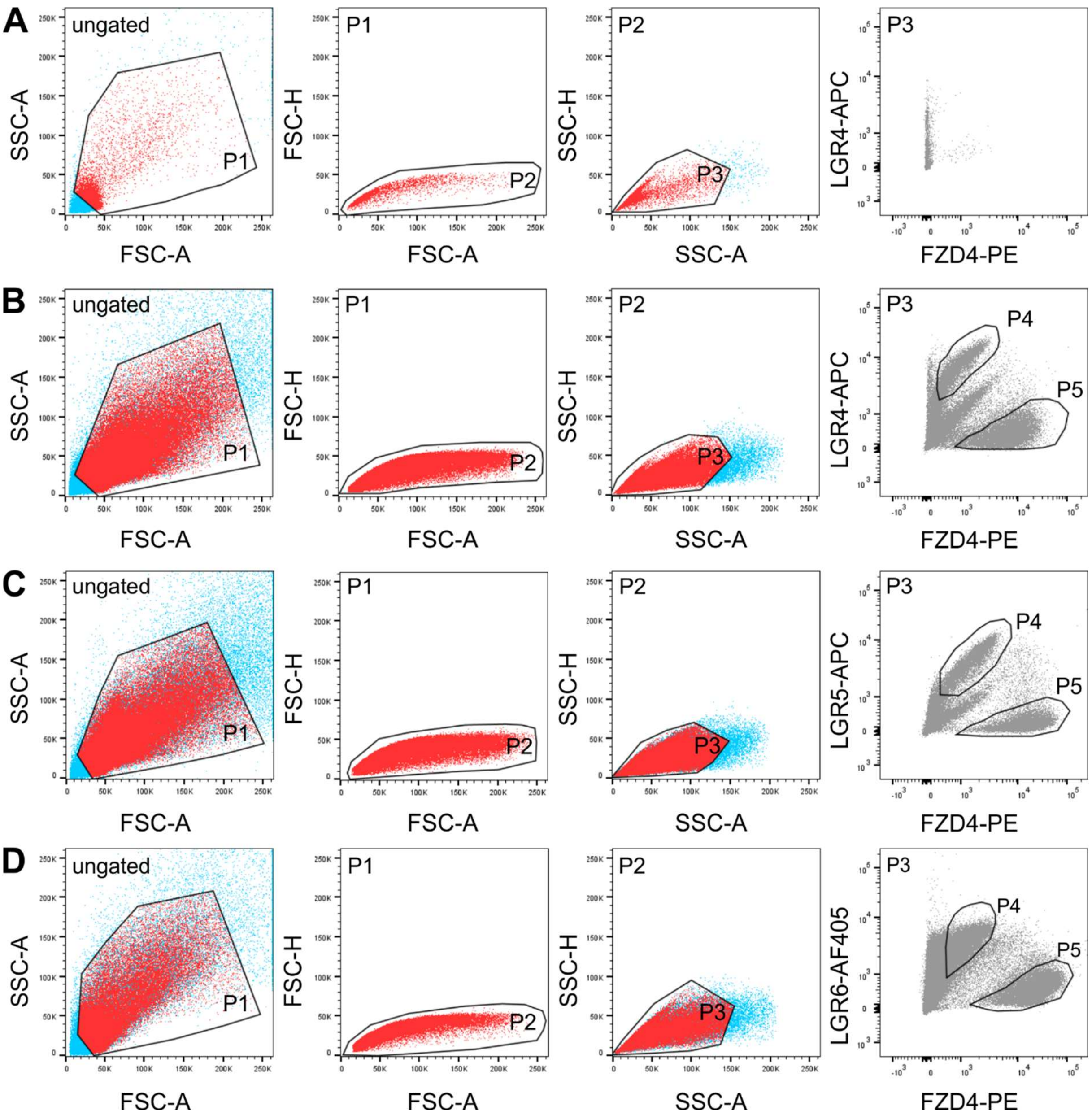


**Suppl. Fig. 9: Fluorescence-activated cell sorting (FACS) of Fzd4 and LGR marked cells derived from human *Tunica muscularis*.** The scatter blots represent the gating strategy (ungated, P1–P3) for Fzd4$^+$LGR$^+$ (P4) and Fzd4$^+$LGR$^-$ (P5) cell pools. **A-D:** depicts one representative experiment for the unstained control **(A)**, as well as the FZD4/LGR4- **(B)**, the FZD4/LGR5- **(C)** and the FZD4/LGR6-co-stained cell population **(D)**. Each population, that underwent further gating are colored in red. All events counted were first gated as P1 to exclude dead cells and debris, as these could be found at the bottom left corner of the first dot blot of each sample. Next, P1 was gated regarding its properties in forward (P2) and sideward (P3) scatter mode to exclude doublets and aggregates. Finally, P3 was gated regarding its FZD4/LGR property. See also the supplementary data file, that summarizes the gating data for all experiments used in this study. Related to Figure 5. Source data are provided as a Source data file.

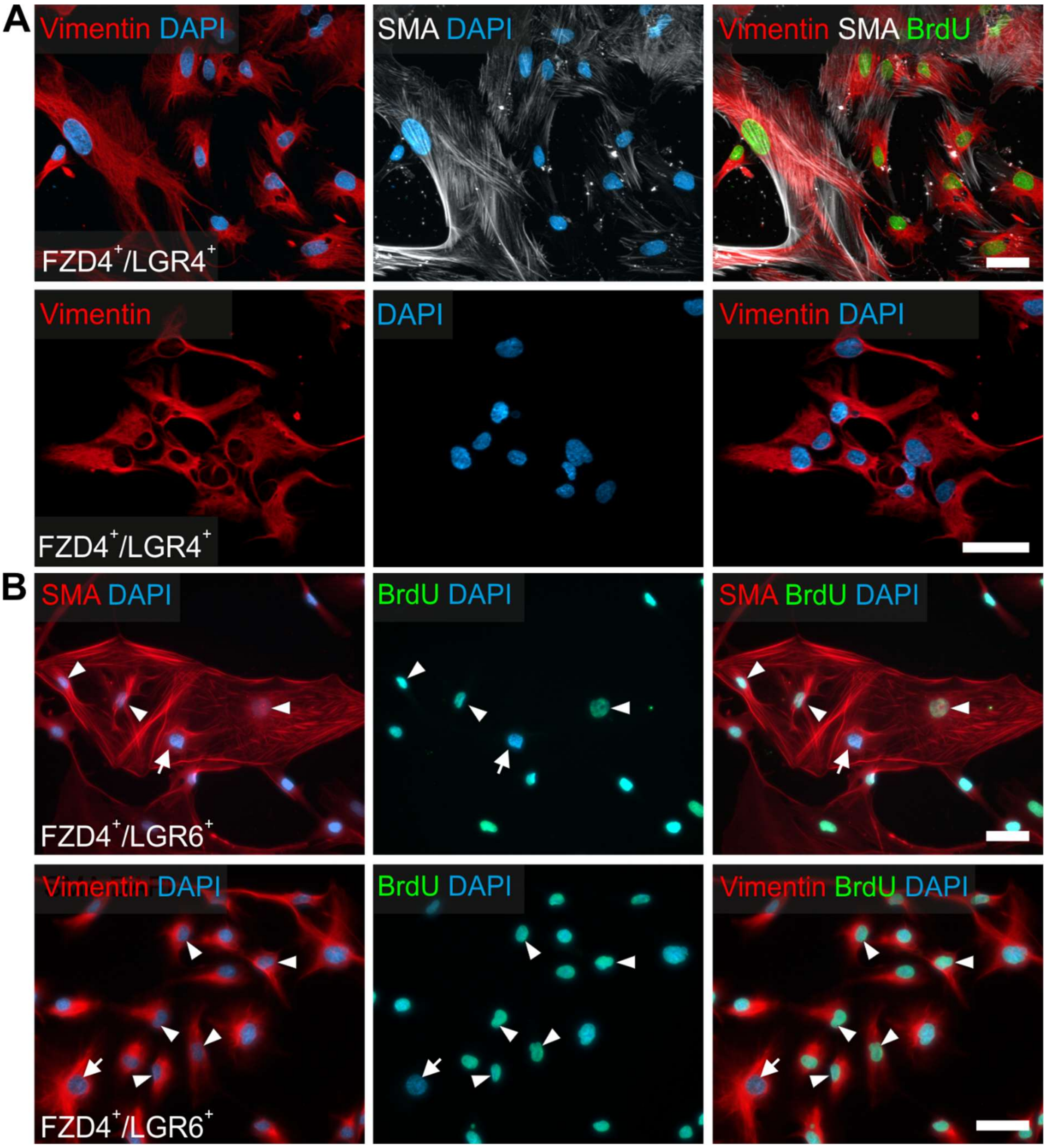


**Suppl. Fig. 10: Mesenchymal cells in FZD4$^+$LGR4$^+$ and FZD4$^+$LGR6$^+$ sorted cell populations.** Immunofluorescence stainings for the fibroblast (vimentin) and smooth muscle cell marker (SMA) in FZD4$^+$LGR4$^+$ (A) and FZD4$^+$LGR6$^+$ cells (B) after differentiation. BrdU-incorporation is shown in green; nuclei are stained using DAPI (blue). FZD4$^+$LGR4$^+$- as well as FZD4$^+$LGR6$^+$-sorted cultures were dominated by BrdU$^+$ (arrowhead) and BrdU$^-$ (arrow) Vimentin$^+$ and SMA$^+$ cells. Related to Figure 6. **Scale bar: 50 µm.**

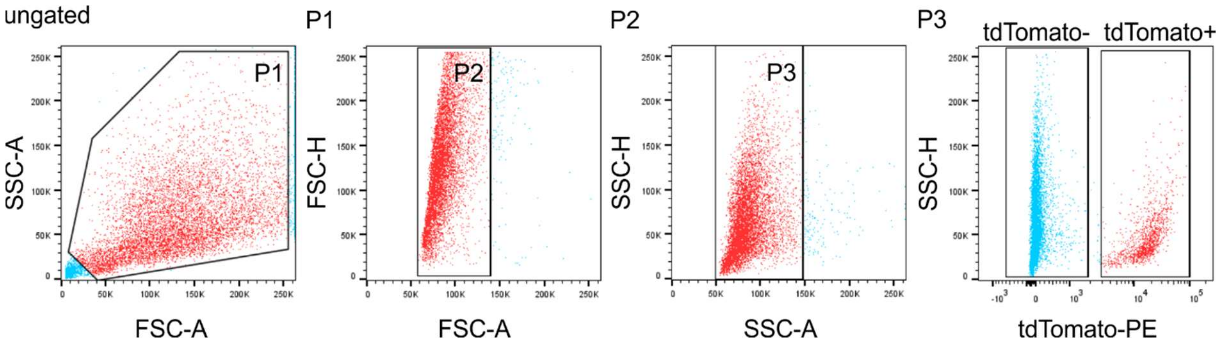


**Suppl. Fig. 11: Fluorescence-activated cell sorting (FACS) of tdTomato expressing neural ENS-cells derived from postnatal day 0 old Wnt1Cre2 mice.** The scatter blots represent the gating strategy (ungated, P1–P3) for tdTomato negative (P4) and tdTomato positive (P5) cell pools. Each population, that underwent further gating are colored in red. All events counted were first gated as P1 to exclude dead cells and debris, as these could be found at the bottom left corner of the first dot blot of each sample. Next, P1 was gated regarding its properties in forward (P2) and sideward (P3) scatter mode to exclude doublets and aggregates. Finally, P3 was gated regarding its tdTomato-expressing property. See also the supplementary data file, that summarizes the gating data for all experiments used in this study. Related to Figure 7. Source data are provided as a Source data file.

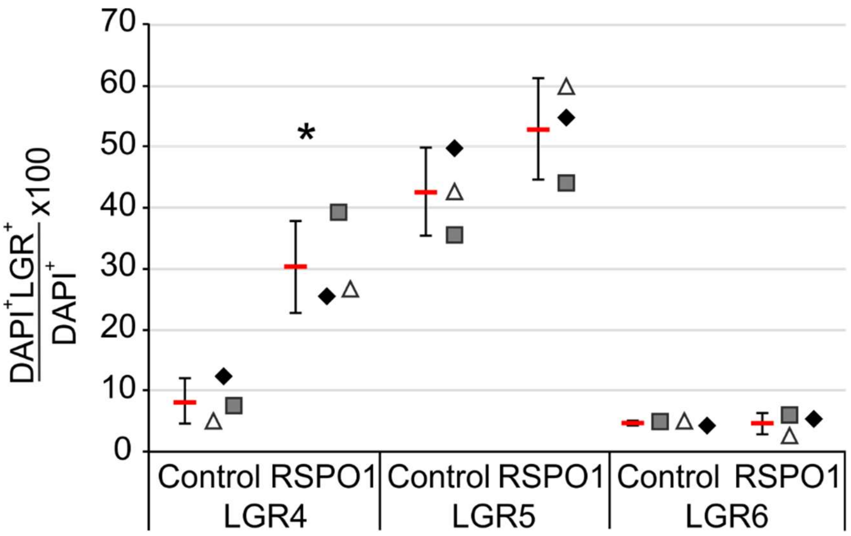


**Suppl. Fig. 12: RSPO1-stimulation increases LGR4- and LGR5-receptor-expression in murine ENS-cells *in vitro*.** RSPO1-treated and unstimulated enterospheres were cultivated under proliferative conditions for 5 div. Fixed-frozen cryosections of these enterospheres were stained for LGR4, LGR5, LGR6, and the nuclear marker DAPI. The bar graph represents the quantification of LGR4$^+$, LGR5$^+$, and LGR6$^+$ cells under control and RSPO1-treated condition. Asterisk indicates significant difference of LGR-expression in RSPO1-stimulated enterospheres compared to control. Data points for independent biological replicates are represented by different symbols. RSPO1-stimulation enhanced significantly the percentage of LGR4$^+$-expressing cells (ANOVA, Fisher LSD post-hoc test, mean±SD: Control: 8.19%±3.72%; RSPO1: 30.3%±7.53%; n=3; P=0.010). The number of LGR5$^+$ cells increased, but not statistically significant (ANOVA, Fisher LSD post-hoc test, mean±SD: Control: 42.6%±7.02%; RSPO1: 52.8%±8.24%; n=3; P=0.179). However, the number of LGR6-expressing cells was not affected by RSPO1-stimulation, arguably due to the small effect size of 3 independent experiments (LGR6: ANOVA, Fisher LSD post-hoc test, mean±SD: Control: 4.64%±0.32%; RSPO1: 4.53%±1.74%; n=3; P=0.918). Related to Figure 7. Source data are provided as a Source data file.

**Supplementary Tables**

**Supplementary Table 1:** Intestinal resectates used in this study. Related to Figures 3-6 and Supplementary Figures 5-10.

| age | sex | Diagnosis | gut region | experiment |
|---|---|---|---|---|
| 3 days | female | imperforate anus | transverse colon | Cell culture |
| 9 months | male | obstruction syndrome | transverse colon | Cell culture |
| 5 months | male | imperforate anus | transverse colon | Cell culture |
| 5 months | female | imperforate anus | transverse colon | Cell culture |
| 3 years / 11 months | female | obstruction syndrome | transverse colon | Fzd4-LGR4 (n1) |
| 12 years / 6 months | female | familial adenomatous polyposis | ileum | Fzd4-LGR4 (n2) |
| 12 years / 3 months | male | obstruction syndrome | transverse colon | Fzd4-LGR4 (n3) |
| 4 months | male | obstruction syndrome | ileum | Fzd4-LGR4 (n4) |
| 2 years / 4 months | female | imperforate anus | ileum | Fzd4-LGR5 (n1) |
| 1 year | male | imperforate anus | ileum | Fzd4-LGR5 (n2) |
| 9 months | female | obstruction syndrome | transverse colon | Fzd4-LGR5 (n3) |
| 1 year / 1 month | male | obstruction syndrome | descendent colon | Fzd4-LGR5 (n4) |
| 2 years / 9 months | male | obstruction syndrome Hirschsprung condition | ileum | Fzd4-LGR5 (n5) |
| 16 years / 4 months | male | obstruction syndrome | ileum | Fzd4-LGR6 (n1) |
| 3 months | male | imperforate anus | ileum | Fzd4-LGR6 (n2) |
| 11 months | male | imperforate anus | transverse colon | Fzd4-LGR6 (n3) |
| 2 years / 1 month | male | obstruction syndrome Hirschsprung condition | ileum | Fzd4-LGR6 (n4) |
| 1 year | female | imperforate anus | transverse colon | Histology |
| 3 months | female | imperforate anus | duodenum | Histology |
| 1 year / 2 months | female | rhabdomyosarcoma | transverse colon | Histology |
| 2 months | male | imperforate anus | descendent colon | Histology |
| 6 months | male | obstruction syndrome | transverse colon | Histology |
| 1 year / 6 months | female | obstruction syndrome | jejunum | Histology |
| 4 months | male | imperforate anus | transverse colon | Histology |
| 7 months | female | imperforate anus | transverse colon | Histology |
| 3 years / 8 months | male | obstruction syndrome | jejunum | Histology |
| 9 months | female | imperforate anus | transverse colon | Histology |
| 4 months | female | imperforate anus | ileum | Histology |
| 11 months | female | obstruction syndrome | ileum | Histology |
| 5 months | male | imperforate anus | descendent colon | Histology |

**Supplementary Table 2:** Expression of LGR receptors in human intestine. Related to Figure 4 and Supplementary Figures 5-8.
(-) no expression, (+) low expression, (++) medium expression, (+++) high expression

| **Small intestine** | | | | | |
|---|---|---|---|---|---|
| **Target Lgr** | submucous plexus (neuron/glia) | myenteric plexus (neuron/glia) | *Tunica muscularis* (circular/longitudinal) | crypts | villus tip |
| **Lgr4** | + (++/+) | + (++/+) | +/+ | + | - |
| **Lgr5** | + (++/+) | + (++/+) | +/+ | + | - |
| **Lgr6** | +++ (+++/+) | +++ (+++/+) | +/+ | - | - |
| **Large intestine** | | | | | |
| **Target Lgr** | submucous plexus (neuron/glia) | myenteric plexus (neuron/glia) | *Tunica muscularis* (circular/longitudinal) | crypts | epithelial surface |
| **Lgr4** | + (++/+) | + (++/+) | +/+ | - | - |
| **Lgr5** | ++ (+++/+) | ++ (+++/+) | +/+ | + | - |
| **Lgr6** | +++ (+++/+) | +++ (+++/+) | +/+ | - | ++ |

### Key Resource Table

| REAGENT/RESOURCES | SOURCE | IDENTIFIER |
|---|---|---|
| **Antibodies** | | |
| anti-rabbit IgG (H+L) AlexaFluor488, dilution: 1:400, host: goat | Invitrogen Thermo Fisher Scientific, MA, USA | Cat. No. A-11008 |
| anti-rabbit IgG (H+L) Alexa Fluor 546, dilution: 1:400, host: goat | Invitrogen Thermo Fisher Scientific, MA, USA | Cat. No. A-11035 |
| anti-rat IgG (H+L) Alexa Fluor 488, dilution: 1:400, host: goat | Invitrogen Thermo Fisher Scientific, MA, USA | Cat. No. A-11006 |
| anti-mouse IgG (H+L) Alexa Fluor 488, dilution: 1:400, host: goat | Invitrogen Thermo Fisher Scientific, MA, USA | Cat. No. A-10680 |
| anti-mouse IgG (H+L) Alexa Fluor 546, dilution: 1:400, host: goat | Invitrogen Thermo Fisher Scientific, MA, USA | Cat. No. A-11030 |
| anti-mouse IgG (H+L) Alexa Fluor 647, dilution: 1:400, host: goat | Invitrogen Thermo Fisher Scientific, MA, USA | Cat. No. A-21235 |
| BrdU, dilution: 1:100, host: rat | Abcam, Cambridge, UK | Cat. No. ab6326 |
| Fzd4-PE (CD344), dilution: 5μl/$10^6$ cells host: mouse, clone: CH3A4A7, Em.: 488 nm, Ex.: 565-605 nm | Bio Legend, CA, USA | Cat. No. 326606 |
| GFAP, dilution: 1:400, host: rabbit | DAKO, Glostrup, Denmark | Cat. No. 0034 |
| HuC/D, dilution: 1:50, host: mouse | Invitrogen Thermo Fisher Scientific, MA, USA | Cat. No. A21271 |
| Ki67, dilution: 1:100, host: mouse | DCS Innovative Diagnostics, Hamburg, Germany | Cat. No. KI68IC002 |
| Ki67, dilution: 1:50, host: mouse | NOVO Castra Leica Biosystems, IL, USA | Cat. No. NCL-MM1 |
| Lgr4-conjugated APC (GPR48), dilution: 10μl/$10^6$ cells, host: mouse, clone: #852229, Em.: 620-650 nm, Ex.: 660-670 nm | R&D Systems, Inc., MN, USA | Cat. No. FAB7750P |
| Lgr4, dilution: 1:50, host: mouse | Invitrogen Thermo Fisher Scientific, MA, USA | Cat. No. SAB370154 |

| | | |
|---|---|---|
| Lgr4, dilution: 1:50,<br>host: mouse | Santa Cruz,<br>Biotechnology, TX, USA | Cat. No. sc-390630 |
| Lgr5-conjugated APC, dilution: 10µl/$10^6$ cells, host: mouse,<br>clone: #707042, Em.: 620-650 nm, Ex.: 660-670 nm | R&D Systems, Inc., MN, USA | Cat. No. FAB8078A |
| Lgr5, dilution: 1:50,<br>host: mouse | Invitrogen Thermo Fisher Scientific, MA, USA | Cat. No. MA5-25644 |
| Lgr6-conjugated AF405,<br>dilution: 10µl/$10^6$ cells, host: mouse, clone: #918726,<br>Em.: 405 nm, Ex.: 421 nm | R&D Systems, Inc., MN, USA | Cat. No. FAB84581V |
| Lgr6, dilution: 1:50,<br>host: rabbit | Invitrogen Thermo Fisher Scientific, MA, USA | Cat. No. PA5/109909 |
| P75, dilution: 1:400,<br>host: rabbit | Millipore Merck,<br>Darmstadt, Germany | Cat. No. AB1554 |
| S100β, dilution: 1:100,<br>host: rabbit | Abcam, Cambridge, UK | Cat. No. ab52642 |
| SMA, dilution: 1:50,<br>host: mouse | DAKO, Glostrup, Denmark | Cat. No. M0851 |
| SOX10, dilution: 1:50,<br>host: mouse | Novus Biologicals, CO, USA | Cat. No. NBP2-59050 |
| Beta-3-Tub, dilution: 1:4000,<br>host: rabbit | Bio Legend, CA, USA | Cat. No. 802001 |
| Vimentin, dilution: 1:100,<br>host: rabbit | Abcam, Cambridge, UK | Cat. No. ab45939 |
| **Biological samples** | | |
| Mice strain C57BL/6J | Jackson Laboratory, Bar Harbor, ME, USA | Stock No. 000664 |
| Mice strain: B6;129S6-Gt(ROSA)26Sortm9(CAG-tdTomato)Hze/J | Jackson Laboratory, Bar Harbor, ME, USA | Stock No. 007914 |
| Mice strain: B6.Cg-Tg(Wnt1-cre)2Sor/J | Jackson Laboratory, Bar Harbor, ME, USA | Stock No. 022501 |

| | | |
|---|---|---|
| Human gut samples,<br>Patient-specific data are summarized in Supplementary Table 1 | Department of Pediatric Surgery, University Children's Hospital Tübingen, Germany | Project Nr. of the local ethical committee 652/2019BO2 and 066/2023BO2 |
| **Chemicals, peptides, and recombinant proteins** | | |
| Ascorbic Acid-2-phosphate, 1 M | Sigma-Aldrich, Taufkirchen, Germany | Cat. No. A-8960-5G |
| B27 Supplement, 50x | gibco® Thermo Fisher Scientific, MA, USA | Cat. No. 17504-044 |
| Bovine serum albumin, 100x | Roth, Karlsruhe, Germany | Cat. No. 031166075 |
| BrdU (5-bromo-2'-deoxyuridine), 10 mM | Roche Diagnostics, Mannheim, Germany | Cat. No. 11299964001 |
| Cibrofloxacin Kabi, 2000 µg/ml | Fresenius Kabi, Bad Homburg, Germany | nA |
| Citric acid monohydrate, 210,14 g/mol | Roth, Karlsruhe, Germany | Cat. No. 5110.2 |
| Collagen-type-I, rat tail, 1 µg/ml | BD Bioscience, Heidelberg, Germany | Cat. No. 354234 |
| Collagenase type XI, 1926 U/mg | Sigma-Aldrich, Taufkirchen, Germany | Cat. No. C9407 |
| DAPI (4',6-diamidino-2-phenylindolestain), 200 ng/ml | Roth, Karlsruhe, Germany | Cat. No. 6335.1 |
| Dispase type II, 1.01 U/mg | Roche Diagnostics, Mannheim, Germany | Cat. No. D4693 |
| Di-Natriumtetraborat-10-hydrat, 201.22 g/mol | Merck, Darmstadt Germany | Cat. No. 0024611 |
| DMEM (Dulbecco's modified Eagle's medium with Ham's F12 medium 1:1), 1x | Life technologies, Darmstadt, Germany | Cat. No. 21331-020 |
| DNase I, 5% | Sigma-Aldrich, Taufkirchen, Germany | Cat. No. DN25 |
| hEGF (human epidermal growth factor), 40 µg/ml | Sigma-Aldrich, Taufkirchen, Germany | Cat. No. E9644 |
| FCS (Fetal calf serum), 100x | Biochrom, Berlin, Germany | Cat. No. S0613 |
| hbFGF (human bone fibroblast-like growth factor), 40 µg/ml | Sigma-Aldrich, Taufkirchen, Germany | Cat. No. F0291 |

| | | |
|---|---|---|
| Gamunex, 100 mg/ml | Talecris Biothera-peutics, NY, USA | Cat. No. 80A2828 |
| Goat serum, 100x | Biochrom, Berlin, Germany | Cat. No. 57288 |
| HBSS (Hanks' balanced salt solution without $Ca^{2+}$ / $Mg^{2+}$), 1x | Sigma-Aldrich, Taufkirchen, Germany | Cat. No. H9394 |
| HBSS (Hanks' balanced salt solution with $Ca^{2+}$ / $Mg^{2+}$), 1x | Sigma-Aldrich, Taufkirchen, Germany | Cat. No. 55037C |
| HCl (hydrochloric acid), 2N | VWR® Chemicals, Darmstadt, Germany | Cat. No.: 310701.5000 |
| Hibernate®-A | gibco® Thermo Fisher Scientific, MA, USA | Cat. No. A12475-01 |
| Kaiser's glycerol gelatine | Merck, Darmstadt, Germany | Cat. No. 109242 |
| L-glutamine, 200 mM | Sigma-Aldrich, Taufkirchen, Germany | Cat. No. G7513 |
| Metronidazole, 5000 µg/ml | B. Braun, Melsungen, Germany | nA |
| N2 supplement, 100x | Life Technologies, Darmstadt, Germany | Cat. No 17502-048 |
| PFA (Paraformaldehyde), 30.39 g/mol | Merck KGaA, Darmstadt, Germany | Cat. No. 1.04005.1000 |
| Penicillin/Streptomycin, 10.000 U/ml/10 mg/ml | Sigma-Aldrich, Taufkirchen, Germany | Cat. No. P0781 |
| Rock Inhibitor, 50 mg | Selleck Chemicals GmbH, Köln, Germany | Cat. No. #688000 |
| R-Spondin1 recombinant mouse, 25 µg | R&D Systems, Inc., MN, USA | Cat. No. 3474-RS |
| R-Spondin1 recombinant human, 25 µg | R&D Systems, Inc., MN, USA | Cat. No. 4645-RS |
| Sucrose (D(+)-Saccharose) | PanReac AppliChem, Darmstadt, Germany | Cat. No. A2211.5000 |
| TissueTek® | Sakura, Staufen, Germany | Cat. No. 4583 |
| Triton® X-100 | Roth, Karlsruhe, Germany | Cat. No. 3051.4 |
| WNT3A recombinant mouse, 2 µg | R&D Systems, Inc., MN, USA | Cat. No. 1324-WNT |

| | | |
|---|---|---|
| WNT3A recombinant human, 10 µg | R&D Systems, Inc., MN, USA | Cat. No. 5036-WN |
| **Critical commercial assays** | | |
| RNeasy Plus Mini Kit | Qiagen, Hilden, Germany | Cat. No. 74104 |
| QuantiTect Reverse Transcription Kit | Qiagen, Hilden, Germany | Cat. No. 205311 |
| PerfeCTa qPCR ToughMix ROX | Quantabio, Berverly, MA, USA | Cat. No. 733-2093 |
| **Oligonucleotides** | | |
| Rspo1, mouse<br>fwd 5´-cgacatgaacaaatgcatca-3´<br>rev 5`-ctcctgacacttggtgcaga-3´ | Roche Diagnostics, Mannheim, Germany | Assay ID: 310599 |
| Rspo2, mouse<br>fwd 5´-gtccaggagatgcaagatgg-3´<br>rev 5`-tcttttgcctttggtgttctc-3´ | Roche Diagnostics, Mannheim, Germany | Assay ID: 310663 |
| Rspo3, mouse<br>fwd 5´-tcaaagggagagcgagga-3´ rev 5`-cagaggaggagcttgtttcc-3´ | Roche Diagnostics, Mannheim, Germany | Assay ID: 310540 |
| Rspo4, mouse<br>fwd 5´-agagactctgcccaggagaa-3´<br>rev 5`-ccaacttcctgtccttacgc-3´ | Roche Diagnostics, Mannheim, Germany | Assay ID: 310519 |
| Lgr4, mouse | Invitrogen Thermo Fisher Scientific, MA, USA | Mm00554385 |
| Lgr5, mouse | Invitrogen Thermo Fisher Scientific, MA, USA | Mm00438890 |
| Lgr6, mouse | Invitrogen Thermo Fisher Scientific, MA, USA | Mm01291336 |
| Rnf43, mouse | Invitrogen Thermo Fisher Scientific, MA, USA | Mm00552558 |
| Znf3, mouse | Invitrogen Thermo Fisher Scientific, MA, USA | Mm01191453 |
| Lrp5, mouse<br>fwd 5´-catggacatccaagtgctga-3´<br>rev 5`-ttgtcctcctcgcatggt-3´ | Roche Diagnostics, Mannheim, Germany | Assay ID: 310507 |
| Lrp6, mouse<br>fwd 5´-tcctcgagctctggcact-3´<br>rev 5`-cctccccactcagtccaata-3´ | Roche Diagnostics, Mannheim, Germany | Assay ID: 310507 |

| | | |
|---|---|---|
| Gapdh, mouse<br>fwd 5´-agcttgtcatcaacgggaag-3´<br>rev 5`-tttgatgttagtggggtctcg-3´ | Roche Diagnostics, Mannheim, Germany | Assay ID: 307884 |
| Hprt, mouse<br>fwd 5´-tcctcctcagaccgctttt-3´<br>rev 5`-cctggttcatcatcgctaatc-3´ | Roche Diagnostics, Mannheim, Germany | Assay ID: 307879 |
| Tbp, mouse<br>fwd 5`-ggcggtttggctaggttt -3´<br>rev 5`-gggttatcttcacacaccatga -3´<br>rev 5`-acccgtgatgggataaacag-3´ | Roche Diagnostics, Mannheim, Germany | Assay ID: RN01455648 |
| **Software and algorithms** | | |
| Axiovision software | Zeiss, Oberkochen, Germany | nA |
| SigmaStat 3.5 software | Systat Software GmbH, Frankfurt, Germany | nA |
| **Other** | | |
| Axio Imager.Z1 | Zeiss, Oberkochen, Germany | nA |
| BD FACS Aria flow cytometer | BD Biosciences, Heidelberg, Germany | nA |
| McIlwain tissue chopper | Mickle Laboratory Engineering Co, Guildford, UK). | nA |
| QIAxcel Advanced | Qiagen, Hilden, Germany | nA |
| StepOnePlus™ Real-Time PCR System | Applied Biosystems, Darmstadt, Germany | nA |
