## Supplementary material for "R-Spondin1 regulates fate of enteric neural progenitors via differential LGR4/5/6-expression in mice and humans": Source data file

**Figure 1C**

| counted sphere number | Control | RSPO1 100 ng/ml | Wnt3a 20 ng/ml | RSPO1 100 ng/ml Wnt3a 20 ng/ml |
|---|---|---|---|---|
| n1 | 452 | 733 | 814 | 754 |
| n2 | 399 | 683 | 540 | 557 |
| n3 | 442 | 786 | 620 | 672 |

| relative values of sphere numbers | Control | RSPO1 100 ng/ml | Wnt3a 20 ng/ml | RSPO1 100 ng/ml Wnt3a 20 ng/ml |
|---|---|---|---|---|
| n1 | 1,00 | 1,62 | 1,80 | 1,67 |
| n2 | 1,00 | 1,71 | 1,35 | 1,40 |
| n3 | 1,00 | 1,78 | 1,40 | 1,52 |
| mean | 1,00 | 1,70 | 1,52 | 1,53 |
| SD | 0,00 | 0,08 | 0,25 | 0,14 |
| P-VALUE | - | 0.005 | 0.005 | 0.003 |

**Statistics**

| sphere number | DEGREES OF FREEDOM | F-VALUE |
|---|---|---|
| Between groups | 3 | 13.055 |
| Residuals | 8 | |
| Total | 11 | |

| Comparison | | | Diff of means | LSD(alpha=0,050) | P | Diff |
|---|---|---|---|---|---|---|
| Wnt3a 20 ng/ml | vs. | Control | 0,704 | 0,274 | <0,001 | Yes |
| Wnt3a 20 ng/ml | vs. | RSPO1 100 ng/ml | 0,185 | 0,274 | 0,159 | No |
| Wnt3a 20 ng/ml | vs. | Wnt3a 20 ng/ml + RSPO1 100 ng/ml | 0,176 | 0,274 | 0,178 | Do Not Test |
| Wnt3a 20 ng/ml + RSPO1 100 ng/ml | vs. | Control | 0,528 | 0,274 | 0,002 | Yes |
| Wnt3a 20 ng/ml + RSPO1 100 ng/ml | vs. | RSPO1 100 ng/ml | 0,00917 | 0,274 | 0,94 | Do Not Test |
| RSPO1 100 ng/ml | vs. | Control | 0,519 | 0,274 | 0,002 | Yes |

| summed up cell volume in µm³ | Control | RSPO1 100 ng/ml | Wnt3a 20 ng/ml | RSPO1 100 ng/ml Wnt3a 20 ng/ml |
|---|---|---|---|---|
| n1 | 25477761 | 61847660 | 57451010 | 65302075 |
| n2 | 17060362 | 27569185 | 32439163 | 35403803 |
| n3 | 119901980 | 343972631 | 327771506 | 329154986 |

| relative values of sphere volume | Control | RSPO1 100 ng/ml | Wnt3a 20 ng/ml | RSPO1 100 ng/ml Wnt3a 20 ng/ml |
|---|---|---|---|---|
| n1 | 1,00 | 2,43 | 2,25 | 2,56 |
| n2 | 1,00 | 1,62 | 1,90 | 2,08 |
| n3 | 1,00 | 2,87 | 2,73 | 2,75 |
| mean | 1,00 | 2,30 | 2,30 | 2,46 |
| SD | 0,00 | 0,64 | 0,42 | 0,35 |
| P-VALUE | - | 0.002 | <0.001 | 0.002 |

**Statistics**

| sphere volume | DEGREES OF FREEDOM | F-VALUE |
|---|---|---|
| Between groups | 3 | 7.975 |
| Residuals | 8 | |
| Total | 11 | |

| Comparison | | | Diff of means | LSD(alpha=0,050) | P | Diff |
|---|---|---|---|---|---|---|
| Wnt3a 20 ng/ml + RSPO1 100 ng/ml | vs. | Control | 1,461 | 0,787 | 0,003 | Yes |
| Wnt3a 20 ng/ml + RSPO1 100 ng/ml | vs. | RSPO1 100 ng/ml | 0,164 | 0,787 | 0,643 | No |
| Wnt3a 20 ng/ml + RSPO1 100 ng/ml | vs. | Wnt3a 20 ng/ml | 0,157 | 0,787 | 0,657 | Do Not Test |
| Wnt3a 20 ng/ml | vs. | Control | 1,304 | 0,787 | 0,005 | Yes |
| Wnt3a 20 ng/ml | vs. | RSPO1 100 ng/ml | 0,00741 | 0,787 | 0,983 | Do Not Test |
| RSPO1 100 ng/ml | vs. | Control | 1,297 | 0,787 | 0,005 | Yes |

**Figure 1E**

| experimental groups | HuC/D+ cells counted per image | mean values of four technical replicates per group | mean values of all experiments | SD | P-VALUE |
|---|---|---|---|---|---|
| Control_n1 | 901 | | | | |
| Control_n1 | 1835 | | | | |
| Control_n1 | 1406 | | | | |
| Control_n1 | 1963 | 6105 | | | |
| Control_n2 | 1570 | | | | |
| Control_n2 | 2031 | | | | |
| Control_n2 | 2005 | | | | |
| Control_n2 | 1607 | 7213 | | | |
| Control_n3 | 2180 | | | | |
| Control_n3 | 1895 | | | | |
| Control_n3 | 825 | | | | |
| Control_n3 | 927 | 5827 | 6382 | 733 | - |
| Wnt3a 20 ng/ml_n1 | 1843 | | | | |
| Wnt3a 20 ng/ml_n1 | 3708 | | | | |
| Wnt3a 20 ng/ml_n1 | 1170 | | | | |
| Wnt3a 20 ng/ml_n1 | 3960 | 10681 | | | |
| Wnt3a 20 ng/ml_n2 | 2300 | | | | |
| Wnt3a 20 ng/ml_n2 | 2738 | | | | |
| Wnt3a 20 ng/ml_n2 | 3363 | | | | |
| Wnt3a 20 ng/ml_n2 | 2168 | 10569 | | | |
| Wnt3a 20 ng/ml_n3 | 2068 | | | | |
| Wnt3a 20 ng/ml_n3 | 2445 | | | | |
| Wnt3a 20 ng/ml_n3 | 2427 | | | | |
| Wnt3a 20 ng/ml_n3 | 2498 | 9438 | 10229 | 688 | ≤0.001 |
| RSPO1 100 ng/ml_n1 | 884 | | | | |
| RSPO1 100 ng/ml_n1 | 4241 | | | | |
| RSPO1 100 ng/ml_n1 | 1606 | | | | |
| RSPO1 100 ng/ml_n1 | 4535 | 11266 | | | |
| RSPO1 100 ng/ml_n2 | 2397 | | | | |
| RSPO1 100 ng/ml_n2 | 2239 | | | | |
| RSPO1 100 ng/ml_n2 | 3093 | | | | |
| RSPO1 100 ng/ml_n2 | 2101 | 9830 | | | |
| RSPO1 100 ng/ml_n3 | 1640 | | | | |
| RSPO1 100 ng/ml_n3 | 1916 | | | | |
| RSPO1 100 ng/ml_n3 | 2145 | | | | |
| RSPO1 100 ng/ml_n3 | 3555 | 9256 | 10117 | 1035 | ≤0.001 |
| Wnt3a 20 ng/ml + RSPO1 100 ng/ml_n1 | 839 | | | | |
| Wnt3a 20 ng/ml + RSPO1 100 ng/ml_n1 | 3227 | | | | |
| Wnt3a 20 ng/ml + RSPO1 100 ng/ml_n1 | 899 | | | | |
| Wnt3a 20 ng/ml + RSPO1 100 ng/ml_n1 | 4948 | 9913 | | | |
| Wnt3a 20 ng/ml + RSPO1 100 ng/ml_n2 | 1933 | | | | |
| Wnt3a 20 ng/ml + RSPO1 100 ng/ml_n2 | 2680 | | | | |
| Wnt3a 20 ng/ml + RSPO1 100 ng/ml_n2 | 2615 | | | | |
| Wnt3a 20 ng/ml + RSPO1 100 ng/ml_n2 | 1994 | 9222 | | | |
| Wnt3a 20 ng/ml + RSPO1 100 ng/ml_n3 | 2021 | | | | |
| Wnt3a 20 ng/ml + RSPO1 100 ng/ml_n3 | 2063 | | | | |
| Wnt3a 20 ng/ml + RSPO1 100 ng/ml_n3 | 2437 | | | | |
| Wnt3a 20 ng/ml + RSPO1 100 ng/ml_n3 | 2004 | 8525 | 9220 | 694 | 0.002 |

**Statistics**

| HuCD | DEGREES OF FREEDOM | F-VALUE |
|---|---|---|
| Between groups | 3 | 15.075 |
| Residuals | 8 | |
| Total | 11 | |

| Comparison | | | Diff of means | LSD(alpha=0,050) | P | Diff |
|---|---|---|---|---|---|---|
| Wnt3a 20 ng/ml | vs. | Control | 3847,667 | 1507,462 | <0,001 | Yes |
| Wnt3a 20 ng/ml | vs. | Wnt3a 20 ng/ml + RSPO1 100 ng/ml | 1009,333 | 1507,462 | 0,161 | No |
| Wnt3a 20 ng/ml | vs. | RSPO1 100 ng/ml | 112 | 1507,462 | 0,868 | Do Not Test |
| RSPO1 100 ng/ml | vs. | Control | 3735,667 | 1507,462 | <0,001 | Yes |
| RSPO1 100 ng/ml | vs. | Wnt3a 20 ng/ml + RSPO1 100 ng/ml | 897,333 | 1507,462 | 0,207 | Do Not Test |
| Wnt3a 20 ng/ml + RSPO1 100 ng/ml | vs. | Control | 2838,333 | 1507,462 | 0,002 | Yes |

**Figure 1F**

| experimental groups | HuC/D+ cells counted per image | HuC/D+ and BrdU+ cells counted per image | total amount of HuC/D+ cells per group | total amount of HuC/D+ and BrdU+ cells per group | percentage of HuC/D+ and BrdU+ cells | mean | SD | P-VALUE |
|---|---|---|---|---|---|---|---|---|
| Control_n1 | 56 | 13 | | | | | | |
| Control_n1 | 9 | 4 | | | | | | |
| Control_n1 | 26 | 9 | | | | | | |
| Control_n1 | 50 | 17 | | | | | | |
| Control_n1 | 29 | 11 | | | | | | |
| Control_n1 | 16 | 7 | | | | | | |
| Control_n1 | 30 | 8 | | | | | | |
| Control_n1 | 40 | 5 | 256 | 74 | 29% | | | |
| Control_n2 | 31 | 9 | | | | | | |
| Control_n2 | 14 | 4 | | | | | | |
| Control_n2 | 13 | 5 | | | | | | |
| Control_n2 | 21 | 5 | | | | | | |
| Control_n2 | 29 | 9 | | | | | | |
| Control_n2 | 18 | 7 | | | | | | |
| Control_n2 | 17 | 3 | | | | | | |
| Control_n2 | 28 | 6 | 171 | 48 | 28% | | | |
| Control_n3 | 81 | 12 | | | | | | |
| Control_n3 | 105 | 20 | | | | | | |
| Control_n3 | 73 | 12 | | | | | | |
| Control_n3 | 39 | 17 | | | | | | |
| Control_n3 | 33 | 8 | | | | | | |
| Control_n3 | 49 | 24 | | | | | | |
| Control_n3 | 64 | 17 | | | | | | |
| Control_n3 | 53 | 13 | 497 | 123 | 25% | 27% | 2% | - |
| Wnt3a 20 ng/ml_n1 | 34 | 14 | | | | | | |
| Wnt3a 20 ng/ml_n1 | 27 | 13 | | | | | | |
| Wnt3a 20 ng/ml_n1 | 53 | 21 | | | | | | |
| Wnt3a 20 ng/ml_n1 | 33 | 13 | | | | | | |
| Wnt3a 20 ng/ml_n1 | 49 | 22 | | | | | | |
| Wnt3a 20 ng/ml_n1 | 59 | 28 | | | | | | |
| Wnt3a 20 ng/ml_n1 | 18 | 8 | | | | | | |
| Wnt3a 20 ng/ml_n1 | 36 | 13 | 309 | 132 | 43% | | | |
| Wnt3a 20 ng/ml_n2 | 52 | 22 | | | | | | |
| Wnt3a 20 ng/ml_n2 | 22 | 10 | | | | | | |
| Wnt3a 20 ng/ml_n2 | 28 | 10 | | | | | | |
| Wnt3a 20 ng/ml_n2 | 26 | 14 | | | | | | |
| Wnt3a 20 ng/ml_n2 | 23 | 12 | | | | | | |
| Wnt3a 20 ng/ml_n2 | 51 | 25 | | | | | | |
| Wnt3a 20 ng/ml_n2 | 66 | 28 | | | | | | |
| Wnt3a 20 ng/ml_n2 | 23 | 14 | 291 | 135 | 46% | | | |
| Wnt3a 20 ng/ml_n3 | 115 | 46 | | | | | | |
| Wnt3a 20 ng/ml_n3 | 136 | 49 | | | | | | |
| Wnt3a 20 ng/ml_n3 | 116 | 51 | | | | | | |
| Wnt3a 20 ng/ml_n3 | 192 | 67 | | | | | | |
| Wnt3a 20 ng/ml_n3 | 88 | 42 | | | | | | |
| Wnt3a 20 ng/ml_n3 | 158 | 51 | | | | | | |
| Wnt3a 20 ng/ml_n3 | 111 | 45 | | | | | | |
| Wnt3a 20 ng/ml_n3 | 120 | 52 | 1036 | 403 | 39% | 43% | 4% | ≤0.001 |
| RSPO1 100 ng/ml_n1 | 38 | 19 | | | | | | |
| RSPO1 100 ng/ml_n1 | 36 | 17 | | | | | | |
| RSPO1 100 ng/ml_n1 | 27 | 11 | | | | | | |
| RSPO1 100 ng/ml_n1 | 46 | 22 | | | | | | |
| RSPO1 100 ng/ml_n1 | 22 | 9 | | | | | | |
| RSPO1 100 ng/ml_n1 | 27 | 12 | | | | | | |
| RSPO1 100 ng/ml_n1 | 46 | 22 | | | | | | |
| RSPO1 100 ng/ml_n1 | 47 | 23 | 289 | 135 | 47% | | | |
| RSPO1 100 ng/ml_n2 | 29 | 12 | | | | | | |
| RSPO1 100 ng/ml_n2 | 28 | 13 | | | | | | |
| RSPO1 100 ng/ml_n2 | 45 | 21 | | | | | | |
| RSPO1 100 ng/ml_n2 | 34 | 14 | | | | | | |
| RSPO1 100 ng/ml_n2 | 55 | 21 | | | | | | |
| RSPO1 100 ng/ml_n2 | 34 | 12 | | | | | | |
| RSPO1 100 ng/ml_n2 | 21 | 13 | | | | | | |
| RSPO1 100 ng/ml_n2 | 35 | 15 | 281 | 121 | 43% | | | |
| RSPO1 100 ng/ml_n3 | 109 | 46 | | | | | | |
| RSPO1 100 ng/ml_n3 | 114 | 50 | | | | | | |
| RSPO1 100 ng/ml_n3 | 85 | 48 | | | | | | |
| RSPO1 100 ng/ml_n3 | 126 | 52 | | | | | | |
| RSPO1 100 ng/ml_n3 | 155 | 54 | | | | | | |
| RSPO1 100 ng/ml_n3 | 108 | 66 | | | | | | |
| RSPO1 100 ng/ml_n3 | 73 | 34 | | | | | | |
| RSPO1 100 ng/ml_n3 | 206 | 90 | 976 | 440 | 45% | 45% | 2% | ≤0.001 |
| Wnt3a 20 ng/ml + RSPO1 100 ng/ml_n1 | 42 | 16 | | | | | | |
| Wnt3a 20 ng/ml + RSPO1 100 ng/ml_n1 | 25 | 11 | | | | | | |
| Wnt3a 20 ng/ml + RSPO1 100 ng/ml_n1 | 26 | 19 | | | | | | |
| Wnt3a 20 ng/ml + RSPO1 100 ng/ml_n1 | 43 | 25 | | | | | | |
| Wnt3a 20 ng/ml + RSPO1 100 ng/ml_n1 | 27 | 18 | | | | | | |
| Wnt3a 20 ng/ml + RSPO1 100 ng/ml_n1 | 23 | 11 | | | | | | |
| Wnt3a 20 ng/ml + RSPO1 100 ng/ml_n1 | 23 | 11 | | | | | | |
| Wnt3a 20 ng/ml + RSPO1 100 ng/ml_n1 | 36 | 26 | 245 | 137 | 56% | | | |
| Wnt3a 20 ng/ml + RSPO1 100 ng/ml_n2 | 53 | 18 | | | | | | |
| Wnt3a 20 ng/ml + RSPO1 100 ng/ml_n2 | 46 | 23 | | | | | | |
| Wnt3a 20 ng/ml + RSPO1 100 ng/ml_n2 | 30 | 13 | | | | | | |
| Wnt3a 20 ng/ml + RSPO1 100 ng/ml_n2 | 35 | 16 | | | | | | |
| Wnt3a 20 ng/ml + RSPO1 100 ng/ml_n2 | 30 | 16 | | | | | | |
| Wnt3a 20 ng/ml + RSPO1 100 ng/ml_n2 | 36 | 26 | | | | | | |
| Wnt3a 20 ng/ml + RSPO1 100 ng/ml_n2 | 44 | 25 | | | | | | |
| Wnt3a 20 ng/ml + RSPO1 100 ng/ml_n2 | 44 | 19 | 318 | 156 | 49% | | | |
| Wnt3a 20 ng/ml + RSPO1 100 ng/ml_n3 | 75 | 30 | | | | | | |
| Wnt3a 20 ng/ml + RSPO1 100 ng/ml_n3 | 81 | 41 | | | | | | |
| Wnt3a 20 ng/ml + RSPO1 100 ng/ml_n3 | 90 | 40 | | | | | | |
| Wnt3a 20 ng/ml + RSPO1 100 ng/ml_n3 | 94 | 55 | | | | | | |
| Wnt3a 20 ng/ml + RSPO1 100 ng/ml_n3 | 98 | 54 | | | | | | |
| Wnt3a 20 ng/ml + RSPO1 100 ng/ml_n3 | 91 | 54 | | | | | | |
| Wnt3a 20 ng/ml + RSPO1 100 ng/ml_n3 | 89 | 49 | | | | | | |
| Wnt3a 20 ng/ml + RSPO1 100 ng/ml_n3 | 249 | 110 | 867 | 433 | 50% | 52% | 4% | ≤0.001 |

**Statistics**

| HuCD+BrdU | DEGREES OF FREEDOM | F-VALUE |
|---|---|---|
| Between groups | 3 | 29.579 |
| Residuals | 8 | |
| Total | 11 | |

| Comparison | | | Diff of means | LSD(alpha=0,050) | P | Diff |
|---|---|---|---|---|---|---|
| Wnt3a 20 ng/ml + RSPO1 100 ng/ml | vs. | Control | 0,248 | 0,064 | <0,001 | Yes |
| Wnt3a 20 ng/ml + RSPO1 100 ng/ml | vs. | Wnt3a 20 ng/ml | 0,0936 | 0,064 | 0,01 | Yes |
| Wnt3a 20 ng/ml + RSPO1 100 ng/ml | vs. | RSPO1 100 ng/ml | 0,052 | 0,064 | 0,098 | No |
| RSPO1 100 ng/ml | vs. | Control | 0,196 | 0,064 | <0,001 | Yes |
| RSPO1 100 ng/ml | vs. | Wnt3a 20 ng/ml | 0,0416 | 0,064 | 0,172 | No |
| Wnt3a 20 ng/ml | vs. | Control | 0,154 | 0,064 | <0,001 | Yes |

**Figure 2C**

| spheres evaluated | Control 7 div | Control 14 div | Control diameter difference between 7 div and 14 div in µm | Control relative sphere diameter change | R-Spondin1 100ng/ml 7 div | R-Spondin1 100ng/ml 14 div | R-Spondin1 100ng/ml diameter difference between 7 div and 14 div in µm | R-Spondin1 100ng/ml relative sphere diameter change |
|---|---|---|---|---|---|---|---|---|
| 1 | 112,1 | 155,7 | 43,6 | 1,4 | 78,0 | 276,2 | 198,2 | 3,5 |
| 2 | 113,4 | 166,1 | 52,8 | 1,5 | 114,8 | 185,1 | 70,4 | 1,6 |
| 3 | 78,0 | 87,3 | 9,3 | 1,1 | 99,8 | 101,8 | 1,9 | 1,0 |
| 4 | 75,2 | 86,7 | 11,6 | 1,2 | 117,0 | 131,9 | 14,9 | 1,1 |
| 5 | 78,1 | 89,2 | 11,2 | 1,1 | 99,5 | 147,7 | 48,2 | 1,5 |
| 6 | 96,5 | 111,9 | 15,3 | 1,2 | 101,8 | 158,6 | 56,8 | 1,6 |
| 7 | 106,5 | 144,6 | 38,0 | 1,4 | 48,9 | 160,5 | 111,6 | 3,3 |
| 8 | 113,6 | 145,0 | 31,4 | 1,3 | 88,3 | 206,8 | 118,5 | 2,3 |
| 9 | 121,8 | 124,4 | 2,7 | 1,0 | 190,6 | 371,6 | 181,1 | 2,0 |
| 10 | 114,2 | 116,0 | 1,8 | 1,0 | 76,2 | 151,8 | 75,5 | 2,0 |
| 11 | 90,8 | 102,6 | 11,9 | 1,1 | 84,0 | 86,7 | 2,6 | 1,0 |
| 12 | 41,3 | 44,2 | 2,9 | 1,1 | 129,9 | 344,5 | 214,6 | 2,7 |
| 13 | 36,0 | 55,7 | 19,7 | 1,5 | 97,3 | 310,0 | 212,7 | 3,2 |
| 14 | 86,5 | 110,8 | 24,3 | 1,3 | 134,8 | 190,7 | 55,9 | 1,4 |
| 15 | 101,5 | 105,2 | 3,8 | 1,0 | 160,1 | 349,7 | 189,6 | 2,2 |
| 16 | 80,7 | 92,4 | 11,8 | 1,1 | 122,3 | 180,7 | 58,4 | 1,5 |
| 17 | 47,4 | 56,0 | 8,6 | 1,2 | 275,4 | 577,1 | 301,7 | 2,1 |
| 18 | 85,7 | 108,6 | 22,8 | 1,3 | 207,5 | 498,1 | 290,6 | 2,4 |
| 19 | 117,6 | 188,3 | 70,7 | 1,6 | 145,4 | 453,5 | 308,1 | 3,1 |
| 20 | 60,0 | 67,1 | 7,2 | 1,1 | 111,3 | 229,3 | 118,0 | 2,1 |
| 21 | 89,7 | 100,7 | 11,0 | 1,1 | 96,8 | 144,1 | 47,3 | 1,5 |
| 22 | 80,8 | 55,1 | -25,8 | 0,7 | 119,8 | 199,4 | 79,6 | 1,7 |
| 23 | 89,1 | 68,1 | -21,0 | 0,8 | 148,9 | 279,7 | 130,8 | 1,9 |
| 24 | 88,3 | 84,2 | -4,0 | 1,0 | 142,0 | 256,5 | 114,6 | 1,8 |
| 25 | 84,2 | 66,0 | -18,2 | 0,8 | 124,5 | 158,5 | 34,0 | 1,3 |
| 26 | 62,7 | 64,4 | 1,8 | 1,0 | 116,1 | 140,5 | 24,4 | 1,2 |
| 27 | 81,2 | 68,1 | -13,1 | 0,8 | 111,7 | 121,3 | 9,6 | 1,1 |
| 28 | 69,7 | 71,8 | 2,1 | 1,0 | 44,2 | 104,7 | 60,5 | 2,4 |
| 29 | 85,6 | 74,6 | -11,0 | 0,9 | | | | |
| 30 | 98,2 | 100,2 | 2,0 | 1,0 | | | | |
| 31 | 54,5 | 101,0 | 46,6 | 1,9 | | | | |
| 32 | 56,3 | 65,6 | 9,3 | 1,2 | | | | |
| 33 | 51,2 | 71,5 | 20,3 | 1,4 | | | | |
| 34 | 109,9 | 74,2 | -35,7 | 0,7 | | | | |
| 35 | 104,3 | 110,1 | 5,8 | 1,1 | | | | |
| 36 | 95,6 | 22,8 | -72,8 | 0,2 | | | | |
| 37 | 89,7 | 117,8 | 28,1 | 1,3 | | | | |
| 38 | 86,0 | 88,9 | 2,9 | 1,0 | | | | |
| 39 | 101,5 | 120,3 | 18,8 | 1,2 | | | | |
| 40 | 76,3 | 37,8 | -38,5 | 0,5 | | | | |
| 41 | 50,0 | 42,2 | -7,8 | 0,8 | | | | |
| 42 | 80,2 | 59,3 | -20,9 | 0,7 | | | | |
| 43 | 71,7 | 84,2 | 12,5 | 1,2 | | | | |
| 44 | 81,2 | 78,0 | -3,2 | 1,0 | | | | |
| 45 | 106,2 | 77,9 | -28,2 | 0,7 | | | | |
| 46 | 92,3 | 98,5 | 6,2 | 1,1 | | | | |
| 47 | 95,8 | 108,3 | 12,5 | 1,1 | | | | |
| median | | | | 1,1 | | | | 1,8 |
| P-VALUE | | | | | | | | ≤0.001 |

| spheres evaluated | Wnt3a 20ng/ml 7 div | Wnt3a 20ng/ml 14 div | Wnt3a 20ng/ml diameter difference between 7 div and 14 div in µm | Wnt3a 20ng/ml relative sphere diameter change | Combination 7 div | Combination 14 div | Combination diameter difference between 7 div and 14 div in µm | Combination relative sphere diameter change |
|---|---|---|---|---|---|---|---|---|
| 1 | 148,8 | 196,4 | 47,6 | 1,3 | 128,6 | 191,2 | 62,6 | 1,5 |
| 2 | 120,5 | 203,0 | 82,5 | 1,7 | 92,2 | 282,3 | 190,0 | 3,1 |
| 3 | 129,6 | 238,7 | 109,1 | 1,8 | 72,1 | 257,3 | 185,3 | 3,6 |
| 4 | 95,2 | 163,8 | 68,5 | 1,7 | 57,0 | 68,5 | 11,5 | 1,2 |
| 5 | 84,9 | 175,1 | 90,2 | 2,1 | 73,9 | 95,9 | 21,9 | 1,3 |
| 6 | 80,2 | 154,5 | 74,3 | 1,9 | 133,5 | 264,5 | 131,1 | 2,0 |
| 7 | 87,8 | 127,9 | 40,2 | 1,5 | 79,7 | 251,3 | 171,6 | 3,2 |
| 8 | 67,5 | 165,1 | 97,7 | 2,4 | 106,5 | 398,1 | 291,6 | 3,7 |
| 9 | 93,0 | 172,9 | 79,8 | 1,9 | 87,0 | 464,5 | 377,5 | 5,3 |
| 10 | 114,1 | 187,7 | 73,6 | 1,6 | 72,2 | 205,5 | 133,3 | 2,8 |
| 11 | 127,8 | 231,8 | 104,0 | 1,8 | 130,7 | 207,9 | 77,2 | 1,6 |
| 12 | 119,4 | 216,6 | 97,2 | 1,8 | 131,7 | 228,7 | 97,0 | 1,7 |
| 13 | 108,0 | 160,0 | 52,0 | 1,5 | 99,3 | 151,4 | 52,0 | 1,5 |
| 14 | 67,3 | 130,0 | 62,7 | 1,9 | 105,0 | 157,4 | 52,4 | 1,5 |
| 15 | 114,6 | 163,5 | 48,9 | 1,4 | 151,9 | 279,9 | 128,0 | 1,8 |
| 16 | 121,1 | 232,5 | 111,4 | 1,9 | 118,0 | 146,6 | 28,6 | 1,2 |
| 17 | 141,2 | 259,9 | 118,7 | 1,8 | 91,9 | 168,9 | 77,0 | 1,8 |
| 18 | 157,5 | 262,1 | 104,6 | 1,7 | 92,6 | 110,7 | 18,1 | 1,2 |
| 19 | 134,9 | 440,3 | 305,4 | 3,3 | 100,2 | 127,0 | 26,8 | 1,3 |
| 20 | 61,8 | 306,1 | 244,3 | 5,0 | | | | |
| 21 | 85,4 | 73,4 | -12,1 | 0,9 | | | | |
| 22 | 92,3 | 113,0 | 20,8 | 1,2 | | | | |
| 23 | 112,5 | 180,0 | 67,5 | 1,6 | | | | |
| 24 | 94,2 | 172,9 | 78,7 | 1,8 | | | | |
| median | | | | 1,8 | | | | 1,7 |
| P-VALUE | | | | ≤0.001 | | | | ≤0.001 |

**Statistics**

| single spheres | DEGREES OF FREEDOM |
|---|---|
| Between groups | 3 |

| Comparison | | | Diff of Ranks | Q | P<0,05 |
|---|---|---|---|---|---|
| Wnt3a 20 ng/ml + RSPO1 100 ng/ml | vs | Control | 51,195 | 5,505 | Yes |
| Wnt3a 20 ng/ml + RSPO1 100 ng/ml | vs | RSPO1 100 ng/ml | 2,434 | 0,239 | No |
| Wnt3a 20 ng/ml + RSPO1 100 ng/ml | vs | Wnt3a 20 ng/ml | 1,059 | 0,101 | Do Not Test |
| Wnt3a 20 ng/ml | vs | Control | 50,136 | 5,842 | Yes |
| Wnt3a 20 ng/ml | vs | RSPO1 100 ng/ml | 1,375 | 0,144 | Do Not Test |
| RSPO1 100 ng/ml | vs | Control | 48,761 | 5,971 | Yes |

**Figure 2F**

| experimental groups | HuC/D+ cells counted of two technical replicates | mean values of all experiments | SD | P-VALUE |
|---|---|---|---|---|
| Control_n1 | 19287 | | | |
| Control_n2 | 15034 | | | |
| Control_n3 | 19896 | 18072 | 2649 | - |
| Wnt3a 20 ng/ml_n1 | 32279 | | | |
| Wnt3a 20 ng/ml_n2 | 31560 | | | |
| Wnt3a 20 ng/ml_n3 | 29554 | 31131 | 1412 | ≤0.001 |
| RSPO1 100 ng/ml_n1 | 30023 | | | |
| RSPO1 100 ng/ml_n2 | 26048 | | | |
| RSPO1 100 ng/ml_n3 | 27728 | 27933 | 1995 | 0.002 |
| Wnt3a 20 ng/ml + RSPO1 100 ng/ml_n1 | 27180 | | | |
| Wnt3a 20 ng/ml + RSPO1 100 ng/ml_n2 | 19891 | | | |
| Wnt3a 20 ng/ml + RSPO1 100 ng/ml_n3 | 24917 | 23996 | 3731 | 0.0023 |

**Statistics**

| HuCD | DEGREES OF FREEDOM | F-VALUE |
|---|---|---|
| Between groups | 3 | 14.102 |
| Residuals | 8 | |
| Total | 11 | |

| Comparison | | | Diff of Means | LSD | P<0,05 | Significance |
|---|---|---|---|---|---|---|
| Wnt3a 20 ng/ml | vs. | Control | 13058,667 | 4883,704 | <0,001 | Yes |
| Wnt3a 20 ng/ml | vs. | Wnt3a 20 ng/ml + RSPO1 100 ng/ml | 7135 | 4883,704 | 0,01 | Yes |
| Wnt3a 20 ng/ml | vs. | RSPO1 100 ng/ml | 3198 | 4883,704 | 0,169 | No |
| RSPO1 100 ng/ml | vs. | Control | 9860,667 | 4883,704 | 0,002 | Yes |
| RSPO1 100 ng/ml | vs. | Wnt3a 20 ng/ml + RSPO1 100 ng/ml | 3937 | 4883,704 | 0,1 | No |
| Wnt3a 20 ng/ml + RSPO1 100 ng/ml | vs. | Control | 5923,667 | 4883,704 | 0,023 | Yes |

**Figure 3C**

| experimental groups | Sphere count | relative sphere cout numbers | median sphere numbers | P-VALUE | summed up cell volume in µm³ | relative sphere volume numbers | median sphere volume | P-VALUE |
|---|---|---|---|---|---|---|---|---|
| Control_n1 | 84 | 1,0 | | | 13435504 | 1,0 | | |
| Control_n2 | 68 | 1,0 | | | 4138698 | 1,0 | | |
| Control_n3 | 246 | 1,0 | | | 22415501 | 1,0 | | |
| Control_n4 | 208 | 1,0 | | - | 41317156 | 1,0 | | - |
| Wnt3a 20 ng/ml_n1 | 225 | 2,7 | | | 27371857 | 2,0 | | |
| Wnt3a 20 ng/ml_n2 | 249 | 3,7 | | | 7012318 | 1,7 | | |
| Wnt3a 20 ng/ml_n3 | 457 | 1,9 | | | 38766352 | 1,7 | | |
| Wnt3a 20 ng/ml_n4 | 353 | 1,7 | 2,3 | 0.028 | 65665739 | 1,6 | 1,7 | 0..023 |
| RSPO1 100 ng/ml_n1 | 209 | 2,5 | | | 22676596 | 1,7 | | |
| RSPO1 100 ng/ml_n2 | 174 | 2,6 | | | 6605951 | 1,6 | | |
| RSPO1 100 ng/ml_n3 | 359 | 1,5 | | | 51715628 | 2,3 | | |
| RSPO1 100 ng/ml_n4 | 353 | 1,7 | 2,1 | 0.028 | 61639546 | 1,5 | 1,6 | 0.023 |
| Wnt3a 20 ng/ml + RSPO1 100 ng/ml_n1 | 236 | 2,8 | | | 24156979 | 1,8 | | |
| Wnt3a 20 ng/ml + RSPO1 100 ng/ml_n2 | 159 | 2,3 | | | 8083775 | 2,0 | | |
| Wnt3a 20 ng/ml + RSPO1 100 ng/ml_n3 | 385 | 1,6 | | | 40702832 | 1,8 | | |
| Wnt3a 20 ng/ml + RSPO1 100 ng/ml_n4 | 411 | 2,0 | 2,2 | 0.028 | 70689498 | 1,7 | 1,8 | 0.023 |

**Statistics**

| number and volume | DEGREES OF FREEDOM |
|---|---|
| Between groups | 3 |

| Volume | | | | | |
|---|---|---|---|---|---|
| **Comparison** | | | **Diff of Ranks** | **q** | **P<0,05** |
| Wnt3a 20 ng/ml + RSPO1 100 ng/ml | vs | Control | 39 | 4,096 | Yes |
| Wnt3a 20 ng/ml + RSPO1 100 ng/ml | vs | RSPO1 100 ng/ml | 13 | 1,803 | No |
| Wnt3a 20 ng/ml + RSPO1 100 ng/ml | vs | Wnt3a 20 ng/ml | 8 | 1,633 | Do Not Test |
| Wnt3a 20 ng/ml | vs | Control | 31 | 4,299 | Yes |
| Wnt3a 20 ng/ml | vs | RSPO1 100 ng/ml | 5 | 1,021 | Do Not Test |
| RSPO1 100 ng/ml | vs | Control | 26 | 5,307 | Yes |
| **Number** | | | | | |
| **Comparison** | | | **Diff of Ranks** | **q** | **P<0,05** |
| Wnt3a 20 ng/ml | vs | Control | 36,5 | 3,833 | Yes |
| Wnt3a 20 ng/ml | vs | RSPO1 100 ng/ml | 9 | 1,248 | No |
| Wnt3a 20 ng/ml | vs | Wnt3a 20 ng/ml + RSPO1 100 ng/ml | 4,5 | 0,919 | Do Not Test |
| Wnt3a 20 ng/ml + RSPO1 100 ng/ml | vs | Control | 32 | 4,438 | Yes |
| Wnt3a 20 ng/ml + RSPO1 100 ng/ml | vs | RSPO1 100 ng/ml | 4,5 | 0,919 | Do Not Test |
| RSPO1 100 ng/ml | vs | Control | 27,5 | 5,613 | Yes |

**Figure 3F**

| experimental groups | HuC/D+ cells counted of four technical replicates | relative values | median | P-VALUE |
|---|---|---|---|---|
| Control_n1 | 16 | 1,0 | | |
| Control_n2 | 22 | 1,0 | | |
| Control_n3 | 23 | 1,0 | | |
| Control_n4 | 118 | 1,0 | | - |
| Wnt3a 20 ng/ml_n1 | 34 | 2,1 | | |
| Wnt3a 20 ng/ml_n2 | 56 | 2,5 | | |
| Wnt3a 20 ng/ml_n3 | 51 | 2,2 | | |
| Wnt3a 20 ng/ml_n4 | 299 | 2,5 | 2,4 | 0.004 |
| RSPO1 100 ng/ml_n1 | 34 | 2,1 | | |
| RSPO1 100 ng/ml_n2 | 45 | 2,0 | | |
| RSPO1 100 ng/ml_n3 | 46 | 2,0 | | |
| RSPO1 100 ng/ml_n4 | 298 | 2,5 | 2,1 | 0.004 |
| Wnt3a 20 ng/ml + RSPO1 100 ng/ml_n1 | 53 | 3,3 | | |
| Wnt3a 20 ng/ml + RSPO1 100 ng/ml_n2 | 91 | 4,1 | | |
| Wnt3a 20 ng/ml + RSPO1 100 ng/ml_n3 | 74 | 3,2 | | |
| Wnt3a 20 ng/ml + RSPO1 100 ng/ml_n4 | 337 | 2,9 | 3,3 | 0.004 |

**Statistics**

| number and volume | DEGREES OF FREEDOM |
|---|---|
| Between groups | 3 |

| Comparison | | | Diff of Ranks | q | P<0,05 |
|---|---|---|---|---|---|
| Wnt3a 20 ng/ml + RSPO1 100 ng/ml | vs | Control | 48 | 5,041 | Yes |
| Wnt3a 20 ng/ml + RSPO1 100 ng/ml | vs | RSPO1 100 ng/ml | 29,5 | 4,091 | Yes |
| Wnt3a 20 ng/ml + RSPO1 100 ng/ml | vs | Wnt3a 20 ng/ml | 18,5 | 3,776 | Yes |
| Wnt3a 20 ng/ml | vs | Control | 29,5 | 4,091 | Yes |
| Wnt3a 20 ng/ml | vs | RSPO1 100 ng/ml | 11 | 2,245 | No |
| RSPO1 100 ng/ml | vs | Control | 18,5 | 3,776 | Yes |

**Figure5, 6 and Supplementary Figure 9**

| **Sort FZD4/LGR4** | | | | | | | | |
|---|---|---|---|---|---|---|---|---|
| Counts (of parent population) | **n1** | **n2** | **n3** | **n4** | **median** | **25% quartile** | **75% quartile** | **IQR** |
| all events | 100000 | 100000 | 100000 | 100000 | 100000 | 100000 | 100000 | 0 |
| viable cells (P1) | 73300 | 71100 | 42400 | 82300 | 72200 | 63925 | 75550 | 11625 |
| single cells FSC (P2) | 73300 | 71100 | 42400 | 82300 | 72200 | 63925 | 75550 | 11625 |
| single cells SSC (P3) | 69269 | 68327 | 41128 | 66169 | 67248 | 59909 | 68562 | 8654 |
| Fzd4+Lgr4+ (P4) | 7990 | 4783 | 1028 | 5691 | 5237 | 3844 | 6266 | 2422 |
| Fzd4+Lgr4- (P5) | 12021 | 7653 | 2180 | 2382 | 5018 | 2332 | 8745 | 6414 |
| %-parent population | **n1** | **n2** | **n3** | **n4** | **median** | **25% quartile** | **75% quartile** | **IQR** |
| all events | 100,0 | 100,0 | 100,0 | 100,0 | 100,0 | 100,0 | 100,0 | 0 |
| viable cells (P1) | 73,3 | 71,1 | 42,4 | 82,3 | 72,2 | 63,9 | 75,6 | 11,63 |
| single cells FSC (P2) | 100,0 | 100,0 | 100,0 | 100,0 | 100,0 | 100,0 | 100,0 | 0,00 |
| single cells SSC (P3) | 94,5 | 96,1 | 97,0 | 80,4 | 95,3 | 91,0 | 96,3 | 5,3 |
| Fzd4+Lgr4+ (P4) | 10,9 | 7,00 | 2,5 | 8,60 | 7,80 | 5,88 | 9,18 | 3,30 |
| Fzd4+Lgr4- (P5) | 16,4 | 11,2 | 5,3 | 3,60 | 8,25 | 4,88 | 12,5 | 7,63 |
| %-total population | **n1** | **n2** | **n3** | **n4** | **median** | **25% quartile** | **75% quartile** | **IQR** |
| all events | 100,0 | 100,0 | 100,0 | 100,0 | 100,0 | 100,0 | 100,0 | 0 |
| viable cells (P1) | 73,3 | 71,1 | 42,4 | 82,3 | 72,2 | 63,9 | 75,6 | 11,63 |
| single cells FSC (P2) | 73,3 | 71,1 | 42,4 | 82,2 | 72,2 | 63,9 | 75,5 | 11,60 |
| single cells SSC (P3) | 69,3 | 68,3 | 41,1 | 66,1 | 67,2 | 59,9 | 68,6 | 8,70 |
| Fzd4+Lgr4+ (P4) | 7,50 | 4,80 | 1,0 | 5,70 | 5,25 | 3,85 | 6,15 | 2,30 |
| Fzd4+Lgr4- (P5) | 11,4 | 7,6 | 2,2 | 2,40 | 5,00 | 2,35 | 8,55 | 6,20 |

| **Sort FZD4/LGR5** | | | | | | | | | |
|---|---|---|---|---|---|---|---|---|---|
| Counts (of parent population) | **n1** | **n2** | **n3** | **n4** | **n5** | **median** | **25% quartile** | **75% quartile** | **IQR** |
| all events | 100000 | 100000 | 100000 | 100000 | 100000 | 100000 | 100000 | 100000 | 0 |
| viable cells (P1) | 61800 | 64200 | 81900 | 69000 | 76200 | 69000 | 64200 | 76200 | 12000 |
| single cells FSC (P2) | 55929 | 58615 | 81736 | 68862 | 76200 | 68862 | 58615 | 76200 | 17585 |
| single cells SSC (P3) | 53524 | 56446 | 71683 | 63766 | 72619 | 63766 | 56446 | 71683 | 15237 |
| Fzd4+Lgr5+ (P4) | 3533 | 5475 | 4946 | 6185 | 6463 | 5475 | 4946 | 6185 | 1239 |
| Fzd4+Lgr5- (P5) | 5459 | 4403 | 3512 | 9501 | 10893 | 5459 | 4403 | 9501 | 5098 |
| %-parent population | **n1** | **n2** | **n3** | **n4** | **n5** | **median** | **25% quartile** | **75% quartile** | **IQR** |
| all events | 100 | 100 | 100 | 100,0 | 100,0 | 100,0 | 100,0 | 100,0 | 0,0 |
| viable cells (P1) | 61,8 | 64,2 | 81,9 | 69,0 | 76,2 | 69,0 | 64,2 | 76,2 | 12,0 |
| single cells FSC (P2) | 90,5 | 91,3 | 99,8 | 99,8 | 100,0 | 99,8 | 91,3 | 99,8 | 8,50 |
| single cells SSC (P3) | 95,7 | 96,3 | 87,7 | 92,6 | 95,3 | 95,3 | 92,6 | 95,7 | 3,10 |
| Fzd4+Lgr5+ (P4) | 6,60 | 9,70 | 6,90 | 9,70 | 8,90 | 8,90 | 6,90 | 9,70 | 2,80 |
| Fzd4+Lgr5- (P5) | 10,2 | 7,80 | 4,90 | 14,9 | 15,0 | 10,2 | 7,80 | 14,9 | 7,10 |
| %-total population | **n1** | **n2** | **n3** | **n4** | **n5** | **median** | **25% quartile** | **75% quartile** | **IQR** |
| all events | 100 | 100 | 100 | 100,0 | 100,0 | 100,0 | 100,0 | 100,0 | 0,0 |
| viable cells (P1) | 61,8 | 64,2 | 81,9 | 69,0 | 76,2 | 69,0 | 64,2 | 76,2 | 12,0 |
| single cells FSC (P2) | 55,9 | 58,6 | 81,7 | 68,8 | 76,1 | 68,8 | 58,6 | 76,1 | 17,5 |
| single cells SSC (P3) | 53,5 | 56,5 | 71,7 | 63,7 | 72,6 | 63,7 | 56,5 | 71,7 | 15,2 |
| Fzd4+Lgr5+ (P4) | 3,50 | 5,50 | 5,00 | 6,20 | 6,50 | 5,50 | 5,00 | 6,20 | 1,20 |
| Fzd4+Lgr5- (P5) | 5,40 | 4,40 | 3,50 | 9,50 | 10,9 | 5,40 | 4,40 | 9,50 | 5,10 |

| **Sort FZD4/LGR6** | | | | | | | | |
|---|---|---|---|---|---|---|---|---|
| Counts (of parent population) | **n1** | **n2** | **n3** | **n4** | **median** | **25% quartile** | **75% quartile** | **IQR** |
| all events | 100000 | 100000 | 100000 | 100000 | 100000 | 100000 | 100000 | 0 |
| viable cells (P1) | 80092 | 77649 | 40300 | 40500 | 59074 | 40450 | 78260 | 37810 |
| single cells FSC (P2) | 64694 | 77274 | 39534 | 40136 | 52415 | 39986 | 67839 | 27854 |
| single cells SSC (P3) | 55010 | 76148 | 37320 | 37768 | 46389 | 37656 | 60295 | 22638 |
| Fzd4+Lgr6+ (P4) | 11442 | 685 | 7725 | 4910 | 6318 | 3854 | 8654 | 4801 |
| Fzd4+Lgr6- (P5) | 2200 | 914 | 368 | 2568 | 1557 | 778 | 2292 | 1514 |
| %-parent population | **n1** | **n2** | **n3** | **n4** | **median** | **25% quartile** | **75% quartile** | **IQR** |
| all events | 100,0 | 100,0 | 100,0 | 100,0 | 100,0 | 100,0 | 100,0 | 0,0 |
| viable cells (P1) | 80,1 | 77,6 | 40,3 | 40,5 | 59,1 | 40,5 | 78,2 | 37,8 |
| single cells FSC (P2) | 80,8 | 99,5 | 98,1 | 99,1 | 98,6 | 93,8 | 99,2 | 5,43 |
| single cells SSC (P3) | 85,0 | 98,5 | 94,4 | 94,1 | 94,3 | 91,8 | 95,4 | 3,60 |
| Fzd4+Lgr6+ (P4) | 2,80 | 1,20 | 20,7 | 13,0 | 7,90 | 2,40 | 14,9 | 12,5 |
| Fzd4+Lgr6- (P5) | 4,00 | 0,90 | 9,90 | 6,80 | 5,40 | 3,23 | 7,58 | 4,35 |
| %-total population | **n1** | **n2** | **n3** | **n4** | **median** | **25% quartile** | **75% quartile** | **IQR** |
| all events | 100,0 | 100,0 | 100,0 | 100 | 100,0 | 100,0 | 100,0 | 0,0 |
| viable cells (P1) | 80,1 | 77,6 | 40,3 | 40,6 | 59,1 | 40,5 | 78,2 | 37,7 |
| single cells FSC (P2) | 64,1 | 77,3 | 39,5 | 14,4 | 51,8 | 33,2 | 67,4 | 34,2 |
| single cells SSC (P3) | 55,0 | 76,1 | 37,3 | 13,6 | 46,2 | 31,4 | 60,3 | 28,9 |
| Fzd4+Lgr6+ (P4) | 1,80 | 0,90 | 7,70 | 1,80 | 1,80 | 1,58 | 3,28 | 1,70 |
| Fzd4+Lgr6- (P5) | 2,60 | 0,70 | 3,70 | 0,90 | 1,75 | 0,85 | 2,88 | 2,03 |

**Figure 7B / Figure 7C / Figure 7D**

| LGR+DAPI+ cells | experiments | LGR4+ cells | LGR5+ cells | LGR6+ cells |
|---|---|---|---|---|
| proliferation_div5 | n1 | 1055 | 5321 | 2 |
| | n2 | 1378 | 3960 | 1 |
| | n3 | 1294 | 4106 | 4 |
| proliferation_div6 | n1 | 2136 | 6320 | 162 |
| | n2 | 2065 | 7003 | 204 |
| | n3 | 1984 | 6935 | 179 |
| proliferation_div9 | n1 | 3987 | 9874 | 280 |
| | n2 | 3753 | 11237 | 341 |
| | n3 | 3520 | 11940 | 297 |
| proliferation_div12 | n1 | 6231 | 18965 | 562 |
| | n2 | 5947 | 15678 | 678 |
| | n3 | 4201 | 17985 | 524 |
| differentiation_div6 | n1 | 6724 | 18745 | 4362 |
| | n2 | 7961 | 13564 | 3954 |
| | n3 | 5901 | 15742 | 3201 |
| differentiation_div9 | n1 | 22643 | 60641 | 32695 |
| | n2 | 20498 | 65832 | 30457 |
| | n3 | 24631 | 66147 | 29842 |
| differentiation_div12 | n1 | 40651 | 89562 | 95632 |
| | n2 | 37896 | 81073 | 100388 |
| | n3 | 37465 | 90332 | 85632 |

| LGR+DAPI+/ DAPI+ cells | experiments | %-LGR4+ cells | mean | SD | P-VALUE | %-LGR5+ cells | mean | SD | P-VALUE | %-LGR6+ cells | median | P-VALUE |
|---|---|---|---|---|---|---|---|---|---|---|---|---|
| proliferation_div5 | n1 | 0,094 | 0,115 | 0,018 | | 0,494 | 0,441 | 0,047 | | 0,000 | 0,000 | |
| | n2 | 0,129 | | | | 0,405 | | | | 0,000 | | |
| | n3 | 0,123 | | | | 0,425 | | | | 0,000 | | |
| proliferation_div6 | n1 | 0,174 | 0,170 | 0,007 | ≤0.001 | 0,567 | 0,538 | 0,028 | ≤0.001 | 0,011 | 0,013 | ≤0.05 |
| | n2 | 0,162 | | | | 0,536 | | | | 0,013 | | |
| | n3 | 0,175 | | | | 0,512 | | | | 0,014 | | |
| proliferation_div9 | n1 | 0,278 | 0,280 | 0,007 | 0.047 | 0,756 | 0,751 | 0,005 | 0.003 | 0,019 | 0,019 | ≤0.05 |
| | n2 | 0,287 | | | | 0,749 | | | | 0,020 | | |
| | n3 | 0,274 | | | | 0,746 | | | | 0,017 | | |
| proliferation_div12 | n1 | 0,309 | 0,304 | 0,026 | ≤0.001 | 0,737 | 0,736 | 0,030 | ≤0.001 | 0,027 | 0,026 | ≤0.05 |
| | n2 | 0,326 | | | | 0,765 | | | | 0,026 | | |
| | n3 | 0,276 | | | | 0,705 | | | | 0,025 | | |
| differentiation_div6 | n1 | 0,286 | 0,288 | 0,024 | ≤0.001 | 0,784 | 0,745 | 0,042 | ≤0.001 | 0,233 | 0,217 | ≤0.05 |
| | n2 | 0,313 | | | | 0,700 | | | | 0,197 | | |
| | n3 | 0,266 | | | | 0,750 | | | | 0,217 | | |
| differentiation_div9 | n1 | 0,324 | 0,313 | 0,023 | 0.047 | 0,825 | 0,836 | 0,018 | 0.003 | 0,434 | 0,414 | ≤0.05 |
| | n2 | 0,287 | | | | 0,856 | | | | 0,369 | | |
| | n3 | 0,327 | | | | 0,826 | | | | 0,414 | | |
| differentiation_div12 | n1 | 0,448 | 0,452 | 0,016 | ≤0.001 | 1,000 | 1,000 | 0,000 | ≤0.001 | 1,000 | 1,000 | ≤0.05 |
| | n2 | 0,469 | | | | 1,000 | | | | 1,000 | | |
| | n3 | 0,438 | | | | 1,000 | | | | 1,000 | | |

**Statistics**

| LGR4+DAPI / DAPI | DEGREES OF FREEDOM | F-VALUE |
|---|---|---|
| Between groups | 6 | 101 |
| Residuals | 14 | |
| Total | 20 | |

| LGR5+DAPI/DAPI | DEGREES OF FREEDOM | F-VALUE |
|---|---|---|
| Between groups | 6 | 101 |
| Residuals | 14 | |
| Total | 20 | |

| LGR6+DAPI/DAPI | DEGREES OF FREEDOM | F-VALUE |
|---|---|---|
| Between groups | 6 | 101 |
| Residuals | 14 | |
| Total | 20 | |

| LGR+DAPI / DAPI | Comparison | Diff of Means | LSD | P<0,05 | Significance |
|---|---|---|---|---|---|
| LGR4 | div5_prol vs div5_diff | - | - | - | - |
| | div6_prol vs div6_diff | 0,118 | 0,0327 | <0,001 | Yes |
| | div9_prol vs div9_diff | 0,0331 | 0,0327 | 0,047 | Yes |
| | div12_prol vs div12_diff | 0,148 | 0,0327 | <0,001 | Yes |
| LGR5 | div5_prol vs div5_diff | - | - | - | - |
| | div6_prol vs div6_diff | 0,207 | 0,0512 | <0,001 | Yes |
| | div9_prol vs div9_diff | 0,0852 | 0,0512 | 0,003 | Yes |
| | div12_prol vs div12_diff | 0,264 | 0,0512 | <0,001 | Yes |
| **LGR+DAPI / DAPI** | **Comparison** | **Diff od Ranks** | **q** | **P<0,05** | |
| LGR6 | div5_prol vs div5_diff | - | - | - | |
| | div6_prol vs div6_diff | 27 | 4,323 | Yes | |
| | div9_prol vs div9_diff | 27 | 4,323 | Yes | |
| | div12_prol vs div12_diff | 27 | 4,323 | Yes | |

**Figure 7E / Figure 7F**

| Ki67+LGR+DAPI+ cells | experiments | LGR4+ cells | LGR5+ cells |
|---|---|---|---|
| proliferation_div5 | n1 | 174 | 1046 |
| | n2 | 162 | 942 |
| | n3 | 173 | 989 |
| proliferation_div6 | n1 | 363 | 1245 |
| | n2 | 341 | 1296 |
| | n3 | 280 | 1436 |
| proliferation_div9 | n1 | 789 | 1645 |
| | n2 | 695 | 1423 |
| | n3 | 702 | 1721 |
| proliferation_div12 | n1 | 613 | 2012 |
| | n2 | 524 | 1998 |
| | n3 | 490 | 2301 |
| differentiation_div6 | n1 | 1082 | 1680 |
| | n2 | 1122 | 1461 |
| | n3 | 987 | 1946 |
| differentiation_div9 | n1 | 5408 | 10 |
| | n2 | 5266 | 19 |
| | n3 | 5387 | 32 |
| differentiation_div12 | n1 | 50 | 46 |
| | n2 | 41 | 40 |
| | n3 | 48 | 55 |

| Ki67+LGR+DAPI+/ LGR+DAPI+ cells | experiments | %-LGR4+ cells | mean | SD | P-VALUE | %-LGR5+ cells | median | 25% quartile | 75% quartile | IQR | P-VALUE |
|---|---|---|---|---|---|---|---|---|---|---|---|
| proliferation_div5 | n1 | 0,165 | 0,139 | 0,02408155 | | 0,197 | 23,788% | 21,7% | 23,9% | 2,21% | |
| | n2 | 0,118 | | | | 0,238 | | | | | |
| | n3 | 0,134 | | | | 0,241 | | | | | |
| proliferation_div6 | n1 | 0,170 | 0,159 | 0,015436067 | 0.853 | 0,197 | 19,699% | 19,1% | 20,2% | 1,10% | ≤0.05 |
| | n2 | 0,165 | | | | 0,185 | | | | | |
| | n3 | 0,141 | | | | 0,207 | | | | | |
| proliferation_div9 | n1 | 0,198 | 0,194 | 0,007819063 | 0.003 | 0,167 | 14,414% | 13,5% | 15,5% | 2,00% | ≤0.05 |
| | n2 | 0,185 | | | | 0,127 | | | | | |
| | n3 | 0,199 | | | | 0,144 | | | | | |
| proliferation_div12 | n1 | 0,098 | 0,101 | 0,014449017 | ≤0.001 | 0,106 | 12,744% | 11,7% | 12,8% | 1,09% | ≤0.05 |
| | n2 | 0,088 | | | | 0,127 | | | | | |
| | n3 | 0,117 | | | | 0,128 | | | | | |
| differentiation_div6 | n1 | 0,161 | 0,156 | 0,013737359 | 0.853 | 0,087 | 10,771% | 9,7% | 11,6% | 1,83% | ≤0.05 |
| | n2 | 0,141 | | | | 0,108 | | | | | |
| | n3 | 0,167 | | | | 0,124 | | | | | |
| differentiation_div9 | n1 | 0,239 | 0,238 | 0,019106782 | 0.003 | 0,000165 | 0,029% | 0,0% | 0,0% | 0,02% | ≤0.05 |
| | n2 | 0,257 | | | | 0,000289 | | | | | |
| | n3 | 0,219 | | | | 0,000484 | | | | | |
| differentiation_div12 | n1 | 0,001 | 0,001 | 0,000103492 | ≤0.001 | 0,001 | 0,051% | 0,1% | 0,1% | 0,01% | ≤0.05 |
| | n2 | 0,001 | | | | 0,000 | | | | | |
| | n3 | 0,001 | | | | 0,001 | | | | | |

**Statistics**

| LGR4+Ki67+DAPI/ LGR4+DAPI | DEGREES OF FREEDOM | F-VALUE |
|---|---|---|
| Between groups | 6 | 72.40 |
| Residuals | 14 | |
| Total | 20 | |

| LGR5+Ki67+DAPI/ LGR5+DAPI | DEGREES OF FREEDOM |
|---|---|
| Between groups | 6 |

| DAPI+Ki67+LGR/LGR+DAPI | Comparison | Diff of Means | LSD | P<0,05 | Significance |
|---|---|---|---|---|---|
| LGR4 | div5_prol vs div5_diff | - | - | - | - |
| | div6_prol vs div6_diff | 0,00236 | 0,0268 | 0,853 | Do Not Test |
| | div9_prol vs div9_diff | 0,044 | 0,0268 | 0,003 | Yes |
| | div12_prol vs div12_diff | 0,0998 | 0,0268 | <0,001 | Yes |
| | **Comparison** | **Diff od Ranks** | **q** | **P<0,05** | |
| LGR5 | div5_prol vs div5_diff | - | - | - | |
| | div6_prol vs div6_diff | 27 | 4,323 | Yes | |
| | div9_prol vs div9_diff | 34 | 4,389 | Yes | |
| | div12_prol vs div12_diff | 18 | 3,795 | Yes | |

**Figure 7G + Suppl. Figure 12**

| DAPI+Ki67+LGR4 | | | | | | | | |
|---|---|---|---|---|---|---|---|---|
| **Control** | DAPI+ | DAPI+Ki67+ | DAPI+Ki67- | DAPI+LGR4+ | DAPI+LGR4- | DAPI+LGR4+Ki67+ | DAPI+LGR4+KI67- | P-VALUE |
| n1 | 1126 | 241 | 885 | 82 | 1044 | 6 | 82 | |
| n2 | 779 | 213 | 566 | 39 | 740 | 3 | 39 | |
| n3 | 668 | 157 | 511 | 82 | 586 | 8 | 82 | |
| calculated percentages | | | | | | | | |
| n1 | 100% | 21% | 79% | 7% | 93% | 7% | 100% | |
| n2 | 100% | 27% | 73% | 5% | 95% | 8% | 100% | |
| n3 | 100% | 24% | 76% | 12% | 88% | 10% | 100% | |
| **RSPO1 100 ng/ml** | DAPI+ | DAPI+Ki67+ | DAPI+Ki67- | DAPI+LGR4+ | DAPI+LGR4- | DAPI+LGR4+Ki67+ | DAPI+LGR4+KI67- | |
| n1 | 1011 | 353 | 658 | 394 | 617 | 158 | 236 | |
| n2 | 1008 | 447 | 561 | 267 | 741 | 120 | 147 | |
| n3 | 1007 | 361 | 646 | 256 | 751 | 125 | 131 | |
| calculated percentages | | | | | | | | |
| n1 | 100% | 35% | 65% | 39% | 61% | 40% | 60% | ≤0.001 |
| n2 | 100% | 44% | 56% | 26% | 74% | 45% | 55% | |
| n3 | 100% | 36% | 64% | 25% | 75% | 49% | 51% | |

**Statistics**

| LGR+DAPI+ | DEGREES OF FREEDOM | DIFFERECES | T-VALUE |
|---|---|---|---|
| LGR4 | 4 | -0,221 | -4,557 |
| LGR5 | 4 | -0,103 | -1.628 |
| LGR6 | 4 | 0,00113 | 0,11 |

**Statistics**

| LGR+Ki67 | DEGREES OF FREEDOM | DIFFERECES | T-VALUE |
|---|---|---|---|
| LGR4 | 4 | -0,364 | -13,799 |
| LGR5 | 4 | -0,193 | -6,765 |

**Figure 7H + Suppl. Figure 12**

| DAPI+Ki67+LGR5 | | | | | | | | |
|---|---|---|---|---|---|---|---|---|
| **Control** | DAPI+ | DAPI+Ki67+ | DAPI+Ki67- | DAPI+LGR5+ | DAPI+LGR5- | DAPI+LGR5+Ki67+ | DAPI+LGR5+KI67- | P-VALUE |
| n1 | 1014 | 271 | 743 | 358 | 656 | 108 | 250 | |
| n2 | 851 | 275 | 576 | 363 | 488 | 113 | 250 | |
| n3 | 680 | 241 | 439 | 338 | 342 | 110 | 228 | |
| calculated percentages | | | | | | | | |
| n1 | 100% | 27% | 73% | 35% | 65% | 30% | 70% | |
| n2 | 100% | 32% | 68% | 43% | 57% | 31% | 69% | |
| n3 | 100% | 35% | 65% | 50% | 50% | 33% | 67% | |
| **RSPO1 100 ng/ml** | DAPI+ | DAPI+Ki67+ | DAPI+Ki67- | DAPI+LGR5+ | DAPI+LGR5- | DAPI+LGR5+Ki67+ | DAPI+LGR5+KI67- | |
| n1 | 1377 | 574 | 803 | 603 | 774 | 284 | 319 | |
| n2 | 741 | 344 | 397 | 444 | 297 | 249 | 195 | |
| n3 | 675 | 267 | 408 | 370 | 305 | 180 | 190 | |
| calculated percentages | | | | | | | | |
| n1 | 100% | 42% | 58% | 44% | 56% | 47% | 53% | 0.002 |
| n2 | 100% | 46% | 54% | 60% | 40% | 56% | 44% | |
| n3 | 100% | 40% | 60% | 55% | 45% | 49% | 51% | |

**Suppl. Figure 12**

| DAPI+Ki67+LGR6 | | | | | | | |
|---|---|---|---|---|---|---|---|
| **Control** | DAPI+ | DAPI+Ki67+ | DAPI+Ki67- | DAPI+LGR6+ | DAPI+LGR6- | DAPI+LGR6+Ki67+ | DAPI+LGR6+KI67- |
| n1 | 823 | 184 | 639 | 38 | 785 | 0 | 38 |
| n2 | 563 | 116 | 447 | 28 | 535 | 0 | 28 |
| n3 | 553 | 136 | 417 | 24 | 529 | 0 | 24 |
| calculated percentages | | | | | | | |
| n1 | 100% | 22% | 78% | 5% | 95% | 0% | 100% |
| n2 | 100% | 21% | 79% | 5% | 95% | 0% | 100% |
| n3 | 100% | 25% | 75% | 4% | 96% | 0% | 100% |
| **RSPO1 100 ng/ml** | DAPI+ | DAPI+Ki67+ | DAPI+Ki67- | DAPI+LGR6+ | DAPI+LGR6- | DAPI+LGR6+Ki67+ | DAPI+LGR6+KI67- |
| n1 | 1195 | 499 | 696 | 69 | 1126 | 0 | 69 |
| n2 | 826 | 338 | 488 | 21 | 805 | 0 | 21 |
| n3 | 796 | 308 | 488 | 42 | 754 | 0 | 42 |
| calculated percentages | | | | | | | |
| n1 | 100% | 42% | 58% | 6% | 94% | 0% | 100% |
| n2 | 100% | 41% | 59% | 3% | 97% | 0% | 100% |
| n3 | 100% | 39% | 61% | 5% | 95% | 0% | 100% |

**Supplementary Figure 2**

**counted positive events**

| experimental groups | DAPI | DAPI+P75+ | DAPI+P75- | DAPI+KI67+ | DAPI+KI67- | DAPI+Ki67+P75+ | P-VALUE |
|---|---|---|---|---|---|---|---|
| Control_n1 | 2316 | 668 | 1648 | 643 | 1673 | 225 | |
| Control_n2 | 1366 | 528 | 838 | 325 | 1041 | 131 | |
| Control_n3 | 1351 | 427 | 924 | 419 | 932 | 127 | |
| RSPO1 100 ng/ml_n1 | 1856 | 714 | 1142 | 631 | 1225 | 378 | |
| RSPO1 100 ng/ml_n2 | 1309 | 507 | 802 | 388 | 921 | 263 | |
| RSPO1 100 ng/ml_n3 | 1026 | 437 | 589 | 290 | 736 | 172 | 0,098 |

| **calculated percentages** | DAPI+P75+ | DAPI+P75- | DAPI+KI67+ | DAPI+KI67- | DAPI+Ki67+P75+/DAPI+P75+ |
|---|---|---|---|---|---|
| Control_n1 | 29% | 71% | 28% | 72% | 34% |
| Control_n2 | 39% | 61% | 24% | 76% | 25% |
| Control_n3 | 32% | 68% | 31% | 69% | 30% |
| mean | 33% | 67% | 28% | 72% | 29% |
| SD | 5% | 5% | 4% | 4% | 4% |

| **calculated percentages** | DAPI+P75+ | DAPI+P75- | DAPI+KI67+ | DAPI+KI67- | DAPI+Ki67+P75+/DAPI+P75+ | P-VALUE |
|---|---|---|---|---|---|---|
| RSPO1 100 ng/ml_n1 | 38% | 62% | 34% | 66% | 53% | |
| RSPO1 100 ng/ml_n2 | 39% | 61% | 30% | 70% | 52% | |
| RSPO1 100 ng/ml_n3 | 43% | 57% | 28% | 72% | 39% | |
| mean | 40% | 60% | 31% | 69% | 48% | |
| SD | 2% | 2% | 3% | 3% | 8% | 0,023 |

**Statistics**

| DAPI+P75+/DAPI+ | **DEGREES OF FREEDOM** | **DIFFERECES** | **T-VALUE** |
|---|---|---|---|
| | 4 | -0,069 | -2,149 |

| DAPI+P75+KI67/DAPI+P75+ | **DEGREES OF FREEDOM** | **DIFFERECES** | **T-VALUE** |
|---|---|---|---|
| | 4 | -0,194 | -3,571 |

**Supplementary Figure 3**

**Sort tdTomato**
**postnatal day 60**
**small intestine**

| Counts (of parent population) | n1 | n2 | n3 | n4 | n5 | n6 | n7 | n8 | n9 | n10 | n11 | n12 | n13 | median | 25% quartile | 75% quartile | IQR |
|---|---|---|---|---|---|---|---|---|---|---|---|---|---|---|---|---|---|
| all events | 100000 | 100000 | 100000 | 100000 | 100000 | 100000 | 100000 | 100000 | 100000 | 100000 | 100000 | 100000 | 100000 | 100000 | 100000 | 100000 | 0 |
| viable cells (P1) | 65600 | 62400 | 67200 | 72100 | 62400 | 59100 | 56500 | 47200 | 51700 | 44400 | 31600 | 45400 | 23900 | 56500 | 45400 | 62400 | 17000 |
| single cells FSC (P2) | 35949 | 45178 | 40387 | 44414 | 41558 | 40661 | 34409 | 28273 | 40895 | 27439 | 20824 | 43312 | 17997 | 40387 | 28273 | 41558 | 13286 |
| single cells SSC (P3) | 34906 | 42467 | 29523 | 38462 | 36862 | 36066 | 31965 | 24597 | 35783 | 23214 | 13640 | 33350 | 13570 | 33350 | 24597 | 36066 | 11469 |
| tdTomato - (P4) | 28588 | 33804 | 22497 | 33577 | 30817 | 26545 | 17421 | 21228 | 26873 | 17224 | 11062 | 27680 | 8196 | 26545 | 17421 | 28588 | 11167 |
| tdTomato + (P5) | 3875 | 5436 | 3868 | 2885 | 4645 | 4977 | 3676 | 2238 | 8051 | 4875 | 2155 | 4469 | 4722 | 4469 | 3676 | 4875 | 1199 |

| %-parent population | n1 | n2 | n3 | n4 | n5 | n6 | n7 | n8 | n9 | n10 | n11 | n12 | n13 | median | 25% quartile | 75% quartile | IQR |
|---|---|---|---|---|---|---|---|---|---|---|---|---|---|---|---|---|---|
| all events | 100,0 | 100,0 | 100,0 | 100,0 | 100,0 | 100,0 | 100,0 | 100,0 | 100,0 | 100,0 | 100,0 | 100,0 | 100,0 | 100,0 | 100,0 | 100,0 | 0 |
| viable cells (P1) | 65,6 | 62,4 | 67,2 | 72,1 | 62,4 | 59,1 | 56,5 | 47,2 | 51,7 | 44,4 | 31,6 | 45,4 | 23,9 | 56,5 | 45,4 | 62,4 | 17,00 |
| single cells FSC (P2) | 54,8 | 72,4 | 60,1 | 61,6 | 66,6 | 68,8 | 60,9 | 59,9 | 79,1 | 61,8 | 65,9 | 95,4 | 75,3 | 65,9 | 60,9 | 72,4 | 11,50 |
| single cells SSC (P3) | 97,1 | 94,0 | 73,1 | 86,6 | 88,7 | 88,7 | 92,9 | 87,0 | 87,5 | 84,6 | 65,5 | 77,0 | 75,4 | 87,0 | 77,0 | 88,7 | 11,7 |
| tdTomato - (P4) | 81,9 | 79,6 | 76,2 | 87,3 | 83,6 | 73,6 | 54,5 | 86,3 | 75,1 | 74,2 | 81,1 | 83,0 | 60,4 | 79,6 | 74,2 | 83,0 | 8,80 |
| tdTomato + (P5) | 11,1 | 12,8 | 13,1 | 7,50 | 12,6 | 13,8 | 11,5 | 9,10 | 22,5 | 21,0 | 15,8 | 13,4 | 34,8 | 13,1 | 11,5 | 15,8 | 4,30 |

| %-total population | n1 | n2 | n3 | n4 | n5 | n6 | n7 | n8 | n9 | n10 | n11 | n12 | n13 | median | 25% quartile | 75% quartile | IQR |
|---|---|---|---|---|---|---|---|---|---|---|---|---|---|---|---|---|---|
| all events | 100,0 | 100,0 | 100,0 | 100,0 | 100,0 | 100,0 | 100,0 | 100,0 | 100,0 | 100,0 | 100,0 | 100,0 | 100,0 | 100,0 | 100,0 | 100,0 | 0 |
| viable cells (P1) | 65,6 | 62,4 | 67,2 | 72,1 | 62,4 | 59,1 | 56,5 | 47,2 | 51,7 | 44,4 | 31,8 | 45,4 | 23,9 | 56,5 | 45,4 | 62,4 | 17,00 |
| single cells FSC (P2) | 35,9 | 45,2 | 40,4 | 44,4 | 66,6 | 40,6 | 34,3 | 27,8 | 40,4 | 27,5 | 17,9 | 39,6 | 19,0 | 39,6 | 27,8 | 40,6 | 12,80 |
| single cells SSC (P3) | 34,9 | 42,5 | 29,5 | 38,4 | 88,7 | 36,0 | 28,5 | 24,2 | 35,3 | 23,2 | 11,6 | 30,5 | 13,6 | 30,5 | 24,2 | 36,0 | 11,80 |
| tdTomato - (P4) | 28,6 | 33,8 | 22,5 | 33,6 | 83,6 | 27,5 | 18,4 | 20,9 | 20,5 | 17,2 | 9,40 | 25,30 | 8,20 | 22,5 | 18,4 | 28,6 | 10,20 |
| tdTomato + (P5) | 3,9 | 5,4 | 3,9 | 2,90 | 12,6 | 5,00 | 3,30 | 2,20 | 8,00 | 5,10 | 1,80 | 4,10 | 4,70 | 4,1 | 3,3 | 5,1 | 1,80 |

**Sort tdTomato**
**postnatal day 60**
**large intestine**

| Counts (of parent population) | n1 | n2 | n3 | n4 | n5 | n6 | n7 | n8 | n9 | n10 | n11 | n12 | n13 | median | 25% quartile | 75% quartile | IQR |
|---|---|---|---|---|---|---|---|---|---|---|---|---|---|---|---|---|---|
| all events | 100000 | 100000 | 100000 | 100000 | 100000 | 100000 | 100000 | 100000 | 100000 | 100000 | 100000 | 100000 | 100000 | 100000 | 100000 | 100000 | 0 |
| viable cells (P1) | 59700 | 60600 | 68700 | 66600 | 64300 | 62800 | 67400 | 52600 | 51700 | 48000 | 39700 | 49600 | 29600 | 59700 | 49600 | 64300 | 14700 |
| single cells FSC (P2) | 33074 | 39390 | 47060 | 44888 | 43981 | 44400 | 44888 | 34137 | 40895 | 18480 | 29854 | 38192 | 15984 | 39390 | 33074 | 44400 | 11326 |
| single cells SSC (P3) | 31321 | 34663 | 35577 | 38604 | 38923 | 38983 | 44754 | 29392 | 35783 | 15024 | 21973 | 30286 | 11205 | 34663 | 29392 | 38604 | 9212 |
| tdTomato - (P4) | 21204 | 26517 | 25722 | 31076 | 30399 | 26041 | 30522 | 22250 | 26873 | 7993 | 17117 | 15809 | 3776 | 25722 | 17117 | 26873 | 9756 |
| tdTomato + (P5) | 9271 | 7141 | 8147 | 6447 | 7668 | 11695 | 8951 | 6055 | 8051 | 6505 | 4482 | 13811 | 6902 | 7668 | 6505 | 8951 | 2445 |

| %-parent population | n1 | n2 | n3 | n4 | n5 | n6 | n7 | n8 | n9 | n10 | n11 | n12 | n13 | median | 25% quartile | 75% quartile | IQR |
|---|---|---|---|---|---|---|---|---|---|---|---|---|---|---|---|---|---|
| all events | 100,0 | 100,0 | 100,0 | 100,0 | 100,0 | 100,0 | 100,0 | 100,0 | 100,0 | 100,0 | 100,0 | 100,0 | 100,0 | 100,0 | 100,0 | 100,0 | 0 |
| viable cells (P1) | 59,7 | 60,6 | 68,7 | 66,6 | 64,3 | 62,8 | 67,4 | 52,6 | 51,7 | 48,0 | 39,7 | 49,6 | 29,6 | 59,7 | 49,6 | 64,3 | 14,70 |
| single cells FSC (P2) | 55,4 | 65,0 | 68,5 | 67,4 | 68,4 | 70,7 | 66,6 | 64,9 | 79,1 | 38,5 | 75,2 | 77,0 | 54,0 | 67,4 | 64,9 | 70,7 | 5,80 |
| single cells SSC (P3) | 94,7 | 88,0 | 75,6 | 86,0 | 88,5 | 87,8 | 99,7 | 86,1 | 87,5 | 81,3 | 73,6 | 79,3 | 70,1 | 86,1 | 79,3 | 88,0 | 8,7 |
| tdTomato - (P4) | 67,7 | 76,5 | 72,3 | 80,5 | 78,1 | 66,8 | 68,2 | 75,7 | 75,1 | 53,2 | 77,9 | 52,2 | 33,7 | 72,3 | 66,8 | 76,5 | 9,70 |
| tdTomato + (P5) | 29,6 | 20,6 | 22,9 | 16,70 | 19,7 | 30,0 | 20,0 | 20,6 | 22,5 | 43,3 | 20,4 | 45,6 | 61,6 | 22,5 | 20,4 | 30,0 | 9,60 |

| %-total population | n1 | n2 | n3 | n4 | n5 | n6 | n7 | n8 | n9 | n10 | n11 | n12 | n13 | median | 25% quartile | 75% quartile | IQR |
|---|---|---|---|---|---|---|---|---|---|---|---|---|---|---|---|---|---|
| all events | 100,0 | 100,0 | 100,0 | 100,0 | 100,0 | 100,0 | 100,0 | 100,0 | 100,0 | 100,0 | 100,0 | 100,0 | 100,0 | 100,0 | 100,0 | 100,0 | 0 |
| viable cells (P1) | 59,7 | 60,6 | 68,7 | 66,6 | 64,3 | 62,8 | 67,4 | 52,8 | 48,2 | 49,0 | 39,7 | 49,8 | 29,6 | 59,7 | 49,0 | 64,3 | 15,30 |
| single cells FSC (P2) | 33,0 | 39,4 | 47,1 | 44,9 | 44,0 | 44,4 | 44,9 | 34,3 | 35,7 | 19,9 | 29,8 | 39,3 | 16,0 | 39,3 | 33,0 | 44,4 | 11,40 |
| single cells SSC (P3) | 31,3 | 34,7 | 35,6 | 38,6 | 38,9 | 39,0 | 40,2 | 29,5 | 31,8 | 15,4 | 22,0 | 30,4 | 11,2 | 31,8 | 29,5 | 38,6 | 9,10 |
| tdTomato - (P4) | 21,2 | 20,5 | 25,7 | 31,1 | 30,4 | 26,0 | 27,5 | 22,3 | 20,7 | 8,20 | 17,1 | 15,9 | 3,80 | 21,2 | 17,1 | 26,0 | 8,90 |
| tdTomato + (P5) | 9,3 | 7,1 | 8,1 | 6,40 | 7,70 | 11,70 | 8,00 | 6,10 | 10,4 | 6,60 | 4,50 | 13,9 | 6,90 | 7,7 | 6,6 | 9,3 | 2,70 |

**Suppl. Fig 4B**

| | Control | | | | R-Spondin1 100ng/ml | | | | Wnt3a 20ng/ml | | | | Combination | | | |
|---|---|---|---|---|---|---|---|---|---|---|---|---|---|---|---|---|
| spheres evaluated | 7 div | 14 div | diameter difference between 7 div and 14 div in µm | relative sphere diameter change | 7 div | 14 div | diameter difference between 7 div and 14 div in µm | relative sphere diameter change | 7 div | 14 div | diameter difference between 7 div and 14 div in µm | relative sphere diameter change | 7 div | 14 div | diameter difference between 7 div and 14 div in µm | relative sphere diameter change |
| 1 | 59,0 | 56,9 | -2,1 | 1,0 | 63,3 | 89,8 | 26,5 | 1,4 | 109,8 | 165,0 | 55,2 | 1,5 | 83,4 | 103,7 | 20,3 | 1,2 |
| 2 | 90,9 | 119,3 | 28,4 | 1,3 | 71,2 | 100,1 | 28,9 | 1,4 | 85,4 | 126,7 | 41,2 | 1,5 | 77,8 | 107,0 | 29,1 | 1,4 |
| 3 | 78,2 | 94,8 | 16,7 | 1,2 | 100,2 | 149,3 | 49,1 | 1,5 | 93,5 | 134,8 | 41,3 | 1,4 | 71,9 | 103,2 | 31,3 | 1,4 |
| 4 | 54,2 | 60,2 | 6,1 | 1,1 | 56,5 | 113,3 | 56,7 | 2,0 | 118,5 | 169,5 | 51,0 | 1,4 | 65,5 | 111,9 | 46,4 | 1,7 |
| 5 | 51,0 | 21,8 | -29,2 | 0,4 | 117,6 | 180,5 | 62,9 | 1,5 | 116,4 | 229,1 | 112,8 | 2,0 | 64,2 | 101,3 | 37,1 | 1,6 |
| 6 | 75,5 | 95,6 | 20,1 | 1,3 | 87,5 | 184,7 | 97,3 | 2,1 | 104,2 | 228,6 | 124,5 | 2,2 | 72,4 | 91,1 | 18,7 | 1,3 |
| 7 | 69,0 | 27,4 | -41,5 | 0,4 | 76,3 | 110,2 | 33,9 | 1,4 | 144,9 | 381,8 | 236,9 | 2,6 | 23,6 | 176,5 | 152,8 | 7,5 |
| 8 | 72,2 | 87,7 | 15,5 | 1,2 | 64,0 | 167,0 | 102,9 | 2,6 | 46,2 | 126,5 | 80,2 | 2,7 | 113,8 | 223,9 | 110,1 | 2,0 |
| 9 | 74,9 | 76,4 | 1,4 | 1,0 | 108,5 | 179,1 | 70,6 | 1,7 | 87,2 | 167,3 | 80,1 | 1,9 | 66,5 | 125,6 | 59,1 | 1,9 |
| 10 | 66,3 | 103,3 | 37,1 | 1,6 | 76,6 | 118,8 | 42,2 | 1,6 | 69,7 | 157,8 | 88,1 | 2,3 | 103,8 | 136,5 | 32,7 | 1,3 |
| 11 | 86,7 | 92,3 | 5,6 | 1,1 | 72,6 | 172,2 | 99,7 | 2,4 | 131,4 | 330,4 | 198,9 | 2,5 | 73,8 | 184,5 | 110,7 | 2,5 |
| 12 | 95,4 | 97,1 | 1,7 | 1,0 | 51,3 | 103,5 | 52,2 | 2,0 | 92,7 | 135,6 | 42,9 | 1,5 | 89,4 | 127,5 | 38,1 | 1,4 |
| 13 | 46,9 | 42,8 | -4,1 | 0,9 | 85,0 | 165,6 | 80,6 | 1,9 | 58,2 | 158,8 | 100,6 | 2,7 | 100,6 | 202,8 | 102,1 | 2,0 |
| 14 | 65,3 | 93,3 | 28,0 | 1,4 | 75,7 | 119,2 | 43,5 | 1,6 | 78,9 | 127,0 | 48,0 | 1,6 | 73,4 | 113,7 | 40,3 | 1,5 |
| 15 | 90,9 | 88,6 | -2,4 | 1,0 | 58,1 | 142,9 | 84,8 | 2,5 | 108,2 | 301,4 | 193,2 | 2,8 | 80,4 | 102,0 | 21,5 | 1,3 |
| 16 | 67,0 | 67,4 | 0,5 | 1,0 | 84,9 | 194,5 | 109,5 | 2,3 | 42,7 | 154,5 | 111,8 | 3,6 | 57,1 | 142,9 | 85,8 | 2,5 |
| 17 | 62,2 | 71,9 | 9,7 | 1,2 | 71,8 | 139,4 | 67,6 | 1,9 | | | | | | | | |
| 18 | 63,7 | 72,7 | 9,0 | 1,1 | 129,7 | 266,0 | 136,3 | 2,1 | | | | | | | | |
| 19 | 68,8 | 69,9 | 1,1 | 1,0 | 50,4 | 124,4 | 74,0 | 2,5 | | | | | | | | |
| 20 | 64,9 | 79,1 | 14,2 | 1,2 | 73,5 | 76,3 | 2,7 | 1,0 | | | | | | | | |
| 21 | 97,6 | 126,1 | 28,5 | 1,3 | | | | | | | | | | | | |
| 22 | 80,6 | 87,1 | 6,5 | 1,1 | | | | | | | | | | | | |
| 23 | 81,6 | 39,4 | -42,2 | 0,5 | | | | | | | | | | | | |
| 24 | 50,3 | 19,4 | -31,0 | 0,4 | | | | | | | | | | | | |
| 25 | 42,6 | 22,1 | -20,5 | 0,5 | | | | | | | | | | | | |
| 26 | 83,4 | 85,6 | 2,1 | 1,0 | | | | | | | | | | | | |
| 27 | 63,2 | 69,4 | 6,2 | 1,1 | | | | | | | | | | | | |
| 28 | 38,2 | 22,5 | -15,7 | 0,6 | | | | | | | | | | | | |
| 29 | 83,2 | 79,3 | -3,8 | 1,0 | | | | | | | | | | | | |
| 30 | 62,4 | 60,1 | -2,3 | 1,0 | | | | | | | | | | | | |
| 31 | 61,2 | 70,2 | 9,0 | 1,1 | | | | | | | | | | | | |
| 32 | 39,3 | 20,4 | -19,0 | 0,5 | | | | | | | | | | | | |
| P-VALUE | | | | | | | | ≤ 0.001 | | | | ≤ 0.001 | | | | ≤ 0.001 |

**Statistics**

| single spheres | DEGREES OF FREEDOM |
|---|---|
| Between groups | 3 |

| Comparison | | | Diff of Ranks | Q | P<0,05 |
|---|---|---|---|---|---|
| Wnt3a 20 ng/ml | vs | Control | 44,844 | 6,15 | Yes |
| Wnt3a 20 ng/ml | vs | 3a 20 ng/ml + RSPO1 100 r | 12,125 | 1,44 | No |
| Wnt3a 20 ng/ml | vs | RSPO1 100 ng/ml | 4,882 | 0,597 | Do Not Test |
| RSPO1 100 ng/ml | vs | Control | 39,962 | 5,695 | Yes |
| RSPO1 100 ng/ml | vs | 3a 20 ng/ml + RSPO1 100 r | 7,243 | 0,885 | Do Not Test |
| Wnt3a 20 ng/ml + RSPO1 100 ng/ml | vs | Control | 32,719 | 4,487 | Yes |

**Suppl. Fig 4D**

| experimental groups | HuC/D+ cells counted of one technical replicates | mean values of all experiments | SD | P-VALUE |
|---|---|---|---|---|
| Control_n1 | 8407 | | | |
| Control_n2 | 7864 | | | |
| Control_n3 | 5467 | 7246 | 1564 | - |
| Wnt3a 20 ng/ml_n1 | 11345 | | | |
| Wnt3a 20 ng/ml_n2 | 11028 | | | |
| Wnt3a 20 ng/ml_n3 | 9255 | 10543 | 1126 | ≤ 0.001 |
| RSPO1 100 ng/ml_n1 | 12418 | | | |
| RSPO1 100 ng/ml_n2 | 10237 | | | |
| RSPO1 100 ng/ml_n3 | 9876 | 10844 | 1375 | 0.002 |
| Wnt3a 20 ng/ml + RSPO1 100 ng/ml_n1 | 11104 | | | |
| Wnt3a 20 ng/ml + RSPO1 100 ng/ml_n2 | 10205 | | | |
| Wnt3a 20 ng/ml + RSPO1 100 ng/ml_n3 | 9647 | 10319 | 735 | 0.0023 |

**Statistics**

| single spheres | DEGREES OF FREEDOM |
|---|---|
| Between groups | 3 |

| Comparison | | | Diff of Means | P | P<0,05 | Significance |
|---|---|---|---|---|---|---|
| RSPO1 100 ng/ml | vs. | Control | 3597,667 | 2334,254 | 0,007 | Yes |
| RSPO1 100 ng/ml | vs. | Wnt3a 20 ng/ml + RSPO1 100 ng/ml | 525 | 2334,254 | 0,618 | No |
| RSPO1 100 ng/ml | vs. | Wnt3a 20 ng/ml | 301 | 2334,254 | 0,774 | Do Not Test |
| Wnt3a 20 ng/ml | vs. | Control | 3296,667 | 2334,254 | 0,012 | Yes |
| Wnt3a 20 ng/ml | vs. | Wnt3a 20 ng/ml + RSPO1 100 ng/ml | 224 | 2334,254 | 0,83 | Do Not Test |
| Wnt3a 20 ng/ml + RSPO1 100 ng/ml | vs. | Control | 3072,667 | 2334,254 | 0,016 | Yes |

**Supplementary Figure 11**

| Sort tdTomato pstnatal day 0_small intestine | | | | | | | |
|---|---|---|---|---|---|---|---|
| Counts (of parent pop | **n1** | **n2** | **n3** | **median** | **25% quartile** | **75% quartile** | **IQR** |
| all events | 100000 | 100000 | 100000 | 100000 | 100000 | 100000 | 0 |
| viable cells (P1) | 78002 | 78960 | 69300 | 78002 | 73651 | 78481 | 4830 |
| single cells FSC (P2) | 76284 | 77578 | 66251 | 76284 | 71268 | 76931 | 5664 |
| single cells SSC (P3) | 75063 | 77035 | 64793 | 75063 | 69928 | 76049 | 6121 |
| tdTomato - (P4) | 59675 | 57083 | 51899 | 57083 | 54491 | 58379 | 3888 |
| tdTomato + (P5) | 13961 | 18180 | 10691 | 13961 | 12326 | 16071 | 3745 |
| %-parent population | **n1** | **n2** | **n3** | **median** | **25% quartile** | **75% quartile** | **IQR** |
| all events | 100,0 | 100,0 | 100,0 | 100,0 | 100,0 | 100,0 | 0 |
| viable cells (P1) | 78,0 | 79,0 | 69,3 | 78,0 | 73,7 | 78,5 | 4,85 |
| single cells FSC (P2) | 97,8 | 98,2 | 95,6 | 97,8 | 96,7 | 98,0 | 1,30 |
| single cells SSC (P3) | 98,4 | 99,3 | 97,8 | 98,4 | 98,1 | 98,9 | 0,8 |
| tdTomato - (P4) | 79,5 | 74,1 | 80,1 | 79,5 | 76,8 | 79,8 | 3,00 |
| tdTomato + (P5) | 18,6 | 23,6 | 16,5 | 18,6 | 17,6 | 21,1 | 3,55 |
| %-total population | **n1** | **n2** | **n3** | **median** | **25% quartile** | **75% quartile** | **IQR** |
| all events | 100,0 | 100,0 | 100,0 | 100,0 | 100,0 | 100,0 | 0 |
| viable cells (P1) | 78,0 | 79,0 | 69,3 | 78,0 | 73,7 | 78,5 | 4,85 |
| single cells FSC (P2) | 76,3 | 77,5 | 66,3 | 76,3 | 71,3 | 76,9 | 5,60 |
| single cells SSC (P3) | 75,1 | 77,0 | 64,8 | 75,1 | 70,0 | 76,1 | 6,10 |
| tdTomato - (P4) | 59,7 | 57,0 | 51,9 | 57,0 | 54,5 | 58,4 | 3,90 |
| tdTomato + (P5) | 13,9 | 18,2 | 10,7 | 13,9 | 12,3 | 16,1 | 3,75 |
